# HERMES: Holographic Equivariant neuRal network model for Mutational Effect and Stability prediction

**DOI:** 10.1101/2024.07.09.602403

**Authors:** Gian Marco Visani, William Galvin, Zac Jones, Michael N. Pun, Eric Daniel, Kevin Borisiak, Utheri Wagura, Armita Nourmohammad

## Abstract

Accurately predicting how amino acid substitutions alter protein function is a central challenge in biology, with applications from interpreting disease variants to designing vaccines and therapeutics. We introduce HERMES, a family of fast, structure-based models that predict mutational effects from the local atomic environment around each residue. Pre-trained on masked amino acid prediction, HERMES shows strong zero-shot performance for predicting changes in thermodynamic stability and protein-protein binding affinity. Analyzing its predictions, we uncover a pre-training bias toward size-conserving substitutions, which we reduce through an amortized fine-tuning strategy that incorporates packing flexibility. When fine-tuned on experimental data, HERMES matches state-of-the-art stability predictors without costly data augmentation. HERMES also identifies antigen-stabilizing mutations across multiple viral envelope proteins, enabling an efficient pipeline for designing mutation libraries for vaccine development. Together, this work establishes HERMES as a fast and practical structure-based framework for mutation screening and offers insight into the mechanisms underlying its predictions.

## Introduction

Understanding how amino acid substitutions affect protein function is fundamental to biological discovery and engineering, with applications spanning disease variant identification [1, 2], enzyme engineering [3, 4], viral escape prediction [5–7], antibody engineering [8], and vaccine antigen stabilization [9–13].

Mutational effects on thermodynamic stability and binding affinity are especially well-studied, as stability is typically prerequisite for function [14] and most biological processes are mediated by binding. These can be measured via denaturation assays [15], surface plasmon resonance [16], or deep mutational scanning [17–19], but such experiments remain laborious despite recent throughput gains [20].

Computational approaches offer an alternative. Molecular dynamics can accurately capture short-time (nanosecond) responses but struggles with the larger conformational changes mutations often induce [21]. Physics-based energy functions such as FoldX [22] and Rosetta [23] remain widely used for mutation-induced stability changes [10], yet are often slow and inaccurate [2]. More recently, machine learning models that predict amino acid propensities at a residue from surrounding sequence [24, 25] or structural context [2, 26–30] have been found to approximate mutational effects across phenotypes.

Recent work improves these pre-trained baselines by modifying the pre-training objective [31, 32] and by fine-tuning for phenotype-specific effects [2, 27, 30, 33], often targeting thermodynamic stability for structure-based models [2, 27, 30]. Notable examples include RaSP [2], a 3D CNN fine-tuned on Rosetta-computed ΔΔ*G* values [34]; Stability-Oracle [27], a graph transformer fine-tuned on experimental stability from the Megascale dataset [20]; and ThermoMPNN [30], which fine-tunes the inverse-folding model ProteinMPNN [35] on a different Megascale subset.

Here, we present HERMES, a family of structure-based models built on our H-CNN architecture [26]. Like H-CNN, HERMES uses a 3D rotationally equivariant, all-atom CNN, but adds implementation improvements yielding a ~ 2.75 *×* speedup and an adaptable architecture for fine-tuning to arbitrary phenotypes. We pre-train HERMES on an inverse-folding objective (predicting a residue’s amino acid from its surrounding local atomic neighborhood), then optionally fine-tune for specific mutational effects. The family comprises three pre-trained variants differing in how they treat structural context: HERMES-*fixed* holds the local environment static; HERMES-*relaxed* allows local relaxation through explicit (but slow) side-chain repacking; and HERMES-*amortized* implicitly encodes substitution-dependent flexibility through amortization, at the speed of HERMES-*fixed*.

We demonstrate the utility of HERMES in both zero-shot and fine-tuned settings for predicting mutational effects on stability and binding. As a potential application, we demonstrate HERMES’ ability to identify mutations in viral envelope proteins that stabilize their metastable pre-fusion conformation; this is a key objective in vaccine design because it preserves the epitopes required to elicit protective neutralizing antibodies [9]. This suggests that HERMES can inform computational mutation library design, enabling more efficient and targeted experimental screening. Beyond these applications, we also analyze the physicochemical reasoning underlying our models, providing mechanistic insight into observed prediction biases. Together, these results establish HERMES as a practical structure-based framework for fast screening of mutations that stabilize proteins and protein complexes via local packing.

Our code is open source at https://github.com/StatPhysBio/hermes/tree/main, letting users run the models presented here and fine-tune HERMES on their own data.

## Results

### HERMES architecture and pre-training on masked amino acid classification

HERMES is a 3D, rotationally equivariant convolutional neural network that predicts the propensity of the 20 different canonical amino acids at a masked (removed) focal residue, given its local atomic neighborhood within the protein structure (Fig. 1A). Building on our prior atomistic model H-CNN [26], HERMES achieves faster inference and supports task-specific fine-tuning (see Methods and refs [36, 37] for details).

**Figure 1.**
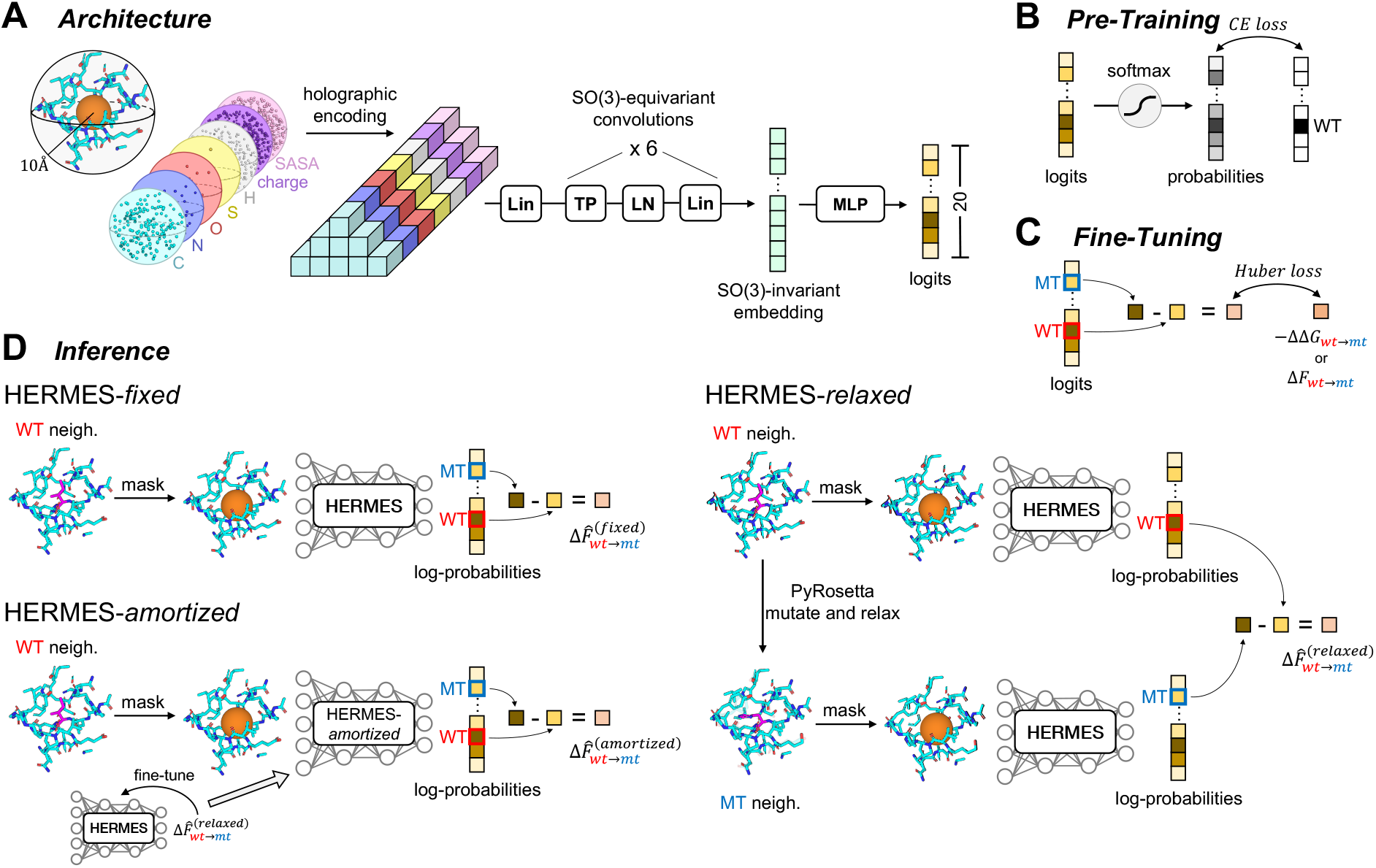
Overview of HERMES. **(A)** Model architecture: HERMES takes as input an all-atom structural neighborhood (10 Å radius) around a masked focal residue. Each atom is represented by its 3D coordinates, element type, partial charge, and solvent-accessible surface area (SASA). The neighborhood is projected onto a Zernike Fourier basis (spherical hologram) and processed by rotation (SO(3)) equivariant layers to produce a rotation-invariant embedding, which is mapped to 20 amino acid-specific logits (Methods).**(B)** Pre-training: models are trained to predict the identity of the masked focal residue from its surrounding atomic neighborhood; logits in (A) are converted to amino acid propensities (probabilities) via a softmax. **(C)** Fine-tuning for mutational effects: model weights are optimized to regress the logit difference between mutant and wild-type amino acids to the corresponding experimental mutational effect. **(D)** Inference protocols. HERMES-*fixed* scores a substitution as the difference between the mutant and wild-type amino acids logits from a single forward pass, conditioned on the masked wild-type neighborhood *X*_wt_. HERMES-*relaxed* conditions the mutant term on an approximate mutant neighborhood 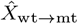 generated in-silico by introducing the mutation on the wild-type structure and locally relaxing the structure with Rosetta [34]. HERMES-*amortized* distills the relaxed protocol by fine-tuning on HERMES-*relaxed* predictions, enabling fast fixed-style inference while retaining relaxation-aware behavior.

We pre-train HERMES to recover the identity of the masked residues on ProteinNet’s CASP12 set, filtered at 30% sequence identity [38] (Fig. 1B). Each model is an ensemble of 10 independently trained networks (3.5M parameters each). To improve robustness for zero-shot applications, we additionally train models with 0.50 Å coordinate noise. Pre-training performance is summarized in Table S1.

### Zero-shot prediction of mutational effects with pre-trained HERMES

The pre-trained HERMES model outputs *P* (*aa*|*X*_*i,aa*_), the probability of assigning amino acid *aa* at residue *i*, given the surrounding atomic environment (neighborhood) *X*_*i,aa*_ in the structure. Crucially, *X*_*i,aa*_ depends on the *aa*: the neighborhood geometry has the fingerprint of the masked amino acid during pre-training.

Conditional models are widely used for zero-shot prediction of mutational effects on protein function (e.g., [24–26]). We approximate the effect of wild-type to mutant (wt → mt) substitution at residue *i* by the log-likelihood ratio between the wild-type and the mutant amino acids, conditioned on the respective local atomic neighborhoods *X*_*i*,wt_ and *X*_*i*,mt_ (omitting the residue index *i* for clarity):

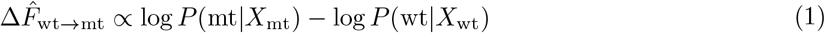

The 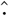 (hat) indicates the model-predicted mutational effects, as opposed to experimental measurements (no hat).

When a mutant structure is unavailable (and thus, the neighborhood *X*_mt_), the wild-type neighborhood can be relaxed *in silico* to accommodate the mutation (e.g., by Rosetta [34]), though this procedure is computationally expensive. A common alternative is to score all mutations using a single (typically wild-type) structure [26, 27, 35]. Here, we consider both of these protocols: **HERMES-*fixed*** evaluates mutational effects using the wild-type structure only (for both mt and wt propensities), while **HERMES-*relaxed*** evaluates the propensity of the mutant amino acid in the Rosetta-relaxed mt neighborhood 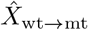, starting from the available wt structure (Fig. 1D). Note that we only relax side-chain atoms as we found that including backbone atoms substantially increased runtime at no benefit to performance; see Methods and Fig. S1 for details. The mutational effect predictions from these two approaches follow,

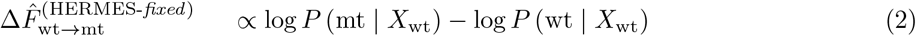

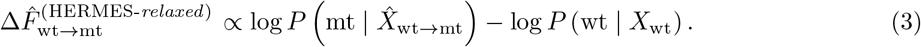

### Fine-tuning HERMES to predict protein function

We can leverage the knowledge of pre-trained HERMES, and further fine-tune the models in a supervised way to predict mutational effects on specified functions. Unlike approaches that train a separate regression head [2, 27, 30], we update the model end-to-end so that its *predicted scores* themselves align with measurements. Specifically, we make the predicted HERMES-*fixed* 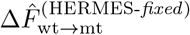 (eq. 3) regress over the experimentally measured mutational effects Δ*F*_wt→mt_ by minimizing a robust Huber loss,

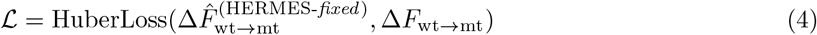

which stabilizes training under outliers while calibrating predictions to the function of interest; see Methods and Fig. 1C for details.

### Efficient encoding of side-chain flexibility with HERMES-*amortized*

Because the side-chain relaxation step in HERMES-*relaxed* is computationally costly (~ 66 times slower than HERMES-*fixed* on a single CPU / A40 GPU, Table S2), we distill HERMES-*relaxed* predictions into the model via a fine-tuning procedure. Concretely, we fine-tune the model so that the mutational-effect predictions produced with the fast HERMES-*fixed* protocol regress to the corresponding HERMES-*relaxed* predictions via Eq. 4 (Fig. 1D). We perform this fine-tuning on a small subset of neighborhoods extracted from the pre-training proteins (~ 15k neighborhoods, 0.5% of the total). The resulting amortized model HERMES-*amortized* runs at HERMES-*fixed* speed (effectively the same protocol, just different model weights) yet closely matches HERMES-*relaxed* performance (Fig. S2). We note that because our HERMES-*relaxed* protocol performs side-chain-only relaxation, HERMES-*amortized* is trained to reflect only this form of relaxation. Backbone relaxation is substantially more expensive and, for HERMES-*relaxed* models, yielded no performance improvement (Fig. S1).

### Predicting mutational effects on thermodynamic fold-stability

The chief task on which we evaluated HERMES was predicting mutational effects on thermodynamic folding stability, defined as the change in Gibbs free energy Δ*G* upon folding. The effect of a mutation wt → mt is thus ΔΔ*G*_wt→mt_ = Δ*G*_mt_ − Δ*G*_wt_. To enable a direct comparison with recent structure-based stability predictors, we fine-tuned and evaluated HERMES on the same benchmark splits used by RaSP [2], Stability-Oracle [27], and ThermoMPNN [30]. Specifically, we (i) trained on the RaSP Rosetta-derived stability dataset and tested on RaSP’s experimental benchmark, (ii) used Stability-Oracle’s curated cDNA117k training set and T2837 test set, and (iii) trained on ThermoMPNN’s Megascale training set and evaluated on its Megascale test set (Methods).

#### Noise and side-chain relaxation improve zero-shot model predictions

We first evaluated the *zero-shot* HERMES models on the T2837 and Megascale test sets (Fig. 2, S3). Pre-training on structures with Gaussian coordinate noise (0.5 Å s.d.) improves performance, consistent with prior work [26, 35]. Relative to HERMES-*fixed*, the packing-aware HERMES-*relaxed* achieves significantly higher recall (0.48 vs. 0.27; p *<* 0.01) at only a slight cost in precision, and thus a higher F1. HERMES-*amortized* performs similarly to HERMES-*relaxed* on T2837, and on Megascale shows slightly lower recall and F1 while still outperforming HERMES-*fixed* (p-values in Fig. S4). Moreover, ensembling the zero-shot predictions across 10 independently trained models did not consistently improve performance for either HERMES-*fixed* or HERMES-*amortized* (Fig. S5A,B).

To test whether packing awareness preferentially helps substitutions that perturb local packing, we stratified mutations by residue size—assigning wild-type and mutant residues to small, medium, or large classes by normalized van der Waals volume [39]—and evaluated stabilizing-versus-destabilizing classification within each wt → mt size-transition group (Fig. 3). Switching from HERMES-*fixed* to HERMES-*relaxed* yielded the largest recall gains for stabilizing substitutions between residues of substantially different sizes. This was most pronounced for large → small substitutions: HERMES-*fixed*, constrained by the rigid wild-type cavity, cannot recognize that void-creating mutations may be stabilizing, whereas HERMES-*relaxed* repacks neighbors to fill the space. Precision changes varied across categories, but F1 improved in all, with the largest gains for small→large substitutions, where relaxation reorganizes the local environment to accommodate bulkier side chains that would otherwise clash. Interestingly, HERMES-*amortized*, which learned side-chain packing implicitly through distillation fine-tuning, showed comparable large → small gains but much weaker small→ large gains. This disparity is unlikely to stem from imbalance in the distillation data, as stabilizing small → large substitutions are not underrepresented relative to large → small ones—if anything, they are more numerous (Figure S6). We speculate instead that this reflects either an inherent limitation of distillation or a need for more distillation data overall. P-values for performance differences between models within each size category, and within models across categories, are respectively in Figs. S7, S8.

**Figure 2.**
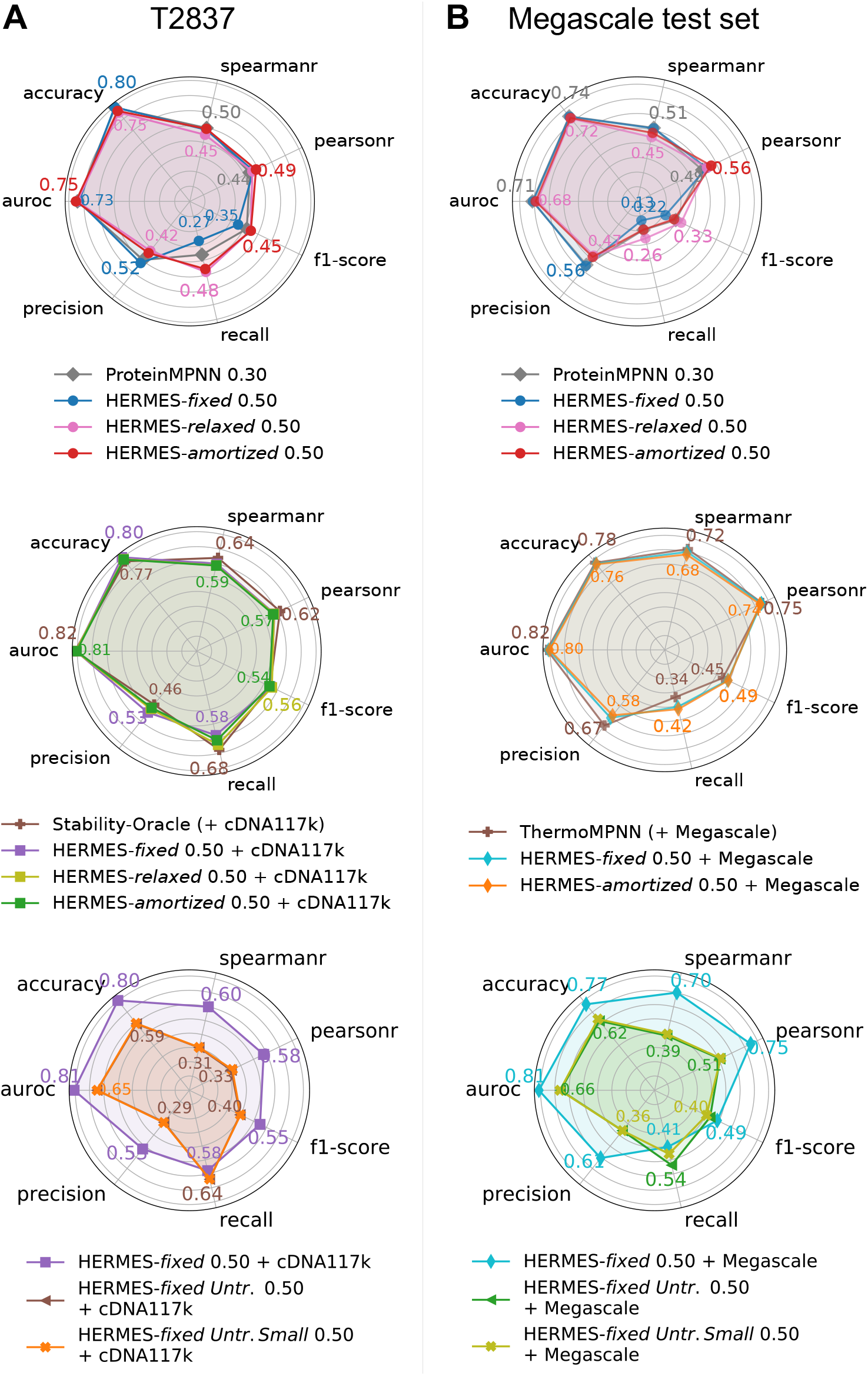
Predicting mutational effects on thermodynamic folding stability. Stabilizing-versus-destabilizing classification metrics are computed using ΔΔ*G <* 0 (experimental) and Δ log *p >* 0 (predicted) as cutoffs for stabilizing mutations. **(A)** T2837 results: zero-shot models (top) and models fine-tuned or only trained on cDNA117k (middle and bottom). **(B)** Megascale test set results: zero-shot models (top) and models fine-tuned or only trained on the Megascale training set (middle and bottom). Model names indicate the architecture, the coordinate-noise amplitude used, and when applicable, the fine-tuning dataset (listed after “+”); *Untr*. is short for *Untrained*, indicating models that had no pre-training and were instead only trained on stability effects. Only models trained with coordinate noise are shown; the noise amplitude is indicated within each model name as standard deviation in Å units. Results for models trained without noise are provided in Fig. S3.

**Figure 3.**
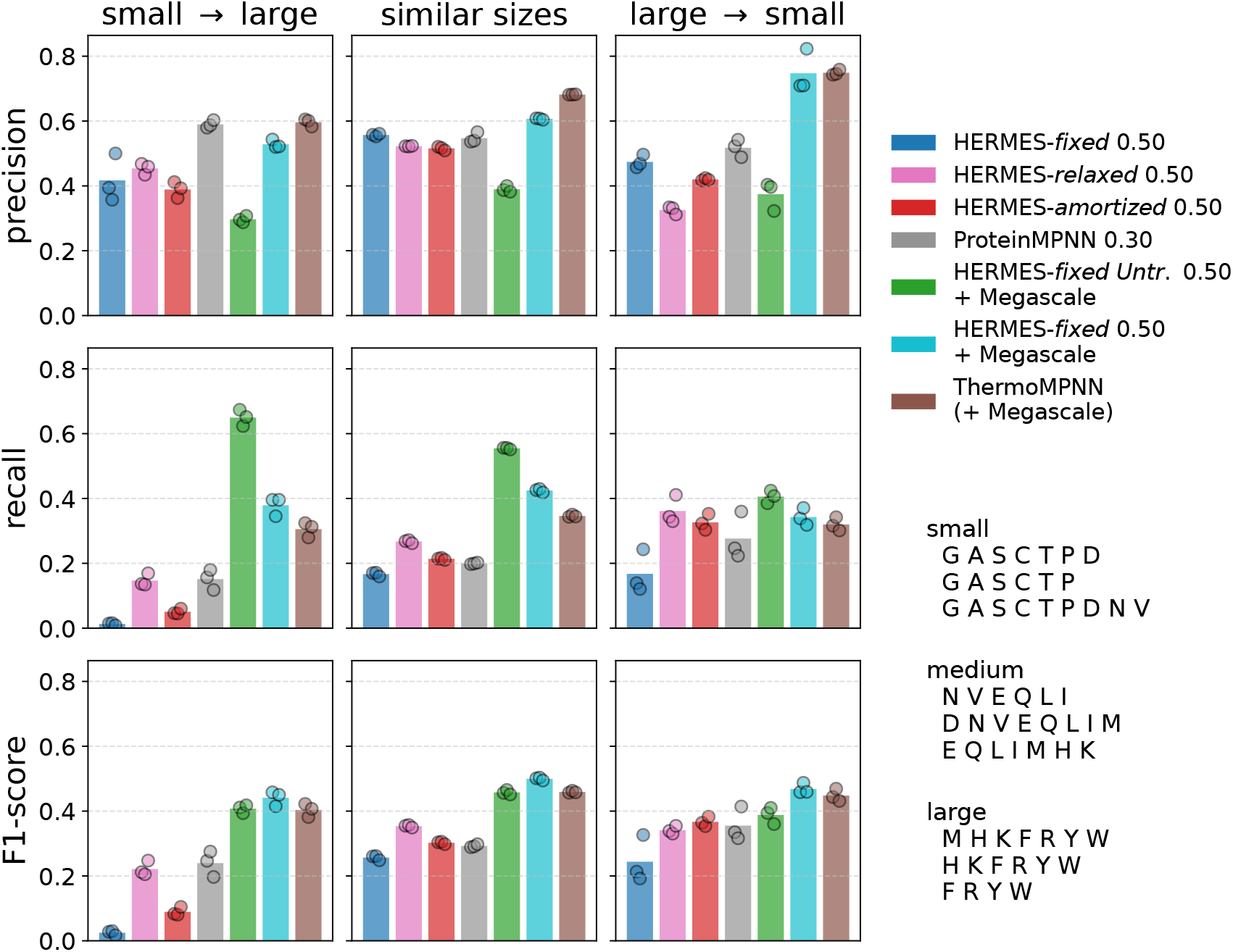
Impact of amino acid size changes on predicting mutational effects on protein stability. Amino acids are grouped into three size classes by normalized van der Waals volume [39], and predictions are stratified by the size classes of the wild-type and mutant residues; “similar sizes” denotes substitutions within the same class. Classification metrics for stabilizing vs. destabilizing mutations (rows) are computed for each stratum (columns), using ΔΔ*G <* 0 (experimental) and Δ log *p >* 0 (predicted) as thresholds for stabilizing mutations. For sensitivity analysis, predictions are bootstrapped under three alternative assignments of amino acids to size classes (bottom right). P-values for pairwise model comparisons are shown in Fig. S7, and p-values comparing each model’s performance on small → large versus large → small substitutions in Fig. S8. Model names indicate the architecture, the coordinate-noise amplitude used during pre-training (standard deviation, in Å), and, where applicable, the fine-tuning dataset (listed after “+”).

For comparison, we evaluated ProteinMPNN [35] on the same tasks. It outperformed HERMES-*fixed* and matched HERMES-*relaxed* on both small → large and large → small mutations (Fig. 3). We attribute this to ProteinMPNN’s input representation: by conditioning only on backbone coordinates and residue identities, ProteinMPNN is less constrained by explicit wild-type side-chain geometry than the all-atom HERMES-*fixed*, enabling implicit reasoning about side-chain flexibility comparable to explicit relaxation in HERMES-*relaxed*. Notably, ProteinMPNN outperformed HERMES-*amortized* on small → large substitutions.

Lastly, to assess the impact of pre-training memorization, we stratified performance by each test protein’s maximum sequence similarity to the models’ pre-training proteins (similarity distributions are shown in Fig. S9, and stratified performances are shown in Figs. S10 and S11). HERMES models are not significantly affected by similarity to the CASP12 pre-training proteins, and HERMES-*amortized* is likewise independent of similarity to the distillation fine-tuning proteins. ProteinMPNN, in contrast, performs better on proteins more similar to its pre-training set, and its performance drops more than HERMES’s in the low-similarity regime—though their performances are not significantly different (Fig. S12).

#### Fine-tuned HERMES models achieve state-of-the-art performance for stability effect prediction

When fine-tuned on the respective stability datasets, HERMES outperformed RaSP (Fig. S13) and matched Stability-Oracle (Fig. 2A, S14) and ThermoMPNN (Fig. 2B), underscoring both the effectiveness of the HERMES architecture and the importance of fine-tuning data for test-time accuracy. HERMES was also robust to ESMFold-resolved structures [40] for either fine-tuning or inference (Fig. S15), supporting use cases where only computationally predicted structures are available. Contrary to zero-shot models, ensembling across 10 fine-tuned networks provided consistent, though marginal, improvements across performance metrics (Fig. S5A, B).

Training a model directly on stability prediction, without HERMES’s amino acid classification pre-training, yielded poor performance on both cDNA117K and Megascale, even after reducing capacity from 3.5M to 50K parameters (Fig. S3). Since shrinking the model did not recover performance, overfitting alone cannot explain the gap; a similar failure mode was reported for ThermoMPNN on Megascale [30]. We take this as evidence that pre-training supplies an inductive bias the stability data are too small to provide: expressive representations of local structural environments that a modest fine-tuning set can reshape into predictors of mutational stability effects, consistent with prior work on transfer learning [27, 30]. Learning both protein-environment structure and its mapping to stability from stability labels alone would likely require substantially more data.

Finally, fine-tuning on stability effects largely eliminated size-dependent bias, yielding comparable recall and F1 scores for small→ large and large → small substitutions (Figs. 3, S8).

### Identifying model biases associated with different amino acid properties

Next we asked whether models favor substitutions associated with specific physico-chemical changes and, if so, whether such preferences reflect true experimental signal or implicit biases learned from architecture or general training.

To this end, we constructed model-average prediction matrices *M* ^*model*^ with entries 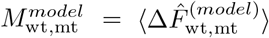, the model’s average prediction for a wt → mt substitution over the Megascale test set (Fig. 4A). For each model we built three such matrices: (i) over all Megascale sites, (ii) over core sites only (SASA *<* 1 Å^2^), and (iii) over surface sites only (SASA *>* 3 Å^2^). Despite differing value scales across models, each matrix quantifies how strongly a model favors each wt → mt substitution, averaged over distinct structural contexts.

**Figure 4.**
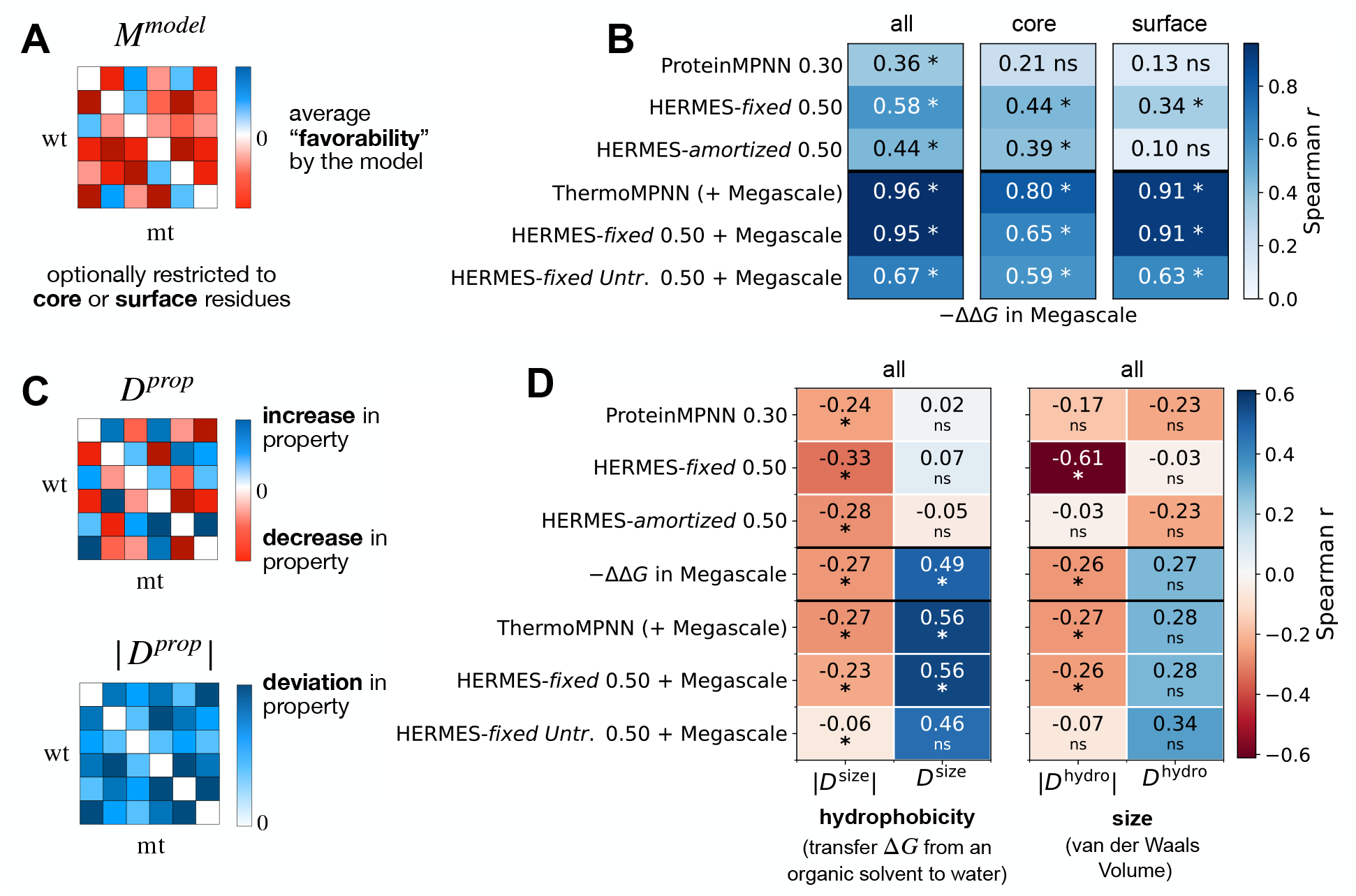
Identifying model biases for different amino acid properties. **(A)** A model’s average prediction matrix *M* ^model^ with entries 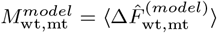 quantifies how much the model favors a given mutation wt → mt. We construct these matrices by averaging model predictions across all sites in the Megascale test set, for which experimental stability values ΔΔ*G* are also available. We consider all sites, as well as the stratified subsets of core sites (SASA *<* 1 Å^2^) and surface sites (SASA *>* 3 Å^2^). The complete matrices of all pairwise amino acid preferences, for each model and for the experimental ΔΔ*G* data, are shown in Figs. S16 and S17. **(B)** Spearman correlations between each model’s matrix and the matrix of average − ΔΔ*G* values, shown for the three site classes (all, core, surface). A strong positive correlation denotes similar mutation preferences between the model and the experimental stability measurements. Stars denote significance (*q <* 0.05; Mantel permutation test with 5000 permutations and FDR-BH correction applied across the whole plot). **(C)** For a given amino acid property (e.g., hydrophobicity or size), we compute a 20 *×* 20 matrix *D*^*prop*^, where each element 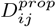 is the change in that property upon mutation from the wild-type *i* to the mutant amino acid *j*; |*D*^*prop*^ |is the corresponding element-wise absolute value. Thus |*D*^*prop*^ | quantifies the absolute change in property value upon mutation, while *D*^*prop*^ quantifies its increase or decrease. **(D)** Spearman correlations of each model’s matrix *M* ^model^ with the property matrices |*D*^*prop*^ | and *D*^*prop*^, for two representative properties: hydrophobicity, measured by the transfer Δ*G* from an organic solvent to water (left), and size, measured by van der Waals volume (right). A positive correlation between *M* ^model^ and |*D*^*prop*^ | indicates that the model favors property-changing mutations, whereas a negative correlation indicates a bias toward property-preserving mutations. A positive (negative) correlation between *M* ^model^ and *D*^*prop*^ indicates a bias toward increasing (decreasing) the property. Stars denote significance (*q <* 0.05; Mantel permutation test with 5000 permutations and FDR-BH correction applied across the whole plot). The complete correlation matrices between all 50 amino acid properties (from ref. [39]) and the model matrices, stratified by residue location (core vs. surface), are shown in Fig. S18.

First, to test whether models’ preferences align with biases in experimental stability measurements, we compared each model matrix against a matrix of experimental − ΔΔ*G* values averaged over the same Megascale test sites. Among zero-shot models, HERMES-*fixed* correlated most strongly with experiment (Fig. 4B). Fine-tuning on the Megascale training set further boosted this correlation for both HERMES-*fixed* and ProteinMPNN (Spearman *r* = 0.95–0.96), as their predictions shifted from amino acid propensities toward stability measurements (Fig. 4B). By contrast, the model trained on stability data alone, without amino acid classification pre-training, correlated more weakly (Spearman *r* = 0.67; Fig. 4B), underscoring the limited structural reasoning attainable from stability data alone, consistent with Fig. 2. Notably, although this un-pre-trained model moderately recovered the experimental mutation ranking, it disproportionately predicted mutations to be stabilizing (positive entries in its *M* ^*model*^), unlike all other models and experiment, which identified mutations to be on average destabilizing (Figs. S16 and S17). Finally, zero-shot models aligned with experimental matrices more strongly at core sites, whereas fine-tuned models aligned more strongly at surface sites (Fig. 4B).

Next, we asked whether these preferences are linked to specific physico-chemical properties of the wild-type and mutant amino acids (Fig. 4C). In Fig. 4D we focused on hydrophobicity and size, quantified respectively by the transfer Δ*G* from an organic solvent to water and the Van der Waals volume [39]. Following ref. [41], we constructed two matrices for each property (*prop*): |*D*^*prop*^|, with entries |*D*^*prop*^|_wt,mt_ = |*prop*(mt) − *prop*(wt)|, the absolute change upon mutation; and *D*^*prop*^, with entries 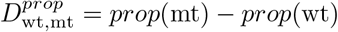, the signed change. Figure 4D shows the correlation of each *M* ^*model*^ with both. Correlation with |*D*^*prop*^ |indicates whether a model favors mutations that differ (*r >* 0) or are similar (*r <* 0) in property value; correlation with *D*^*prop*^ indicates whether it favors increasing (*r >* 0) or decreasing (*r <* 0) that value.

Several observations follow. On size, zero-shot HERMES-*fixed* strongly favors size-preserving mutations (Spearman *r* = − 0.61 with |*D*^size^ |, Fig. 4D), with no directional preference (*r* = − 0.03 with *D*^size^), consistent with the model’s architecture, which recovers a masked amino acid from its rigid atomic neighborhood. In contrast, ProteinMPNN and HERMES-*amortized* are virtually free of size bias, aside from a slight HERMES-*amortized* preference for smaller amino acids at surface residues (Spearman *r* = − 0.39 with *D*^size^, Fig. S18). Fine-tuned models slightly favor preserving size (Spearman *r* = − 0.28 with |*D*^size^|), tracking the experimental ΔΔ*G* measurements and are consistent across core and surface residues (Fig. S18).

On hydrophobicity, models differ markedly. Experiments and fine-tuned models prefer substitutions that *increase* hydrophobicity (Spearman *r* = 0.49–0.56 with *D*^hydro^, Fig. 4D), whereas zero-shot models show no directional preference.

Stratifying between core and surface residues, biophysical intuition would predict a preference for *decreasing* hydrophobicity at the surface. All zero-shot models show this, with HERMES-*amortized* most strongly among them (Spearman *r* = − 0.46 with *D*^hydro^), whereas experiments and fine-tuned models mildly favor *increasing* it (Spearman *r* = 0.28 with *D*^hydro^)—possibly reflecting a need to keep the small Megascale proteins soluble [42] (Fig. S18).

Beyond the two properties we examined in detail here—the Gibbs energy of transfer from an organic solvent to water, and the Van der Waals volume—we performed the same analysis for the other 48 amino acid properties specified in ref. [39] and listed in Table S3, which broadly capture hydrophobic, steric, and electronic characteristics. Figure S18 compares each model’s biases against the shifts associated with all these properties. For hydrophobic properties, the inferred biases are generally consistent across properties within each model, whereas steric and electronic biases are more heterogeneous, hinting at greater variability among the properties associated with each of these categories. Taken together, these analyses show models’ biases and how they relate to the physico-chemical basis of protein stability, biases in the training data, and architectural choices.

### Reversibility and path-independence of HERMES predictions with respect to mutations

Mutational effects are, in principle, reversible: the *α* → *β* effect should equal the reverse in magnitude and oppose it in sign, Δ*F*_*α*→*β*_ =− Δ*F*_*β*→*α*_. Moreover, equilibrium quantities like protein stability free energy are state functions, so a multi-step substitution’s net effect depends only on the initial and final amino acids, not the path: Δ*F*_*α*→*γ*_ = Δ*F*_*α*→*β*_ + Δ*F*_*β*→*γ*_ for a path *α*→ *β*→ *γ*.

HERMES predicts mutation effects (both zero-shot and fine-tuned) as differences in amino acid-specific log-probabilities log *p*(*aa*|*X*_*aa*_), which under a fixed structural context satisfy reversibility by construction:

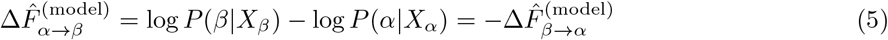

where *X*_*α*_ = *X*_*β*_ for HERMES-*fixed* and HERMES-*amortized*, and 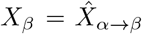 for HERMES-*relaxed*; path-independence follows similarly. By contrast, other structural models such as Stability-Oracle [27] require explicit *×* 19 data augmentation (“thermodynamic permutation augmentation”) to enforce these properties.

Alternatively, reversibility can be assessed in a structure-conditioned manner [2, 30], computing forward (*α* → *β*) and reverse (*β* → *α*) effects from their outgoing contexts—*α* → *β* conditioned on *X*_*α*_ and *β* → *α* on *X*_*β*_ (Methods). Here reversibility is not automatic, since the two directions use different structures, but a well-balanced model should be near-reversible, making this a stringent test of bias. We measured it on the Ssym dataset (wild-type and single-mutant structures for 352 mutations across 19 proteins [43]), computing for each mutation a “forward” effect (wt → mt, on the wild-type structure) and a “reverse” effect (mt → wt, on the mutant structure).

Zero-shot HERMES models and ProteinMPNN predict stability effects more accurately for forward (wt → mt) than reverse substitutions, consistent with prior observations [2, 30] (Fig. 5A). Adding coordinate noise during pre-training and fine-tuning on stability effects each reduced this forward-reverse disparity but did not eliminate it, while models trained *only* on stability effects showed little disparity, albeit with substantially worse performance. Across all pre-trained models, forward effects from the wild-type have larger magnitudes than the corresponding reverse effects (Fig. 5B, scatterplots), a bias stemming from the elevated log-probabilities assigned to wild-type amino acids in wild-type neighborhoods log *p*(wt | *X*_wt_) (Fig. 5B, density plots)—a wild-type preference that may reflect pre-training memorization. Consistently, stratifying Ssym proteins by sequence similarity to the pre-training set (Methods) modestly reduced this preference for lower-similarity proteins (Fig. 5B). A more detailed characterization is left to future work.

**Figure 5.**
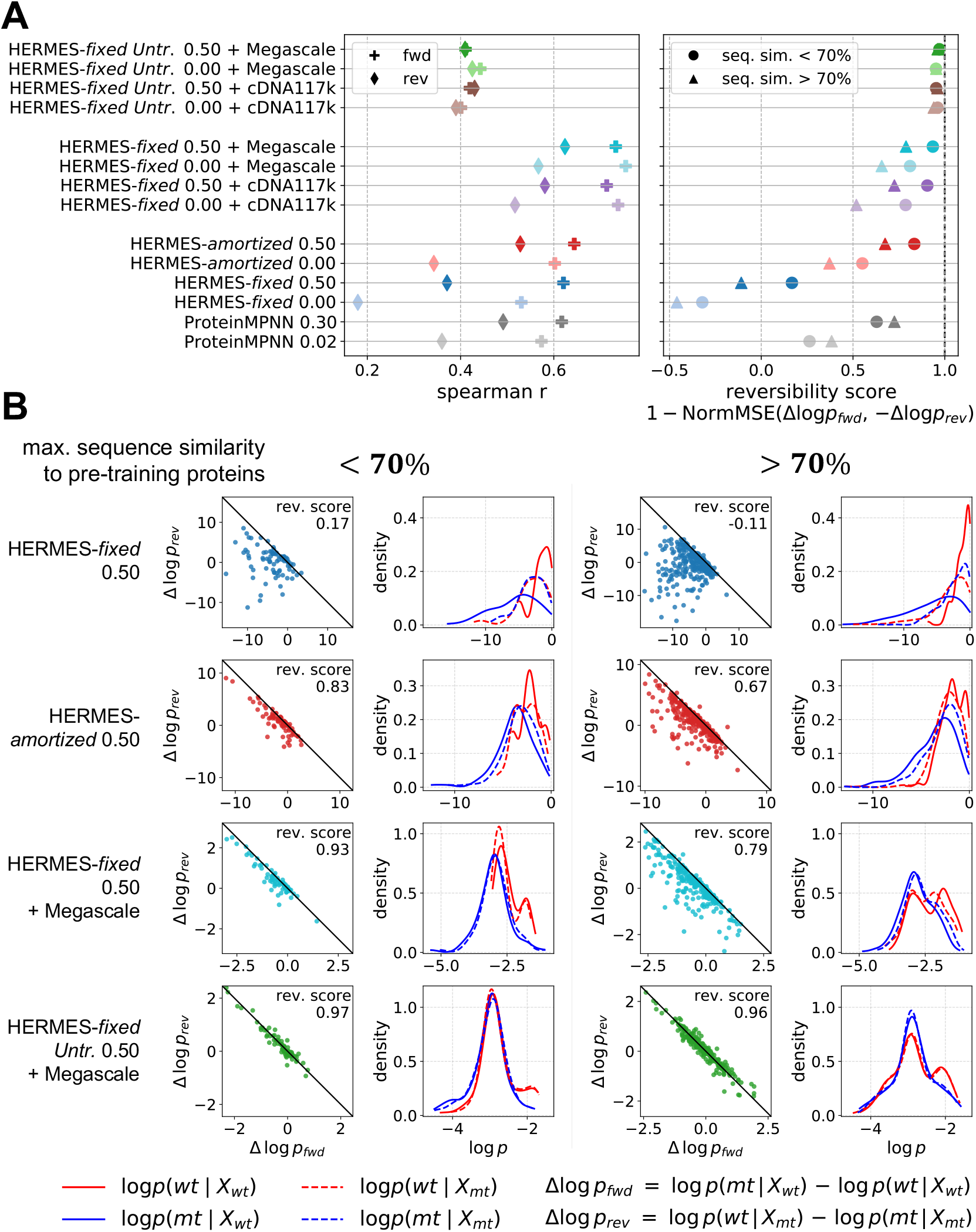
Identifying model biases with structure-conditioned reversibility. **(A)** (Left) Spearman correlations between experimental stability changes ΔΔ*G* and model predictions on the Ssym dataset. For each model, we report correlations for forward substitutions (Δ log *p*_fwd_ vs. ΔΔ*G*) and reverse substitutions (Δ log *p*_rev_ vs. − ΔΔ*G*). (Right) Reversibility scores across all models (higher is more reversible; Eq. 9), stratified by each protein’s maximum sequence similarity to the pre-training proteins (≥ 70% vs. *<* 70%). **(B)** For each similarity split (columns) and four representative HERMES models (rows), the left panel shows scatter plots of Δ log *p*_fwd_ vs. Δ log *p*_rev_; an unbiased model should show strong anti-correlation, as summarized by the reversibility score. Reversibility is highest for the model not pre-trained on wild-type amino acid classification (green; bottom row) and is consistently higher for the low-similarity subset. In each column, the right panel shows the distributions of the log-probabilities composing Δ log *p*_fwd_ = log *p*(mt | *X*_wt_) − log *p*(wt | *X*_wt_) (solid lines) and Δ log *p*_rev_ = log *p*(wt | *X*_mt_) − log *p*(mt | *X*_mt_) (dashed lines). All models except the one *not* pre-trained on wildtype amino acid classification (bottom row) exhibit elevated log *p*(wt | *X*_wt_).

### Predicting antigen stabilization with HERMES for vaccine design

Structure-based vaccine design seeks to increase vaccine efficacy by engineering viral envelope glycoproteins (antigens) to be more stable in their pre-fusion than post-fusion conformation (antigen stabilization), thereby enriching presentation of neutralization-relevant epitopes and biasing elicited immune responses toward protective specificities. In practice, this introduces a minimal set of mutations to the fusion protein that improve pre-fusion stability and, optionally, decrease post-fusion stability, while preserving the relevant epitopes [9–13]. Mutations are often classified by stabilization mechanism: most increase prefusion stability with little regard for post-fusion, while some–notably proline mutations–also de-stabilize post-fusion conformations (see Fig. S20A for examples).

Because experimental antigen-stabilization campaigns are costly and screening assays are low-throughput [44–46], computational modeling has been used to design mutational libraries for targeted screening. These efforts have relied largely on physics-based scoring functions such as Rosetta [10, 13], with a few closed-source machine-learning attempts [11]. Here, we test whether HERMES can serve as an open-source tool for such library design campaigns.

Because no large-scale datasets of validated stabilizing mutations exist, we manually collected 33 previously reported antigen-stabilizing mutations across five viral antigens with reliable pre-fusion structures in the PDB. We evaluated how well different models rank these validated mutations as pre-fusion stabilizing within their sites, categorizing each predicted stabilizing mutation as strongly or moderately suggested, corresponding to per-site top-3 (*r*_mt_ ≤ 3) and top-6 (*r*_mt_ ≤ 6) ranked recall, respectively (Methods).

Figures 6A, B, S21, S19 and Table S4 summarize predictions across the 33 stabilizing mutations. Among the machine-learning methods, HERMES-*amortized* performs best: it ranks 24 stabilizing mutations better (lower) than wild-type, with 19 of them strongly suggested (*r*_mt_ ≤ 3). Notably, stability fine-tuning does not improve performance over the corresponding zero-shot models on this task (Table S4), possibly due to a distribution mismatch: fine-tuning mutations are mostly surface mutations from the small proteins that constitute the Megascale database, whereas antigen-stabilizing mutations are mostly buried (Fig. S22).

**Figure 6.**
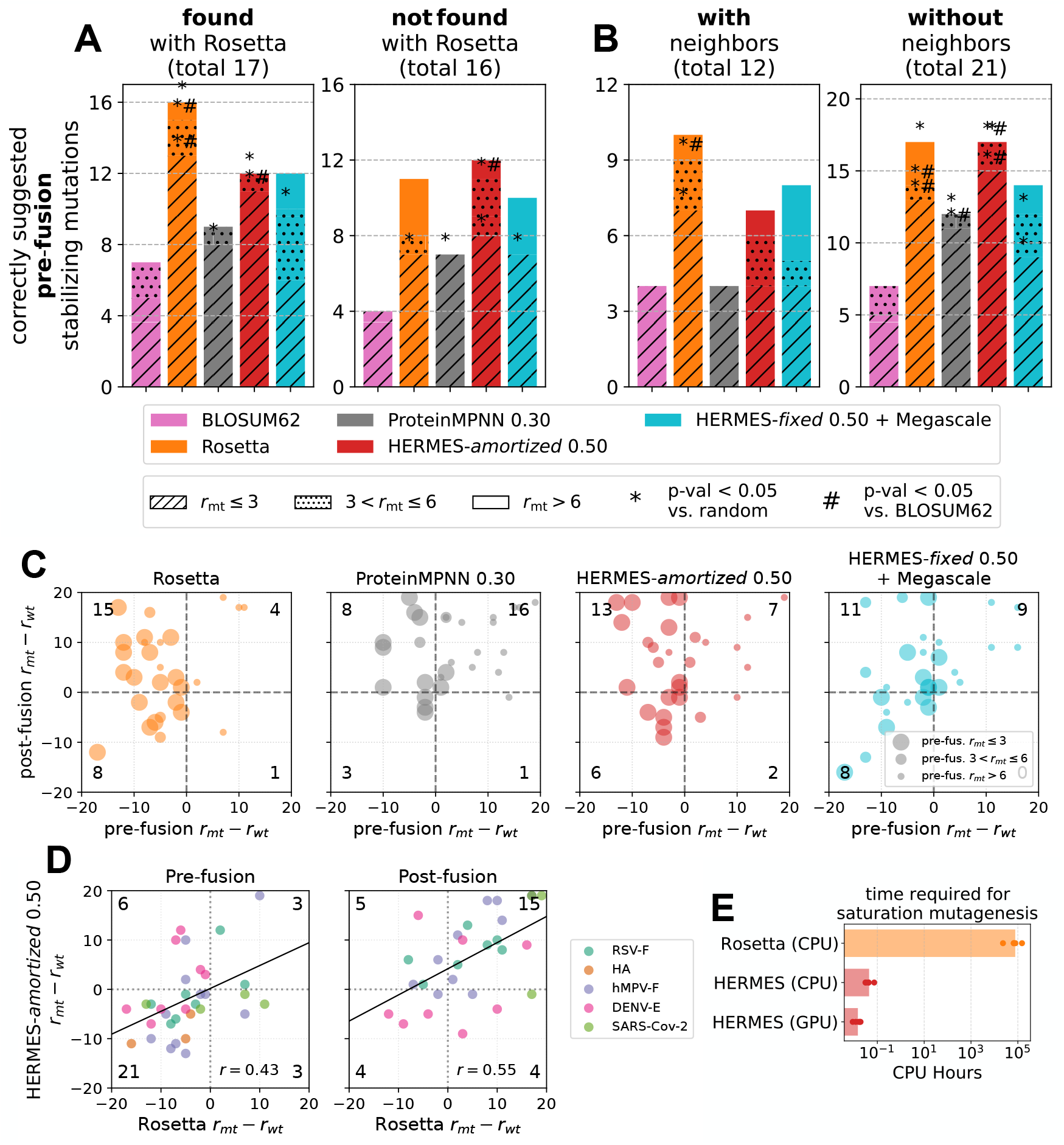
Retrospective analysis to predict pre-fusion antigen-stabilizing mutations. We consider 33 mutations across 5 diverse viral antigens, each experimentally validated to stabilize the pre-fusion conformation relative to the post-fusion conformation, as reported in refs. [9–13]. We quantify each model’s ability to recognize these validated mutations as pre-fusion-stabilizing as follows. For each mutated site, we rank every possible amino acid substitution by the model-predicted score and compare the rank of the validated mutant (*r*_mt_) to that of the wild-type (*r*_wt_). A *negative* (*positive*) rank difference predicts the substitution to be *favorable* (*unfavorable*) given the input structure. We evaluate predictions on both the pre-fusion and post-fusion structures; only 28 of the 33 mutations had a matching post-fusion structure (see Fig. S19 for PDB IDs). **(A, B)** Counts of correctly prioritized pre-fusion-stabilizing mutations (pre-fusion *r*_mt_ *< r*_wt_), stratified for each model by predicted rank group: strongly suggested (*r*_mt_ ≤ 3), moderately suggested (3 *< r*_mt_ ≤ 6), or weakly suggested (*r*_mt_ *>* 6). In (A), performance is further stratified by whether each mutation was originally identified using Rosetta in the study of origin. In (B), it is stratified by whether the mutation has close neighbors (within 10 Å) in the stabilized protein complex (Methods). For comparison, BLOSUM62 is used to rank substitutions into the (*r*_mt_ ≤ 3) and (3 *< r*_mt_ ≤ 6) groups; under this scheme the wild-type residue is always ranked first. Statistical enrichment is assessed by comparing the number of strong and moderate recalls to a random-ranking baseline and to BLOSUM62 (denoted by ∗ and #, respectively, when the binomial-test corrected *p*-value *<* 0.05; see Methods). Fig. S19 shows each model’s predictions for all 33 mutations. **(C)** Pre-fusion versus post-fusion rank differences, as predicted by four models for the 28 mutations. Mutations in the top-left quadrant are predicted to be both favorable for the pre-fusion and unfavorable for the post-fusion conformation (the desired combination). **(D)** HERMES-*amortized* versus Rosetta rank differences. Predictions by the two models are mildly correlated (Pearson *r* shown). **(E)** Estimated times for saturation-mutagenesis predictions on the pre-fusion conformations of the viral antigens considered in this study; each dot is an antigen. Times are for a single CPU and a single NVIDIA A40 GPU. See Table S5 for precise per-antigen times.

For comparison, we also scored each mutant with Rosetta [34] (Methods), which recovers stabilizing mutations on par with HERMES-*amortized* (Fig. 6A and Table S4). However, 17 of these variants were originally selected using Rosetta-based screening, so Rosetta achieves near-perfect recall on them but significantly lower recall on the rest (Fig. 6A). Moreover, Rosetta is far more computationally expensive than the machine-learning models, limiting its utility for high-throughput saturation mutagenesis library design (Table S5, Fig. 6E).

All models except HERMES-*fixed* and ThermoMPNN significantly outperform random ranking in recovering strong/moderate stabilizing mutations (Table S4). As an additional baseline, the BLOSUM62 substitution matrix (Methods) recovers significantly fewer strong/moderate stabilizing mutations than HERMES-*amortized* (Fig. 6), underscoring the value of structural context for predicting stabilizing substitutions; all binomial-test p-values in Fig. S24.

We also note that two of the five wild-type antigens have homologs above 50% sequence identity in HERMES’ pre-training data (one above 95%), versus four with close homologs in ProteinMPNN’s (two above 95%; see Table S7). However, given the small sample size, we cannot assess how such sequence homology might contribute to the models’ predictions (Fig. S23).

Next, we examined model performance across mutation types (Table S6, Fig. S20B). HERMES*amortized* most reliably recovers pre-fusion-stabilizing proline mutations (7 of 8), and all models recover proline and electrostatic mutations above the BLOSUM62 baseline. Cavity-filling mutations show weaker gains, likely because such substitutions (a smaller residue replaced by a larger hydrophobic one) are common in natural sequences, and thus, partly captured by BLOSUM62, whereas proline and charged substitutions are more context-dependent and benefit more from structure-aware modeling; see SI for a per-antigen discussion.

Predicting synergistic mutations (i.e., those reported to stabilize the complex through specific interactions between them) is difficult for nearly all methods (Fig. S20B, SI). Because our evaluation scores each site independently, holding other residues fixed, HERMES cannot capture stabilization from synergistic combinations of mutations; ProteinMPNN, ThermoMPNN, and Rosetta are held to the same protocol for fairness, though they can in principle score multi-residue variants jointly. HERMES’s predictions are affected whenever two mutated residues lie close in space, regardless of true synergy: when predicting at residue A, a neighboring mutated residue B appears in A’s local environment with its *wildtype* rather than mutant identity, biasing the prediction at A. If the two residues are far enough apart, neither appears in the other’s environment, so they fo not directly affect each other’s predictions—though HERMES would still miss any allosteric synergy between them. To quantify this effect, we stratified performance by whether mutations had close neighbors in the stabilized antigen, finding, as expected, reduced performance in that setting for both HERMES and ProteinMPNN (Fig. 6B; see Methods for details).

Although stabilizing the pre-fusion conformation is the primary mechanism for effective antigen design, destabilizing the post-fusion conformation is also a viable strategy and often pursued jointly [10]. We therefore ran predictions on post-fusion structures where available (28 of 33 mutations). HERMES-*amortized* and Rosetta predict the most mutations to be simultaneously pre-fusion-stabilizing and post-fusion-destabilizing (Figs. 6C and S19). While the true post-fusion effects are unknown, these results highlight both models’ potential to identify mutations with the desired dual effect.

So far, our recall analysis has identified HERMES-*amortized* and Rosetta as the two best models at recovering known antigen-stabilizing mutations. However, per-site recall on a small set of pre-verified mutations does not directly reflect performance in a design campaign, where precision across all candidate sites is equally critical. To demonstrate design utility, we devised workflows for proposing novel antigen-stabilizing mutations. Because HERMES-*amortized* and Rosetta have similar but complementary predictive power (Fig. 6D), we hypothesized that combining them could improve the likelihood of success [47]. Since Rosetta is too slow to score all possible mutations under reasonable computational budgets (Fig. 6E), we screen all mutations with HERMES-*amortized* and filter the top hits with Rosetta, whose track record in identifying experimentally validated antigen-stabilizing mutations supports its use as a filter [10, 13].

Given the difficulty of large-scale experimental validation, we use Rosetta as an oracle to assess HERMES-based library design. This is a common strategy in computational protein design when experimental screens are inaccessible (analogous to using AlphaFold as an oracle elsewhere [48]). We designed two HERMES-*amortized* pipelines—one stabilizing the pre-fusion core, the other additionally destabilizing post-fusion—and benchmarked them against ProteinMPNN, BLOSUM62, and random baselines, scoring each suggested mutation with Rosetta to define “Rosetta hits” (Methods).

On RSV-F and DENV-E (Figs. 7 and S25), HERMES-*amortized* yields more Rosetta hits than all baselines, except ProteinMPNN in the “stabilize pre-fusion, destabilize post-fusion” pipeline. Agreement with Rosetta is strongest in the “stabilize pre-fusion core” pipeline, where 7 of the top 10 mutations are Rosetta-validated, versus 4 for ProteinMPNN and 2 at random (Fig. 7). HERMES-*amortized* suggests both cavity-filling mutations in the core (small hydrophobic → large hydrophobic, polar→ hydrophobic) and electrostatic mutations (polar → charged) that Rosetta agrees with (Fig. 7A, middle). Mutations suggested by HERMES-*amortized* are listed in Supplementary Data 1.

**Figure 7.**
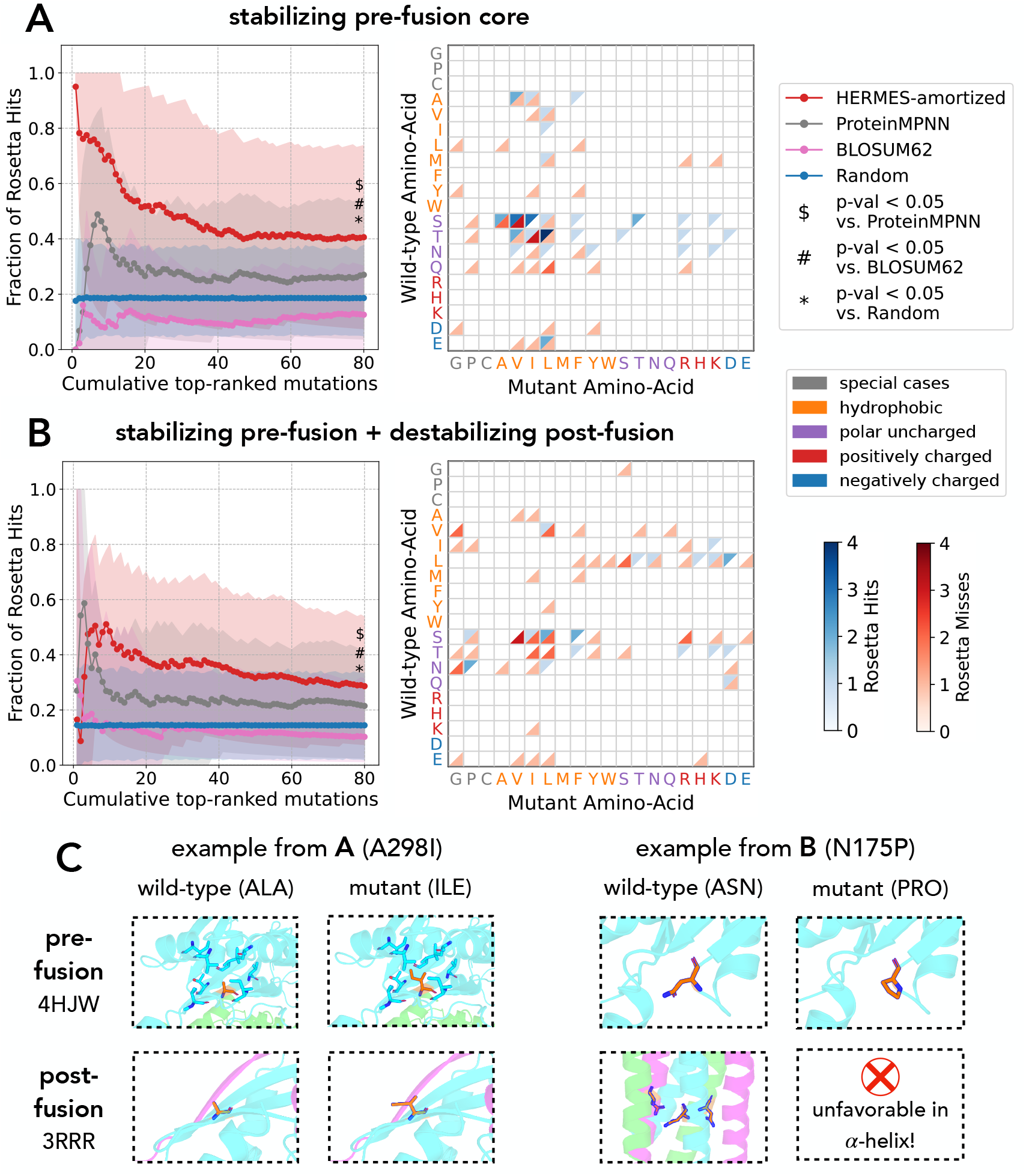
Computational workflow with HERMES for designing a mutational library for antigen stabilization in RSV-F. We evaluate computational workflows for designing antigen-stabilizing mutations in RSV-F, in which candidate mutations are suggested by each model (colors) and Rosetta is then used as an oracle to assess the success of the suggested mutations per model. We assess two workflows in which we suggest **(A)** stabilizing mutations at the core of the pre-fusion structure, and **(B)** mutations that stabilize the pre-fusion and destabilize the post-fusion conformation (Methods). **(A, B)** Library-design results for each model using (A) the “stabilize pre-fusion core” pipeline and (B) the “stabilize pre-fusion and destabilize post-fusion” pipeline. **Left:** Rosetta hit rate as a function of the number of top-*k* suggested mutations, shown for both protocols. For all methods except Random, mutations are ranked by score; for Random, we show the average hit rate over 20 random samples of *k* mutations drawn from a pool of 160 randomly selected mutations. Rosetta hit rates were computed over 200 bootstrap samples (with replacement) of the 50 REU scores; solid lines and markers denote the mean across bootstrap samples, and shaded regions denote the 5–95 percentile range. All BLOSUM62 suggestions had a score ≥ 1 and are ranked by score, as for HERMES and ProteinMPNN. P-values were computed via a Wilcoxon signed-rank test and Bonferroni-corrected for multiple testing within each panel. **Right:** Heatmaps show the number of Rosetta hits (blue) and misses (red) for each substitution (pair of amino acids: wild-type → mutant) suggested by HERMES-*amortized*; here, hits were computed using all 50 REU scores rather than bootstrap samples. **(C)** Examples of promising antigen-stabilizing mutations identified by each pipeline. A298I is cavity-filling for pre-fusion, while likely being neutral for post-fusion due to being at the surface. N175P destabilizes post-fusion by placing a proline in a *α*-helix.

Among the HERMES-suggested, Rosetta-validated mutations, A298I and N175P stand out (Fig. 7). In pre-fusion, A298 lies in an under-packed core region surrounded by hydrophobic residues (bar a single threonine), and mutation to isoleucine improves packing without requiring neighborhood relaxation. Although identified without regard for post-fusion, site 298 lies on the post-fusion surface, so isoleucine would likely not further stabilize post-fusion (Fig. 7A, right). We subsequently found A298I covered in a patent [49], corroborating its stabilization potential. N175P, from the post-fusion-aware pipeline, is a canonical proline substitution: in pre-fusion the site sits on a loop (a sharp turn) that accommodates prolines, whereas in post-fusion it lies on an *α*-helix, which generally cannot (Fig. 7B, right).

Overall, HERMES-*amortized* can accelerate antigen-stabilization campaigns by rapidly evaluating all possible mutations and proposing a pool of high-confidence candidates for experimental validation, optionally filtered with Rosetta.

Several limitations should be noted. First, we use only relative ranks, whereas absolute score magnitudes carry additional information for more robust prioritization. Second, experimental validation frequently involved multi-mutation constructs [9–13], complicating attribution of stabilization to individual residues. Third, the extent to which each mutation stabilizes pre-fusion versus destabilizes post-fusion is often not disentangled experimentally. Fourth, the modest number of curated mutations limits the statistical power of the retrospective analysis. Nonetheless, we expect a fast HERMES-type pipeline can make library design more efficient, enabling more targeted experimental testing.

### Predicting binding effect of mutations

Predicting mutational effects on binding affinity is a central step in target-specific protein design. In prior work, we showed HERMES predicts how mutations in short peptide antigens alter binding to T-cell receptors within MHC complexes, and leveraged this for de novo design of peptide antigens that elicit specific T-cell responses [48]. Here, we evaluate HERMES on the single-point mutation effects from the SKEMPI v2.0 dataset [55], which spans a substantially broader range of protein-protein interactions and binding-affinity perturbations.

In a zero-shot setting, HERMES exhibits measurable predictive signal (HERMES-*fixed* 0.50; Spearman’s *ρ* = 0.286), comparable to ProteinMPNN, whereas physics-based approaches (Rosetta [34] and FoldX [22]) perform modestly better (Spearman’s *ρ* ≈ 0.35). In contrast to stability prediction, HERMES-*fixed* correlates more strongly with binding effects than HERMES-*amortized*. Interestingly, fine-tuning for stability slightly improves binding-effect prediction(Table 1).

**Table 1.** Benchmark on predicting mutational effects on protein-protein binding in SKEMPI. The Spearman/Pearson correlations between model-predicted effects of single point mutations and experimental values from the SKEMPI v2.0 dataset [55] are reported both across mutations within each structure individually (“per-structure” correlations), and over mutations pooled across all complexes (“overall” correlations). Entries marked with ∗ are taken from ref. [33] and, for machine learning-based methods (all except Rosetta and FoldX), were obtained using 3-fold cross-validation over SKEMPI complexes. Entries marked with *†*are taken from ref. [54], and were obtained via 5-fold cross-validation on the SKEMPI complex structures. We evaluate HERMES using three train/test splits defined from SKEMPI metadata, with increasing difficulty: (i) *Easy*, a random split; (ii) *Medium*, which groups sites with similar binding sites (hold-out proteins) into the same split; (iii) *Difficult*, which groups sites from the same held-out protein types (functional classes) into the same split (see Methods for details). The HERMES models with most comparable training procedure to the other machine learning models are those fine-tuned on the *Easy* split, for which we used 3-fold cross-validation with splits defined by PDB complex. Note that, similar to HERMES, the machine learning models in ref. [33] were fine-tuned on SKEMPI ΔΔ*G*_binding_ labels only, whereas Pythia-PPI [54] was trained on a mixture of SKEMPI binding labels and FireProtDB stability labels [56] and further refined via self-distillation.

| Method | Per-Struct.<br>Pearson | Per-Struct.<br>Spearman | Overall<br>Pearson | Overall<br>Spearman |
| --- | --- | --- | --- | --- |
| Rosetta* [50] | 0.328 | 0.299 | 0.311 | 0.347 |
| FoldX* [51] | 0.391 | 0.364 | 0.356 | 0.351 |
| DDGPred* [52] | 0.371 | 0.343 | 0.652 | 0.439 |
| ESM-IF* [53] | 0.231 | 0.209 | 0.296 | 0.287 |
| MIF-Net.* [33] | 0.395 | 0.348 | 0.667 | 0.480 |
| RDE-Net.* [33] | 0.469 | 0.433 | 0.642 | 0.527 |
| Vanilla Pythia PPI† [54] | 0.478 | 0.449 | 0.709 | 0.537 |
| Pythia-PPI† [54] | 0.565 | 0.527 | 0.785 | 0.637 |
| ProteinMPNN 0.02 | 0.281 | 0.282 | 0.331 | 0.315 |
| ProteinMPNN 0.30 | 0.270 | 0.255 | 0.334 | 0.289 |
| HERMES-*fixed* 0.00 | 0.306 | 0.287 | 0.285 | 0.272 |
| HERMES-*fixed* 0.50 | 0.317 | 0.308 | 0.291 | 0.286 |
| HERMES-*amortized* 0.00 | 0.231 | 0.241 | 0.290 | 0.242 |
| HERMES-*amortized* 0.50 | 0.222 | 0.239 | 0.276 | 0.222 |
| HERMES-*fixed* 0.00 + cDNA117k | 0.347 | 0.331 | 0.380 | 0.342 |
| HERMES-*fixed* 0.50 + cDNA117k | 0.305 | 0.294 | 0.344 | 0.288 |
| HERMES-*fixed* 0.00 + SKEMPI Easy | 0.471 | 0.433 | 0.578 | 0.476 |
| HERMES-*fixed* 0.50 + SKEMPI Easy | 0.430 | 0.389 | 0.512 | 0.420 |
| HERMES-*fixed* 0.00 + SKEMPI Medium | 0.472 | 0.430 | 0.576 | 0.466 |
| HERMES-*fixed* 0.50 + SKEMPI Medium | 0.407 | 0.368 | 0.497 | 0.403 |
| HERMES-*fixed* 0.00 + SKEMPI Difficult | 0.435 | 0.398 | 0.395 | 0.380 |
| HERMES-*fixed* 0.50 + SKEMPI Difficult | 0.399 | 0.359 | 0.328 | 0.322 |

Next, we fine-tuned HERMES directly on SKEMPI using 3-fold cross-validation. To control for information leakage from structural similarity, we introduced three homology-aware splitting strategies of increasing difficulty (*Difficult, Medium*, and *Easy*); the most stringent split (*Difficult*) prevents complexes from the same interaction “class” (e.g., antibody-antigen, TCR-pMHC) from appearing in different folds (Methods). Fine-tuned HERMES models achieve substantially stronger correlations, including under the *Difficult* split, indicating that fine-tuning captures binding-affinity patterns that generalize across interaction classes (Table 1). Stratifying by mutation location shows that both HERMES and ProteinMPNN are driven primarily by mutations at the interface *core* (as defined in ref. [57]; Fig. S26), suggesting these models are best applied to interface mutations only. This is consistent with HERMES’s strict locality: interface-proximal mutations act through local effects HERMES can model, whereas allosteric effects of distant mutations cannot be captured. Interestingly, although ProteinMPNN learns mutational patterns across the entire structure, its performance on distant mutations is no better than HERMES-*fixed*.

Finally, we compared HERMES to several baselines trained and evaluated on SKEMPI using 3- or fold cross-validation under protocols analogous to—but not identical to—our *Easy* split. Because this approach does not control for homology between train and test sets, the resulting numbers should not be taken as evidence of generalizable learning; moreover, these baselines do not report results under more stringent splits, and not all provide training code. HERMES fine-tuned on SKEMPI-*Easy* performs competitively with RDE-Network and MIF-Network [33], but trails Pythia-PPI (Table 1). RDE-Network and MIF-Network, like HERMES, are fine-tuned directly on SKEMPI ΔΔ*G* labels, whereas Pythia-PPI employs additional training procedures that further boost performance (Methods). These procedures are in principle applicable to HERMES; we leave their systematic evaluation, and a broader assessment of HERMES as a framework for binding-affinity prediction, to future work.

## Discussion

We introduced HERMES, a family of fast, structure-based machine learning models for predicting the effects of mutations on protein function. HERMES leverages SO(3)-equivariant neural networks operating on local atomic neighborhoods in a protein structure to predict amino acid propensities, whose differences can be used to predict mutational effects. HERMES captures mutational impacts on thermodynamic stability and protein-protein binding affinity, and it provides a practical tool for antigen stabilization in vaccine design.

A central finding is that encoding side-chain flexibility is essential for accurate predictions across mutations involving amino acids of different sizes. HERMES-*fixed*, a lightweight protocol that evaluates mutations in the rigid wild-type structure, exhibits strong bias toward size-conserving substitutions. Explicitly modeling relaxation via Rosetta [34], (HERMES-*relaxed* protocol) resolves this bias but incurs substantial computational cost. Our amortized approach offers an effective compromise: by fine-tuning on relaxed predictions for just 0.5% of pre-training data, HERMES-*amortized* learns implicit side-chain flexibility at HERMES-*fixed* speed.

Beyond zero-shot use, HERMES can be fine-tuned directly on experimental labels using a simple end-to-end procedure. Notably, both the native and the fine-tuned models use the difference of amino acid scores for predicting mutational effects, which yields an inherent reversibility with respect to mutation order–a thermodynamic consistency property that is other models enforce via explicit 19 *×* data augmentation [27]. With fine-tuning, HERMES achieves competitive accuracy on thermodynamic stability benchmarks when trained on the same data as prior methods [2, 27, 30].

We also demonstrate that pre-training on wild-type amino acid classification remains necessary for strong performance; models trained only on currently-available experimental stability data fail. Notably, pre-trained models exhibit “wild-type preference,” assigning elevated probabilities to wild-type residues in wild-type structural contexts. This bias correlates with sequence similarity to pre-training proteins, suggesting partial memorization rather than purely generalizable learning. Fine-tuning partially ameliorates but does not eliminate this effect.

A key application we highlight is structure-based vaccine design. On the limited available data comprising 33 verified antigen-stabilizing mutations across five viruses, HERMES-*amortized* performs well at recovering these mutations. Building on this, we developed two design pipelines for antigen-stabilizing mutation libraries that use HERMES-*amortized* to rapidly score all possible mutations across an antigen, then filter the top candidates with Rosetta. The model’s strict locality makes it straightforward and fast to apply to mutational effects driven by local, short-range interactions in large antigens and assemblies, which can be challenging for architectures that model global interactions.

Our results show that local all-atom environments modeled by HERMES can encode multiple stabilization mechanisms, while also underscoring limitations for mechanisms that depend on long-range coupling, or multi-site epistasis. Interestingly, stability fine-tuning does not consistently transfer to antigen stabilization and can even degrade performance. One plausible explanation is dataset mismatch: the high-throughput stability datasets used for fine-tuning [20] are enriched for small domains with relatively few buried residues, whereas vaccine antigens are often larger and multi-domain, with a greater proportion of stabilizing mutations occurring at buried sites. Closing this gap will likely require task-aligned fine-tuning data or broader supervision that better reflects antigen structure and design objectives.

We further evaluated HERMES on predicting changes in protein-protein binding affinity. HERMES shows measurable zero-shot predictive signal—particularly at the interface—and supervised fine-tuning substantially improves performance.

Beyond the size-dependent and wild-type biases that relaxations and fine-tuning ameliorate, HERMES’s principal limitation is its strict locality: by construction, its predictions depend only on the focal residue’s atomic neighborhood within a 10 Å radius. HERMES can therefore capture only mutational effects driven by short-range interactions and local packing, as evidenced by its failure to predict binding-affinity changes for mutations outside the protein-protein interface. This locality is a double-edged sword: it restricts the operational domain of HERMES, but also reduces input complexity, removes constraints on protein size, and improves scalability. We therefore propose the HERMES family as a suite of well-characterized models suited to fast screening of mutations that stabilize (or destabilize) protein tertiary and quaternary structures via local packing.

Several directions could extend HERMES’s capabilities. On the locality front, future work could better characterize when locality suffices and develop multi-scale approaches that combine local all-atom predictors with complementary global or multi-site models. More broadly, larger and more diverse fine-tuning datasets—particularly for antigen stabilization and binding—would likely improve performance, since our results indicate that training data dominates architectural differences in determining benchmark performance.

## Acknowledgement

This work has been supported by the National Institute of General Medical Sciences (Award Number R35 GM142795) and the National Institute of Allergy and Infectious Diseases (Award Number U19AI181881) of the National Institutes of Health. The content is solely the responsibility of the authors and does not necessarily represent the official views of the National Institutes of Health. This work has also been supported by the CAREER award from the National Science Foundation grant 2045054, the Royalty Research Fund from the University of Washington no. A153352, the Allen School Computer Science & Engineering Research Fellowship from the Paul G. Allen School of Computer Science & Engineering at the University of Washington, and the Microsoft Azure award from the eScience institute at the University of Washington. This work is also supported, in part, through the Departments of Physics and Computer Science and Engineering, and the College of Arts and Sciences at the University of Washington. U.W. acknowledges the REU program at the Department of Physics at the University of Washington, which is supported by the National Science Foundation grant 2243362. The numerical analyses in this work were completed on Hyak, the University of Washington’s high performance computing cluster, which is funded by the University of Washington student technology fee.

## Methods

### HERMES architecture

The HERMES architecture closely follows our recent developments of SO(3)-equivariant neural network models for protein structures [26, 36, 37]. For completeness, we summarize here the main methodological components and refer the reader to those works for additional details.

The input to HERMES is a point cloud of atoms in a protein structure, which we term a *neighborhood*. Each neighborhood is centered at the C-*α* atom of a focal residue and includes all atoms within a 10 Å radius of this center. Optionally, we add Gaussian noise with standard deviation 0.50 Å to the atomic coordinates to achieve further model robustness. The output of HERMES is a 20-dimensional representation of the neighborhood that is invariant to 3D rotations about its center (i.e., SO(3)-invariant). To obtain SO(3) invariance, HERMES is constructed from SO(3)-equivariant layers that progressively compute higher-level SO(3)-invariant features, which are then passed to a final multilayer perceptron (MLP) with a 20-dimensional output. Conceptually, symmetry awareness in HERMES is achieved through two main components: (i) a (holographic) encoding of the neighborhood into a basis suitable for SO(3)-equivariant operations, and (ii) processing of the input via a stack of SO(3)-equivariant neural network layers to learn an expressive and SO(3)-invariant representation of the neighborhood.

#### Holographic encoding of atomic protein structure neighborhoods

We first represent the atomic point cloud of each structural neighborhood as a density function obtained by superposing (weighted) Dirac-δ functions, indicating the presence of atoms at a given position in space: *ρ*(**r**) =∑ _*i*∈points_ *ω*_*i*_δ(**r**_*i*_ − ***r***; here, *ω*_*i*_ indicates the weight associated with point *i* at position **r**_*i*_, and **r**_*i*_ can be decomposed in its constituents spherical components (*r*_*i*_, *θ*_*i*_, *ϕ*_*i*_). We then use 3D Zernike Fourier Transform (ZFT) [26] of the density function to encode the neighborhood into a convenient SO(3) equivariant basis,

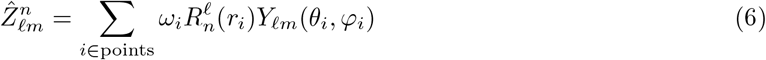

where *Y*_*m*_(*θ*, φ) is the spherical harmonics of integer degree *ℓ* ≥ 0 and integer order *m* ≤ |*ℓ*|, and *R*^*n*^(*r*) is the radial Zernike polynomial in 3D with integer radial frequency *n* ≥ 0 and degree *ℓ*. 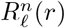 is non-zero only when *n* − *ℓ* is even and ≥ 0. We keep coefficients of up to and including ℓ = 5, and, for every ℓ, we keep the first 11 non-zero radial frequencies.

Notably, the spherical harmonics that describe the angular component of ZFT arise from the irreducible representations of the 3D rotation group SO(3), and form a convenient basis under rotation in 3D (see the Appendix of [37] for a formal mathematical introduction). Zernike projections in spherical Fourier space can be understood as a superposition of spherical holograms of an input point cloud, and thus, we term this operation an *holographic encoding* of the data [26, 36]. The resulting holograms are primarily indexed by the degree ℓ: different values of ℓ encode components that transform in specific ways under 3D rotations. For example, the ℓ = 0 component is rotation-invariant.

Following [26, 36], we incorporate atom-level features by partitioning the holographic encoding into multiple *channels* (Fig. 1A). Specifically, we use separate channels for C, N, O, S, computationally added hydrogens, partial charge, and solvent-accessible surface area (SASA). Partial charge and SASA are defined for all atoms, and their values are incorporated through the weights *ω*_*i*_.

#### SO(3)-equivariant neural network architecture

The neighborhood holograms are then processed by a stack of SO(3)-equivariant layers (Fig. 1A). The final SO(3)-invariant representation is obtained by reading out the *f* = 0 component of the last layer and passing it through an MLP to produce a 20-dimensional embedding. We employ three types of SO(3)-equivariant building blocks, inspired by the H-CNN architecture [26] and also detailed in ref. [37]: (i) SO(3)-equivariant linear layers (Lin), which mix channels while preserving the SO(3) transformation properties; (ii) tensor-product nonlinearities (TP), which couple different irreducible components; and (iii) SO(3)-equivariant layer normalization (LN).

All SO(3)-equivariant components are implemented using `e3nn` primitives [58]. Within the same framework, we re-implemented the H-CNN architecture described in [26] for comparison. HERMES achieves a forward pass that is approximately 2.75*×* faster than H-CNN, while using a similar number of parameters (~ 3.5M).

### Pre-processing of protein structure data

To pre-process protein structure data, we developed two distinct pipelines based on either (i) PyRosetta [34] or (ii) Biopython [59] together with other open-source tools, with code adapted from [2]. The PyRosetta-based pipeline is considerably faster but requires a license, whereas the Biopython-based pipeline is fully open source. We train models using data generated by both pipelines, with the constraint that the Pre-processing pipeline used at inference must match that used during training. Performance differences between the two pipelines are minor (Fig. S13 and S27, and Table S1); unless otherwise stated, we report results obtained with the PyRosetta pipeline, which yields slightly better performance and has faster Pre-processing runtime.

For the PyRosetta workflow, we use PyRosetta functionalities for all Pre-processing steps: repairing PDB files by adding missing residues, adding hydrogen atoms, assigning partial atomic charges, computing solvent-accessible surface areas (SASA); we ignore non-canonical amino acids. For the open-source Pre-processing workflow (“Biopython” pipeline), we proceed as follows. First, we use OpenMM [60] to repair PDB files by adding missing residues and substituting non-canonical residues with their canonical counterparts. Hydrogen atoms are then added using the `reduce` program [61]. Partial atomic charges are assigned from the AMBER99sb force field [62], and SASA are computed using Biopython [59].

Both pre-processing pipelines retain atoms belonging to non-protein residues and ions, in contrast to the RaSP pre-processing procedure [2]. Notably, the PyRosetta-based pipeline does not replace non-canonical residues.

### HERMES pre-training

We pre-trained HERMES using an inverse folding objective, in which the model predicts the identity of a masked focal amino acid (i.e., the native residue in the structure) given its surrounding atomic neighborhood. This training task is analogous to that used for H-CNN [26]. Like H-CNN, HERMES is trained on data derived from ProteinNet’s [38] 30% sequence-identity clustering of PDB structures available at the time of CASP12. Specifically, we used ProteinNet’s “training 30” set randomly split in 12,842 training and 1,026 validation PDB structures, and ProteinNet’s “validation” set of 213 PDB structures for testing. Importantly, ProteinNet’s clustering contains only some chains within the structures, deduplicated at 30% sequence identity, and thus the number of unique chains is smaller than the total number of chains in all the structures.

Model parameters were optimized for 10 epochs using the Adam optimizer [63] with a learning rate of 10^−3^. We selected the model checkpoint with the lowest validation loss at the end of each epoch. A single HERMES model is implemented as an ensemble of 10 independently trained neural network instances, whose predictions are averaged at inference time. Pre-training a single network instance required approximately 40 minutes per epoch on a single NVIDIA A40 GPU.

### Rosetta relaxation of mutant structures for the HERMES-*relaxed* protocol

For the HERMES-*relaxed* protocol, we use PyRosetta [34] to generate and locally refine mutant protein structures. Specifically, the focal residue is first substituted with the desired mutant amino acid, after which all side-chain atoms within 12 Å of the focal residue’s C-*α* atom are relaxed using the FastRelax protocol with the `ref2015_cart` energy function and a single relaxation cycle.

We performed a targeted ablation study to select the relaxation parameters. These experiments indicate that allowing backbone atoms to relax slightly degrades performance, and that ensembling over multiple independent relaxation runs does not improve accuracy (Fig. S1A), while substantially increasing computational cost (Fig. S1B). Thus, all results for the HERMES-*relaxed* protocol are reported using side-chain-only relaxation with a single FastRelax cycle.

### Training HERMES-*amortized* to implicitly learn about relaxation

We fine-tuned HERMES to align HERMES-*fixed* predictions to those of HERMES-*relaxed*, on a subset of the pre-training protein sites. Specifically, we uniformly sampled 10% of the proteins pre-training training set (1,284), and from those we uniformly sampled 5% of the sites. We similarly sampled 1% of the sites from the proteins of the pre-training validation set, and 5% of the pre-training testing set.

### HERMES fine-tuning for downstream tasks

In all analyses, we fine-tuned HERMES models under a Huber Loss objective with hyperparameter δ = 1.0, for 15 epochs using the Adam optimizer [63] with a learning rate of 10^−3^ and a batch size of 128 mutations, selecting the model checkpoint with the lowest validation loss at the end of each epoch. The only exception is the models without pre-training, which are trained for 25 epochs.

By convention, we define model outputs such that higher predicted values correspond to more favorable mutation effects. Accordingly, when fine-tuning on experimental ΔΔ*G* measurements–where lower values indicate greater stability–we trained the model to predict − ΔΔ*G*. This sign convention ensures consistent interpretation of model outputs across tasks and is fixed throughout all reported analyses.

To speed-up convergence during fine-tuning, we first rescale the weight matrix and bias vector of the network’s output layer so that the mean and variance of the output logits match those of the validation scores. This initialization step requires a forward pass through the validation data to estimate the mean and variance, but it makes the model outputs immediately on the same scale as the experimental scores. Thus, fine-tuning avoids spending early epochs merely adjusting output magnitudes.

Overall, fine-tuning is computationally efficient. Training a single neural network instance requires approximately 2.5 minutes per epoch on the cDNA117k dataset (~ 117,000 mutations) and about 4 minutes per epoch on the Megascale training dataset (~ 217,000 mutations) on a single NVIDIA A40 GPU.

### Fine-tuning datasets

#### Stability effect prediction

For protein stability prediction, we fine-tuned and evaluated the performance of HERMES on the same datasets of three recent structure-based predictors of mutational stability effects–RaSP [2], Stability-Oracle [27], and ThermoMPNN [30]–so results are comparable. The data from these models are used in following way:

##### RaSP dataset

We used the RaSP dataset [2] as provided by the authors on their github repository (https://github.com/KULL-Centre/_2022_ML-ddG-Blaabjerg). As in the original RaSP paper, the target values are Rosetta-computed stability changes (ΔΔ*G*), which are known to be reliable primarily within the range [-7, 1] kcal/mol. To account for this, RaSP applies a sigmoid transformation to the raw ΔΔ*G* values prior to training, effectively saturating the targets outside this interval.

We adopt the same sigmoid transformation, with one modification: we center the transformed values such that ΔΔ*G* = 0 maps to zero after transformation. This centering is required by the HERMES output parameterization, in which the predicted stability change for a mutation to the same amino acid is constrained to be zero (i.e.,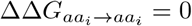). While this property holds for physical ΔΔ*G* values, it is not preserved by the uncentered sigmoid transform used in RaSP. Our centered transformation therefore ensures consistency between the model’s output space and the physical interpretation of stability changes, and it follows,

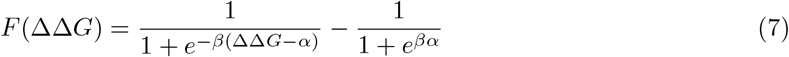

##### Stability-Oracle datasets

Stability-Oracle introduces two curated datasets that we use in this work [27]:

i. cDNA117k (training set): derived from the Megascale cDNA display proteolysis dataset # 1 [20]. The original Megascale dataset reports approximately 850,000 thermodynamic folding stability measurements (Δ*G*) across 354 natural and 188 de novo mini-protein domains (40-72 amino acids in length). Following the Stability-Oracle protocol, mutations are filtered to enforce at most 30% sequence identity with the test set, yielding a reduced set of approximately 117,000 mutation-induced stability changes (ΔΔ*G*), referred to as cDNA117k.
ii. T2837 (test set): assembled by combining several commonly used benchmarking datasets of experimentally measured ΔΔ*G* values, including S669 [43], myoglobin [28], ssym [64], and p53 [65]. The resulting test set comprises 2,837 mutations. We obtained the cDNA117k and T2837 datasets from the Stability-Oracle [27] github repository (https://github.com/danny305/StabilityOracle/tree/master). At the time of access, the residue indices provided in the dataset did not correspond to the residue numbering in the original PDB files, but instead to an intermediate, post-processed representation that was not documented in sufficient detail to allow straightforward recovery of the original numbering. To ensure consistency with structural data, we manually corrected the residue numbers in the CSV files to match those in the corresponding PDB structures. The corrected versions of these datasets are included in our repository.

##### ThermoMPNN dataset

ThermoMPNN also leveraged Megascale [20], but used datasets # 2 and # 3, curated and split by the authors at a 25% sequence-identity cutoff into 216k training and 28k test mutation effects [30]. We used the train, valid, and test splits of the Megascale dataset as curated in ThermoMPNN, and provided on their github repository (https://github.com/Kuhlman-Lab/ThermoMPNN). Throughout the text, we refer to these datasets as the Megascale training (train + valid) and testing sets. We downloaded the structures’ `.pdb` files from https://zenodo.org/records/7992926. We note that the Megascale training set was not controlled for maximum similarity with the T2837 dataset, and six of the T2837 proteins (~ 5%) have sequence homologs in the Megascale training set with sequence similarity above 90%.

#### Binding effect prediction

For predicting mutational effects on binding affinity, we fine-tuned HERMES on the SKEMPI v2.0 dataset [55], using 3-fold cross-validation.

##### SKEMPI dataset

After removing duplicate experimental entries, SKEMPI v2.0 contains 5,713 measurements of binding free-energy changes (ΔΔ*G*^binding^) spanning 331 protein-protein complex structures. We further restrict the dataset to complexes with at least 10 annotated mutations, resulting in 116 structures and 5,025 mutations. Finally, we retain only single-point mutations, yielding 93 structures and 3,485 mutations.

SKEMPI provides metadata explicitly designed to support leakage-aware evaluation via two fields: *hold-out type* and *hold-out proteins* (see SKEMPI documentation: https://life.bsc.es/pid/skempi2/info/faq_and_help). Briefly, *hold-out type* groups complexes into broad categories (e.g., protease-inhibitor, antibody-antigen, and pMHC-TCR), while *hold-out proteins* lists PDB identifiers and/or hold-out types that should be co-held-out to avoid training on closely related binding sites. Using this information, we define three split strategies:

i. *Easy:* random splitting without using hold-out metadata.
ii. *Medium:* we enforce that all entries linked via a mutation’s *hold-out proteins* annotation are assigned to the same split (but we do not additionally group by *hold-out type*).
iii. *Difficult:* we enforce that all complexes sharing the same *hold-out type* are assigned to the same split, producing a stringent evaluation of generalization across interaction classes.

In cases where a complex is associated with multiple hold-out types, we assign it to a single type by randomly selecting one of the available labels.

### Baseline models

Here, we describe the baseline models used to benchmark HERMES on mutational effect prediction.

#### ProteinMPNN [35]

ProteinMPNN is an inverse-folding model that samples amino acid sequences conditioned on a protein backbone (optionally with a partial sequence fixed). Because it also outputs per-site amino acid probabilities, we used it to score mutational effects via the log-likelihood ratio in Eq. 1. As for HERMES, we evaluate ProteinMPNN models trained with two noise levels: 0.02 Å(virtually no noise) and 0.30 Å. Unlike HERMES, however, predictions of ProteinMPNN are not an ensembled across models. We used, and provide, scripts to infer mutation effects built upon a public fork of the ProteinMPNN repository (https://github.com/gvisani/ProteinMPNN-copy).

#### ThermoMPNN [30]

ThermoMPNN is a thermodynamic stability predictor built on top of Protein-MPNN. For a given structure, it extracts the final ProteinMPNN residue embedding at the site of interest and feeds it to a separate head that predicts per amino acid Δ*G* values, from which ΔΔ*G* is computed. Similar to HERMES, this formulation enforces the permutation symmetry of mutational effects by construction, without requiring data augmentation. For our experiments, we used native functionalities in the ThermoMPNN repository (https://github.com/Kuhlman-Lab/ThermoMPNN).

#### Stability-Oracle [27]

Similar to HERMES, Stability-Oracle is trained in two stages. First, a graph-attention network is pre-trained to predict masked amino acids from their local atomic environment (“neighborhood”). Next, the embeddings from the pre-trained model is used to regress over mutation-effect. Specifically, for a target site on a structure, the masked-neighborhood embedding *h* is extracted from the pre-trained network and concatenated separately with embeddings of the “from” and “to” amino acids. Each concatenated input is passed through a transformer to produce amino acid specific embeddings 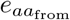 and 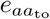, whose difference 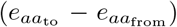 is fed to a two-layer MLP to predict 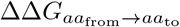. This construction is permutation-symmetric up to the final MLP, since each *e*_*aa*_ is computed independently; symmetry is broken only by the MLP, and would have been preserved by a bias-free linear layer. The original method enforces symmetry during training via 19 *×* data augmentation. To our knowledge, the method does not employ ensembling. We report performance scores calculated using predictions provided in the Stability-Oracale’s repository (https://github.com/danny305/StabilityOracle).

#### RaSP [2]

Similar to HERMES, RaSP is trained in two steps. First, a 3D CNN is pre-trained to predict masked amino acids from their local atomic environment (“neighborhood”). Then, a small fully-connected neural network with a single output is trained to regress over mutation effects, using as input neighborhoods’ embeddings from the 3DCNN, the one-hot encodings of wildtype and mutant amino acids, and the wildtype and mutant amino acids’ frequencies in the pre-training data. RaSP is fine-tuned on the stability effect of mutations ΔΔ*G*, computationally determined with Rosetta [34]. We report performance scores calculated using predictions provided in the authors’ repository (https://github.com/KULL-Centre/_2022_ML-ddG-Blaabjerg).

#### RDE-Network [33]

RDE stands for Rotamer Density Estimator. This model consists of first a graph neural network encoder that is trained via a normalizing flow objective to predict distributions of side-chain conformations. Then, a prediction head is added, and it is trained on ΔΔ*G*_binding_ effects from the SKEMPI dataset. We report performance scores as provided in ref. [33].

#### MIF-Network [33]

This model’s architecture is the same as the encoder in RDE-Network. It is first pre-trained on the task of wild-type amino acid classification. Then, a prediction head is added to the encoder, and it is trained on ΔΔ*G*_binding_ effects from the SKEMPI dataset [55] in the same manner as RDE-Network. We report performance scores as provided in Ref. [33].

#### Pythia-PPI [54]

Pythia-PPI uses a graph neural network encoder pre-trained for wild-type amino acid classification, followed by a ΔΔ*G* prediction module with two heads: one for ΔΔ*G*_stability_ and one for ΔΔ*G*_binding_. The model is fine-tuned jointly on stability labels from FireProtDB [56] and binding labels from SKEMPI [55], using a validation selected 20:80 stability:binding mixing ratio. The resulting checkpoint (“Vanilla Pythia-PPI”) is then further trained via self-distillation by fine-tuning on its own predictions over all SKEMPI complex structures to obtain the final model (Pythia-PPI). We should note that these two procedures (i.e., joint fine-tuning on stability and binding with a validation selected mixing ratio and self-distillation) could also be applied to HERMES to potentially improve performance. We report performance scores as provided in ref. [54].

### ESMFold for computational modeling of protein structures

For the analysis in Fig. S15, we used the ESM Metagenomic Atlas API to fold each sequence individually (https://esmatlas.com/resources?action=fold).

### Sequence similarity calculation between training proteins and test proteins

In Figures 5, S9, S10, S11 and S12, results are stratified by maximum similarity to relevant pre-training proteins. In Table S7, we report the sequence similarity between each wild-type antigen and the pre-training proteins. To compute protein-protein sequence similarity, we used BLASTp via NCBI BLAST+ [66] with individual chains in the test set as queries, and individual chains in the training set as the database. For each query chain, we considered the maximum similarity to any chain in the database, where similarity is computed as length of sequence overlap divided by the length of the query chain. We used the length of the query, rather than the more common minimum length between query and hit, as it better captures how many predictions - which occur at individual query *residues* - are likely to have been influenced by pre-training memorization. We only considered sequence alignments with coverage (for both query and target) over 30%.

### Structure-conditioned reversibility analysis

In Fig. 5 we characterized structure-conditioned reversibility on the SSym dataset [64], which contains wild-type and single-mutant structures for 352 mutations across 19 proteins [43].

We assess structure-conditioned reversibility by whether the effects of forward *α* → *β* and reverse *β*→ *α* mutations are equal, using the outgoing structural context to compute mutational effects. Specifically, the forward mutational effect 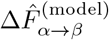 is computed by conditioning on the structure *X*_*α*_, and the reverse effect 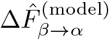 is computed using *X*_*β*_,

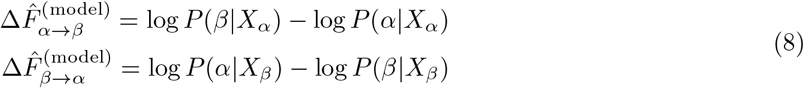

With this definition, we do not expect reversibility, as forward and reverse transitions are conditioned on different structures.

#### Reversibility score

For the analysis in Fig. 5 we construct a reversibility score as a mean squared error normalized to be between −1 and 1,

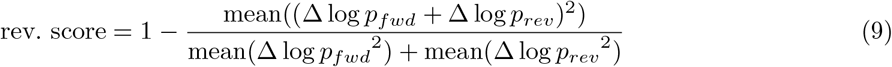

where Δ log *p*_*fwd*_ = log *p*(mt|*X*_wt_)−log *p*(wt|*X*_wt_) is the forward mutational effect computed by conditioning on the wild type structure, and Δ log *p*_*rev*_ = log *p*(wt |*X*_mt_) − log *p*(mt| *X*_mt_) is the reverse mutational effect computed by conditioning on the mutants structure. The averages (mean) are computed over the 352 mutations in the SSym dataset, or stratified as needed by the desired criteria. With this definition, a larger value of **rev. score** implies more degree of reversibility between forward and reverse mutations.

#### Sequence similarity calculation between the Ssym dataset and CASP12 pre-training proteins

Using BLASTP as mentioned in the previous section, we found that 9 Ssym proteins have similarity below 70% to the pre-training proteins (only 5 are below 50%, none are below 40%), and 10 proteins have similarity above 70% (comprising the two panels in Fig. 5B).

### Statistical comparison of model performances on classification metrics

#### Comparing models’ performances on the same dataset

Figures 2 and 3 report classification performance (precision, recall, F1, etc.) for predicting the stability effects of mutations across models. Figures S4 and S7report pairwise significance tests (p-values) for differences in these metrics using permutation tests, with the null hypothesis that two models’ predictions are exchangeable. To compute these p-values, for each model pair, we generated permuted prediction sets by swapping paired prediction between models with probability 0.5, and repeat this procedure for 1000 random seeds. The p-value is the fraction of permutations in which the metric difference between the permuted sets is at least as large (≥) as the observed difference. We correct the computed p-values for multiple comparisons using Holm-Bonferroni across all model pairs within each metric (i.e., within each panel in Figs. S4 and S7).

#### Comparing a model’s performance across different datasets

Figures 3 and S8 respectively report classification performance for predicting mutation stability effects across mutation size categories, and pairwise significance tests for differences in these metrics between mutation size categories. Similarly, Figure S10, S11 and S12 report correlation and classification performance across subsets of test proteins with varying maximum similarity to the pre-training proteins.

In both cases, we estimate p-values by bootstrap resampling. For each dataset pair, we draw 1000 bootstrap samples of the predictions within each dataset, recompute the metric difference on each resample, and define the p-value as the fraction of replicates in which the resampled difference is at least as large as the observed one. In the first case, we resample mutations directly. In the second, we resample proteins and then adjust the total number of mutations to match the original dataset size by randomly dropping mutations or duplicating them as needed. In both cases, we apply a Holm–Bonferroni correction across all dataset pairs considered for a given metric.

#### Retrospective recall analysis of antigen-stabilizing mutations

We manually collected 33 previously reported antigen-stabilizing mutations across five viral antigens with reliable pre-fusion structures in the PDB: Influenza HA [12] (3), RSV-F [9, 67] (7), hMPV-F [10, 11, 13] (11), DENV-E [13] (8), and SARS-CoV-2 spike [68] (4 mutations).

In Figures 6, S20 and Table S4, we report, for each model, the number of antigen stabilizing mutations retrieved (*x*) out of a total of *N* = 33 experimentally verified such mutations. Retrieval is based on amino acid ranking: a mutation is counted as retrieved if (i) the mutant amino acid is ranked better than the wild type (*r*_mt_ *< r*_wt_), and (ii) that its rank is below or equal to a certain threshold *R*, with *R* = 3 for “strongly suggested” and *R* = 6 for “moderately suggested.” These thresholds correspond to per-site top-3 and top-6 recall, respectively; a mutation with *r*_mt_ *< r*_wt_ but *r*_mt_ *>* 6 is instead “weakly suggested,” even though it has a better rank than the wild-type. Ranks are taken among the 20 amino acids at each position and computed conditioning on the pre-fusion structure, effectively predicting the effects on absolute pre-fusion stability. Because this scheme groups physico-chemically similar mutations into the same rank class, models are generally not penalized for ranking a biophysically similar alternative above a known stabilizing mutant (see Fig. S21 and SI for a more detailed discussion). Moreover, this scheme is insensitive to differences in score (logit) scale across models (Fig. S21), making cross-model comparisons more robust.

We assess significance in the number of retrieved antigen-stabilizing mutations by a given model with a one-sided binomial test: for each model, the p-value is the probability under a null model of recovering at least *x* successes in *N* trials with per-mutation success probability *p*_null_, i.e., *p* = 1 − BinomCDF(*x* − 1, *N, p*_null_). We adjust p-values for multiple testing using Holm-Bonferroni, and report the resulting p-values in Fig. S24.

To compute the p-values, we consider two null models:

i. **Random null**. For each mutation, we sample ranks for the mutant and wild type uniformly without replacement: *r*_mt_ ~ Unif*{*1, …, 20*}* and *r*_wt_ ~ Unif{1, …, 20} *\ r*_mt_. A mutation is a “success” if *r*_mt_ *< r*_wt_ and *r*_mt_ ≤ *R*, which occurs with probability 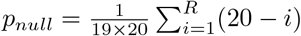.
ii. **BLOSUM62 null**. We simply set *p*_*null*_ = *x*_*B*62_*/N*, where *x*_*B*62_ is the number of mutations with BLOSUM62 rank 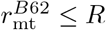. We note that the rank of the wild-type is always 1 for BLOSUM62, so we omit the condition that the rank of the mutant has to be lower than the rank of the wild-type.

### Identifying neighboring mutations to stratify antigen-stabilizing mutations

The stabilized antigens considered in this study contain multiple mutations that, in most cases, have not been evaluated individually for their stabilization potential. For each mutation in a given stabilized antigen, we determine whether it has at least one “close neighbor” among the antigen’s other mutations, based on the distance between their residues in the wild-type structure. Specifically, we define residue Y as a close neighbor of residue X if any atom of Y lies within 10 Å of the C*α* of residue X. Under this definition, atoms of close by residue Y—and hence its amino acid identity—are part of HERMES’s input when predicting the effect of a mutation at residue X.

### Using ProteinMPNN to score antigen-stabilizing mutations

When scoring homo-oligomeric antigens with ProteinMPNN, care must be taken to avoid information leakage between equivalent positions on different chains. Specifically, when scoring a mutation at a given site, we masked the wild-type identity at that site on every homomeric chain simultaneously and predicted log-probabilities with logits tied across those positions.

### Using Rosetta to score antigen-stabilizing mutations

In Figures 6, S19, S20, S21and Tables S4, S5, we reported the Rosetta scores for the verified antigen-stabilizing mutations. In Figures 7 and S25 we used Rosetta scores as oracles for mutations suggested by other models. To compute these scores, we used the PyRosetta software [34] to model protein structures, with wild-type structures serving as template. For each target sequence containing a single point mutation, we threaded the mutant sequence onto all chains of the oligomeric template. Each resulting model was then subjected to a high-resolution refinement protocol using Rosetta’s FastRelax application. This protocol involved five cycles of side-chain rotamer repacking followed by gradient-based energy minimization of backbone (φ, *ψ*) and side-chain (*χ*) torsion angles, guided by the `ref2015_cart` all-atom energy function. To improve conformational sampling, we repeated the threading-and-relaxation procedure 20 times per mutation for use when analyzing validated mutations, and 50 when used as an oracle. For each mutant, we report the mean Rosetta Energy Unit (REU) score of the 20% lowest-energy (most favorable) relaxed models (5 for the analysis against validated mutations, and 13 when used as an oracle).

### Design of screening libraries to identify antigen-stabilizing mutations

Using HERMES-*amortized*, we devised two pipelines to identify candidate antigen-stabilizing mutations.

1. *Stabilize pre-fusion core:* Given a pre-fusion structure, we scored all possible mutations at core sites (average SASA *<* 1 Å ^2^) by Δ log *p*^pre-f^, where higher scores indicate stronger predicted stabilization. We then took the top 80 mutations, all predicted stabilizing (Δ log *p*^pre-f^ *>* 0) and capped at 3 per site to encourage site variability. This pipeline ignores post-fusion effects.
2. *Stabilize pre-fusion and destabilize post-fusion:* We took the top 80 mutations (again capped at 3 per site, but with no location constraint) ranked by Δ log *p*^pre-f^ − Δ log *p*^post-f^, requiring both Δ log *p*^pre-f^ *>* 0 (stabilizing) and Δ log *p*^post-f^ *<* 0 (destabilizing).

As baselines, we sampled mutations with ProteinMPNN, BLOSUM62, and at random. ProteinMPNN mutations were selected exactly as with HERMES (80 per pipeline). For BLOSUM62, we randomly selected 80 sites per pipeline (pre-fusion core only, or unrestricted) and took the top two mutations at each site by score. Random baselines drew 160 mutations under the same site constraints (max 3 per site; core only or unrestricted). Across all models and pipelines, we excluded mutations to and from cysteine, both to avoid breaking disulfide bonds and because stabilizing cysteine mutations typically come in pairs to form new ones.

To validate the suggestions, we computed Rosetta scores as for the verified antigen-stabilizing mutations (see above), but with a larger sample size. Each mutation receives an energy score *E*_ros_ in Rosetta Energy Units (REU, lower is better) and is counted as a “Rosetta hit” if 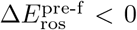 (pipeline 1), or both 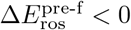 and 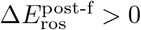 (pipeline 2).

## References

[1] Gerasimavicius, L., Liu, X. & Marsh, J. A. Identification of pathogenic missense mutations using protein stability predictors. Scientific Reports 10, 15387 (2020). URL https://www.nature.com/articles/s41598-020-72404-w.

[2] Blaabjerg, L. M. et al. Rapid protein stability prediction using deep learning representations. eLife 12, e82593 (2023). URL 10.7554/eLife.82593.

[3] Ishida, T. Effects of Point Mutation on Enzymatic Activity: Correlation between Protein Electronic Structure and Motion in Chorismate Mutase Reaction. Journal of the American Chemical Society 132, 7104–7118 (2010). URL 10.1021/ja100744h.

[4] Wang, X. et al. D3DistalMutation: a Database to Explore the Effect of Distal Mutations on Enzyme Activity. Journal of Chemical Information and Modeling 61, 2499–2508 (2021). URL 10.1021/acs.jcim.1c00318.

[5] Thadani, N. N. et al. Learning from prepandemic data to forecast viral escape. Nature 622, 818–825 (2023). URL https://www.nature.com/articles/s41586-023-06617-0.

[6] Luksza, M. & Lässig, M. A predictive fitness model for influenza. Nature 507, 57–61 (2014).

[7] Neher, R. A., Russell, C. A. & Shraiman, B. I. Predicting evolution from the shape of genealogical trees. Elife 3 (2014).

[8] Hie, B. L. et al. Efficient evolution of human antibodies from general protein language models. Nature Biotechnology 42, 275–283 (2024). URL https://www.nature.com/articles/s41587-023-01763-2.

[9] McLellan, J. S. et al. Structure-Based Design of a Fusion Glycoprotein Vaccine for Respiratory Syncytial Virus. Science 342, 592–598 (2013). URL https://www.science.org/doi/10.1126/science.1243283.

[10] Gonzalez, K. J. et al. A general computational design strategy for stabilizing viral class I fusion proteins. Nature Communications 15, 1335 (2024). URL https://www.nature.com/articles/s41467-024-45480-z.

[11] Bakkers, M. J. G. et al. Efficacious human metapneumovirus vaccine based on AI-guided engineering of a closed prefusion trimer. Nature Communications 15, 6270 (2024). URL https://www.nature.com/articles/s41467-024-50659-5.

[12] Milder, F. J. et al. Universal stabilization of the influenza hemagglutinin by structure-based redesign of the pH switch regions. Proceedings of the National Academy of Sciences 119, e2115379119 (2022). URL https://www.pnas.org/doi/full/10.1073/pnas.2115379119.

[13] Phan, T. T. N. et al. A conserved set of mutations for stabilizing soluble envelope protein dimers from dengue and Zika viruses to advance the development of subunit vaccines. Journal of Biological Chemistry 298, 102079 (2022). URL https://www.sciencedirect.com/science/article/pii/S0021925822005191.

[14] Rocklin, G. J. et al. Global analysis of protein folding using massively parallel design, synthesis, and testing. Science 357, 168–175 (2017). URL https://www.science.org/doi/10.1126/science.aan0693.

[15] Lindorff-Larsen, K. & Teilum, K. Linking thermodynamics and measurements of protein stability. Protein Engineering, Design and Selection 34, gzab002 (2021). URL 10.1093/protein/gzab002.

[16] Karlsson, R. SPR for molecular interaction analysis: a review of emerging application areas. Journal of Molecular Recognition 17, 151–161 (2004). URL https://onlinelibrary.wiley.com/doi/abs/10.1002/jmr.660. eprint: https://onlinelibrary.wiley.com/doi/pdf/10.1002/jmr.660.

[17] Fowler, D. M. & Fields, S. Deep mutational scanning: a new style of protein science. Nature Methods 11, 801–807 (2014). URL https://www.nature.com/articles/nmeth.3027.

[18] Kinney, J. B. & McCandlish, D. M. Massively parallel assays and quantitative Sequence–Function relationships. Annu. Rev. Genomics Hum. Genet. 20, 99–127 (2019).

[19] Starr, T. N. et al. Deep Mutational Scanning of SARS-CoV-2 Receptor Binding Domain Reveals Constraints on Folding and ACE2 Binding. Cell 182, 1295–1310.e20 (2020). URL https://www.ncbi.nlm.nih.gov/pmc/articles/PMC7418704/.

[20] Tsuboyama, K. et al. Mega-scale experimental analysis of protein folding stability in biology and design. Nature 620, 434–444 (2023). URL https://www.nature.com/articles/s41586-023-06328-6.

[21] Gapsys, V., Michielssens, S., Seeliger, D. & de Groot, B. L. Accurate and Rigorous Prediction of the Changes in Protein Free Energies in a Large-Scale Mutation Scan. Angewandte Chemie International Edition 55, 7364–7368 (2016). URL https://onlinelibrary.wiley.com/doi/abs/10.1002/anie.201510054. xeprint: https://onlinelibrary.wiley.com/doi/pdf/10.1002/anie.201510054.

[22] Schymkowitz, J. et al. The FoldX web server: an online force field. Nucleic Acids Research 33, W382–W388 (2005). URL 10.1093/nar/gki387.

[23] Kellogg, E. H., Leaver-Fay, A. & Baker, D. Role of conformational sampling in computing mutation-induced changes in protein structure and stability. Proteins: Structure, Function, and Bioinformatics 79, 830–838 (2011). URL https://onlinelibrary.wiley.com/doi/abs/10.1002/prot.22921. eprint:https://onlinelibrary.wiley.com/doi/pdf/10.1002/prot.22921.

[24] Riesselman, A. J., Ingraham, J. B. & Marks, D. S. Deep generative models of genetic variation capture the effects of mutations. Nature Methods 15, 816–822 (2018). URL https://www.nature.com/articles/s41592-018-0138-4.

[25] Meier, J. et al. Language models enable zero-shot prediction of the effects of mutations on protein function. Advances in Neural Information Processing Systems, Vol. 34, 29287–29303 (Curran Associates, Inc., 2021). URL https://proceedings.neurips.cc/paper/2021/hash/f51338d736f95dd42427296047067694-Abstract.html.

[26] Pun, M. N. et al. Learning the shape of protein microenvironments with a holographic convolutional neural network. Proceedings of the National Academy of Sciences 121 (2024). URL https://www.pnas.org/doi/10.1073/pnas.2300838121.

[27] Diaz, D. J. et al. Stability Oracle: a structure-based graph-transformer framework for identifying stabilizing mutations. Nature Communications 15, 6170 (2024). URL https://www.nature.com/articles/s41467-024-49780-2.

[28] Li, B., Yang, Y. T., Capra, J. A. & Gerstein, M. B. Predicting changes in protein thermodynamic stability upon point mutation with deep 3D convolutional neural networks. PLOS Computational Biology 16, e1008291 (2020). URL https://journals.plos.org/ploscompbiol/article?id=10.1371/journal.pcbi.1008291.

[29] Benevenuta, S., Pancotti, C., Fariselli, P., Birolo, G. & Sanavia, T. An antisymmetric neural network to predict free energy changes in protein variants. Journal of Physics D: Applied Physics 54, 245403 (2021). URL 10.1088/1361-6463/abedfb.

[30] Dieckhaus, H., Brocidiacono, M., Randolph, N. Z. & Kuhlman, B. Transfer learning to leverage larger datasets for improved prediction of protein stability changes. Proceedings of the National Academy of Sciences 121, e2314853121 (2024). URL https://www.pnas.org/doi/10.1073/pnas.2314853121.

[31] Gordon, C. W., Lu, A. X. & Abbeel, P. Protein Language Model Fitness is a Matter of Preference(2024). URL https://openreview.net/forum?id=UvPdpa4LuV.

[32] Gong, C. et al. Evolution-Inspired Loss Functions for Protein Representation Learning (2024). URL https://openreview.net/forum?id=y5L8W0KRUX&referrer=%5Bthe%20profile%20of%20Chengyue%20Gong%5D(%2Fprofile%3Fid%3D~Chengyue Gong1).

[33] Luo, S. et al. Rotamer Density Estimator is an Unsupervised Learner of the Effect of Mutations on Protein-Protein Interaction (2022). URL https://openreview.net/forum?id=X9Yl1K2mD.

[34] Chaudhury, S., Lyskov, S. & Gray, J. J. PyRosetta: a script-based interface for implementing molecular modeling algorithms using Rosetta. Bioinformatics (Oxford, England) 26, 689–691 (2010).

[35] Dauparas, J. et al. Robust deep learning–based protein sequence design using ProteinMPNN. Science378, 49–56 (2022). URL https://www.science.org/doi/10.1126/science.add2187.

[36] Visani, G. M., Pun, M. N., Angaji, A. & Nourmohammad, A. Holographic-(V)AE: An end-to-end SO(3)-equivariant (variational) autoencoder in Fourier space. Physical Review Research 6, 023006 (2024). URL https://link.aps.org/doi/10.1103/PhysRevResearch.6.023006.

[37] Visani, G. M., Galvin, W., Pun, M. & Nourmohammad, A. H-Packer: Holographic Rotationally Equivariant Convolutional Neural Network for Protein Side-Chain Packing. Proceedings of the 18th Machine Learning in Computational Biology meeting, 230–249 (PMLR, 2024). URL https://proceedings.mlr.press/v240/visani24a.html.

[38] AlQuraishi, M. ProteinNet: a standardized data set for machine learning of protein structure. BMC Bioinformatics 20, 311 (2019). URL 10.1186/s12859-019-2932-0.

[39] Mei, H., Liao, Z. H., Zhou, Y. & Li, S. Z. A new set of amino acid descriptors and its application in peptide QSARs. Peptide Science 80, 775–786 (2005). URL https://onlinelibrary.wiley.com/doi/abs/10.1002/bip.20296. eprint: https://onlinelibrary.wiley.com/doi/pdf/10.1002/bip.20296.

[40] Lin, Z. et al. Evolutionary-scale prediction of atomic-level protein structure with a language model.Science 379, 1123–1130 (2023). URL https://www.science.org/doi/10.1126/science.ade2574.

[41] Pyo, A. G. T. et al. Data-Driven Discovery of Biophysical T Cell Receptor Cospecificity Rules. PRX Life 3, 033005 (2025). URL https://link.aps.org/doi/10.1103/14j1-wrh5.

[42] Broom, A., Jacobi, Z., Trainor, K. & Meiering, E. M. Computational tools help improve protein stability but with a solubility tradeoff. The Journal of Biological Chemistry 292, 14349–14361 (2017). URL https://pmc.ncbi.nlm.nih.gov/articles/PMC5582830/.

[43] Pancotti, C. et al. Predicting protein stability changes upon single-point mutation: a thorough comparison of the available tools on a new dataset. Briefings in Bioinformatics 23, bbab555 (2022). URL 10.1093/bib/bbab555.

[44] Che, Y. et al. Rational design of a highly immunogenic prefusion-stabilized F glycoprotein antigen for a respiratory syncytial virus vaccine. Science Translational Medicine 15, eade6422 (2023). URL https://www.science.org/doi/10.1126/scitranslmed.ade6422.

[45] Rahban, M. et al. Stabilization challenges and aggregation in protein-based therapeutics in the pharmaceutical industry. RSC Advances 13, 35947–35963 (2023). URL 10.1039/d3ra06476j.

[46] Tan, T. J. C. et al. High-throughput identification of prefusion-stabilizing mutations in SARS-CoV-2 spike. Nature Communications 14, 2003 (2023). URL https://www.nature.com/articles/s41467-023-37786-1.

[47] Dietterich, T. G. Ensemble Methods in Machine Learning. Multiple Classifier Systems, 1–15 (Springer, Berlin, Heidelberg, 2000).

[48] Visani, G. M. et al. T cell receptor specificity landscape revealed through de novo peptide design. Proceedings of the National Academy of Sciences 122, e2504783122 (2025). URL https://www.pnas.org/doi/10.1073/pnas.2504783122.

[49] Kwong, P. et al. Prefusion RSV F proteins and their use (2017). URL https://patents.google.com/patent/US9738689B2/en.

[50] Park, H. et al. Simultaneous Optimization of Biomolecular Energy Functions on Features from Small Molecules and Macromolecules. Journal of Chemical Theory and Computation 12, 6201–6212 (2016). URL 10.1021/acs.jctc.6b00819.

[51] Delgado, J., Radusky, L. G., Cianferoni, D. & Serrano, L. FoldX 5.0: working with RNA, small molecules and a new graphical interface. Bioinformatics 35, 4168–4169 (2019). URL 10.1093/bioinformatics/btz184.

[52] Shan, S. et al. Deep learning guided optimization of human antibody against SARS-CoV-2 variants with broad neutralization. Proceedings of the National Academy of Sciences 119, e2122954119 (2022). URL https://www.pnas.org/doi/full/10.1073/pnas.2122954119.

[53] Hsu, C. et al. Learning inverse folding from millions of predicted structures. Proceedings of the 39th International Conference on Machine Learning, 8946–8970 (PMLR, 2022). URL https://proceedings.mlr.press/v162/hsu22a.html.

[54] Tao, F., Sun, J., Gao, P., Gao, G. F. & Wu, B. Reliable prediction of protein–protein binding affinity changes upon mutations with Pythia-PPI. National Science Review 12, nwaf231 (2025). URL 10.1093/nsr/nwaf231.

[55] Jankauskaitė, J., Jiménez-Garcĺa, B., Dapkūnas, J., Fernández-Recio, J. & Moal, I. H. SKEMPI 2.0: an updated benchmark of changes in protein–protein binding energy, kinetics and thermodynamics upon mutation. Bioinformatics 35, 462–469 (2019). URL 10.1093/bioinformatics/bty635.

[56] Stourac, J. et al. FireProtDB: database of manually curated protein stability data. Nucleic Acids Research 49, D319–D324 (2021). URL 10.1093/nar/gkaa981.

[57] Levy, E. D. A Simple Definition of Structural Regions in Proteins and Its Use in Analyzing Interface Evolution. Journal of Molecular Biology 403, 660–670 (2010). URL https://www.sciencedirect.com/science/article/pii/S0022283610010168.

[58] Geiger, M. & Smidt, T. e3nn: Euclidean Neural Networks (2022). URL http://arxiv.org/abs/2207.09453. ArXiv:2207.09453 [cs].

[59] Cock, P. J. A. et al. Biopython: freely available Python tools for computational molecular biology and bioinformatics. Bioinformatics 25, 1422–1423 (2009). URL 10.1093/bioinformatics/btp163.

[60] Eastman, P. et al. OpenMM 4: A Reusable, Extensible, Hardware Independent Library for High Performance Molecular Simulation. Journal of Chemical Theory and Computation 9, 461–469 (2013). URL 10.1021/ct300857j.

[61] Word, J. M., Lovell, S. C., Richardson, J. S. & Richardson, D. C. Asparagine and glutamine: using hydrogen atom contacts in the choice of side-chain amide orientation1. Journal of Molecular Biology 285, 1735–1747 (1999). URL https://www.sciencedirect.com/science/article/pii/S0022283698924019.

[62] Ponder, J. W. & Case, D. A. in Force Fields for Protein Simulations Advances in Protein Chemistry, Vol. 66 of Protein Simulations 27–85 (Academic Press, 2003). URL https://www.sciencedirect.com/science/article/pii/S006532330366002X.

[63] Kingma, D. P. & Ba, J. Bengio, Y. & LeCun, Y. (eds) Adam: A Method for Stochastic Optimization. (eds Bengio, Y. & LeCun, Y.) 3rd International Conference on Learning Representations, ICLR 2015, San Diego, CA, USA, May 7–9, 2015, Conference Track Proceedings (2015). URL http://arxiv.org/abs/1412.6980.

[64] Pucci, F., Bernaerts, K. V., Kwasigroch, J. M. & Rooman, M. Quantification of biases in predictions of protein stability changes upon mutations. Bioinformatics 34, 3659–3665 (2018). URL 10.1093/bioinformatics/bty348.

[65] Caldararu, O., Blundell, T. L. & Kepp, K. P. A base measure of precision for protein stability predictors: structural sensitivity. BMC Bioinformatics 22, 88 (2021). URL 10.1186/s12859-021-04030-w.

[66] Camacho, C. et al. BLAST+: architecture and applications. BMC bioinformatics 10, 421 (2009).

[67] Lee, Y.-Z. et al. Rational design of uncleaved prefusion-closed trimer vaccines for human respiratory syncytial virus and metapneumovirus. Nature Communications 15, 9939 (2024). URL https://www.nature.com/articles/s41467-024-54287-x.

[68] Hsieh, C.-L. et al. Structure-based design of prefusion-stabilized SARS-CoV-2 spikes. Science 369, 1501–1505 (2020). URL https://www.science.org/doi/10.1126/science.abd0826.

