## Supplementary Information for "HERMES: Holographic Equivariant neuRal network model for Mutational Effect and Stability prediction"

### Extended analysis of antigen stabilization

Here, we present a more detailed analysis of our antigen stabilization predictions from Figures 6, 7 and Table S4. For each antigen, we contextualize the candidate mutations by their local structural environment and examine the characteristics and physico-chemical features of the HERMES-predicted top-ranked antigen-stabilizing substitutions (Fig. S21). This analysis is intended to help practitioners incorporate these models into design workflows and to motivate quantitative, context-dependent success criteria for real-world stabilization tasks. As demonstrated in the main text (Figs. 6, 7), we find that comparing the rank of the wild-type residue to that of putative stabilizing substitutions is an informative diagnostic of model behavior and performance.

**RSV-F.** RSV-F Cav1 stabilizing mutations S190F and V207L are well-predicted by different HERMES models (the stability-fine-tuned models trained on +cDNA117k and +Megascale, and HERMES-*amortized*), whereas Rosetta and ProteinMPNN struggle to identify these mutations. We therefore focus on these two residues to analyze, post hoc, the structural context underlying HERMES' predictions, with the goal of clarifying which classes of stabilizing mutations HERMES is best suited to characterize.

Both S190F and V207L enhance hydrophobic packing in an underpacked region near the trimer apex (Fig. S28). V207L is a significantly more anticipated mutation than S190F, indicated by a positive BLOSUM62 score; this is reflected in HERMES assigning a higher rank to the wild-type Val207 relative to the mutant Leu (Fig. S21 and Table S4). Notably, the highest-ranking residue predicted by HERMES models at position 207 is Ile (Fig. S21). Inspection of the native wild-type structure (PDB ID: 4JHW [1]) suggests that an Ile mutation would pack exceptionally well in the hydrophobic pocket surrounding residue 207, suggesting V207I might outperform the identified V207L mutation (Fig. S28A).

At position 190, the top-ranked substitutions from the wild-type Ser are Val or Ile. Structural examination suggests both residues can be accommodated within the existing 4JHW structure pocket without requiring the repacking of surrounding residues (Fig. S28B). Conversely, the known stabilizing mutation, Phe, appears to slightly overpack the region in the 4JHW crystal structure (Fig. S28B). While the HERMES-*fixed* model does not strongly prioritize Phe, both HERMES stability-fine-tuned models rank Phe third, behind the smaller Val and Ile. This indicates that fine-tuning on $\Delta\Delta G$ datasets enhances implicit reasoning regarding local repacking and structural relation possibilities, particularly for larger residues. The HERMES-*amortized* model also outperforms the HERMES-*fixed* model, likely due to the model's de-emphasis on strict steric constraints (Table S4).

The RSV-F TriC "acid-patch neutralization" substitutions D486H, D489H, and E487Q were stabilizing only in combination with F488W [1]— four tightly clustered residues acting synergistically (epistatically)—and the DENV-E pair A259W/T262R together introduce a favorable cation-$\pi$ interaction.

**Universal-HA.** We observe robust performance in recovering Universal-HA pH switch mutations across all tested models. All three mutations carry "unanticipated" BLOSUM62 scores.

The mutational-effect heatmaps (Fig. S21) show that, for HA, ProteinMPNN's preferences are narrowly concentrated, strongly favoring the reported stabilizing substitutions, whereas HERMES-*amortized*, HERMES-*fixed* (0.50 + Megascale), and Rosetta yield substantially broader preference profiles. For H355W, all HERMES models correctly indicate that the wild-type His is disfavored, but they rank other bulky aromatics (Tyr or Phe) above the stabilizing Trp. Despite being a known stabilizing mutation, Trp does not appear to fit in the 7VDF crystal structure without clashes (Fig. S29). This discrepancy may stem from the limitations of rigid-body mutagenesis (e.g., PyMOL's wizard), as subtle backbone movements are likely required to accommodate the mutation. This finding cautions against over-interpreting apparent clashes in a single experimentally determined static crystal structure, especially for bulky substitutions, and motivates follow-up evaluation with relaxation-aware structure prediction models or physics-based repacking/scoring tools to capture local flexibility and repacking that may render seemingly sterically forbidden mutations feasible.

**hMPV-F.** We first analyze stabilizing mutations in the M-104 hMPV variant [2]. The A159L substitution requires the nearby Ile-137 to adopt an alternative rotamer [2]. The two $\Delta\Delta G$-fine-tuned HERMES models prioritize Val (V) and Ile (I) over Leu (L) at position 159. We note that mutation to either Val or Ile at position 159 would require both Ile-137 and Leu-141 to adopt different rotamers to accommodate their beta-branched side chains (Fig. S30C). Nonetheless, A159V and A159I remain plausible stabilizing mutations and therefore, warrant experimental screening. ProteinMPNN identifies Ala as the most favorable residue at this position, indicating limited sensitivity to larger hydrophobic substitutions. By comparison, the $\Delta\Delta G$-fine-tuned HERMES models rank the wild-type Ala substantially lower (ordinal rank 5–7), while consistently favoring larger hydrophobic residues. Low rankings for buried hydrophobic positions may therefore indicate suboptimal core packing, particularly when alternative residues with greater side-chain volume are predicted. Although Leu is not the top-ranked substitution, the $\Delta\Delta G$-fine-tuned HERMES models correctly capture the preference for a residue larger than Ala to improve packing within this pocket.

The V203I substitution was recovered by nearly all models. A Val to Ile substitution is highly anticipated, with a BLOSUM62 score of 3. Notably, the $\Delta\Delta G$-fine-tuned HERMES models rank Ile as more favorable than the wild-type Val, whereas ProteinMPNN, ThermoMPNN, HERMES-*fixed*, and HERMES-*amortized* prefer the wild-type Val. In accordance with this mutation being more highly anticipated, it is likely that the pocket occupied by V203 is less underpacked relative to A159L in the M-104 variant, with the wild-type residue still being quite highly-favored. The subtle nature of Val to Ile mutations may result in less signal overall when comparing amino acid ranks as we do in this work.

The V449D substitution replaces a surface-exposed hydrophobic residue with a polar side chain and introduces hydrogen-bonding contacts to Asn298 (Fig. S30A). ProteinMPNN, ThermoMPNN, and all HERMES variants instead rank Glu as the top substitution; structural inspection suggests that Glu could form similar hydrogen bonds, and may therefore also be stabilizing (Fig. S30A). Overall, the models performed well in identifying mutations that enable additional hydrogen bonding.

For V430Q, the predicted $\Delta\Delta G$ reported in the original study is small in magnitude (albeit negative), suggesting that it should not be classified as strongly stabilizing, as suggested in the original study [2]. Consistent with this, neither of the $\Delta\Delta G$–fine-tuned HERMES models prioritize V430Q. Interestingly, the only model that ranks V430Q as the top substitution is HERMES-*fixed*.

In the MPV-2c variant [3], recovery of V112R depends on providing the model with the trimeric PDB. The introduced Arg forms interprotomer van der Waals contacts and intraprotomer hydrogen bonds that stabilize the prefusion trimer (Fig. S30B). Because this mechanism is inherently multimeric and not captured by a strictly local energetic proxy, it may explain why the $\Delta\Delta G$-fine-tuned HERMES models do not prioritize V112R. In contrast, HERMES-*amortized* ranks Arg second, slightly favoring the more conservative Leu, which could plausibly adjust hydrophobic interprotomer contacts.

D209E is correctly predicted by ProteinMPNN, HERMES-*fixed*, and HERMES-*amortized*, whereas the $\Delta\Delta G$-fine-tuned HERMES models perform poorly in proposing this mutation. In particular, the $\Delta\Delta G$-fine-tuned models rank bulky hydrophobics (Leu, Met, Ile, Val) ahead of the charge-conserving Glu. Although the charged substitution D209E plausibly stabilizes the prefusion state via favorable polar interactions, the stabilizing potential of these alternative hydrophobic substitutions remains untested.

E453P replaces a glutamate with proline, a substitution that can be strongly stabilizing by rigidifying locally flexible regions. Among the evaluated methods, only HERMES-*amortized* predicts this mutation. ProteinMPNN ranks the wild-type Glu as most favored and Pro as most disfavored (rank 1 vs. 20), while ThermoMPNN partially recovers the substitution but still favors the native Glu. Consistent with these mixed signals, the original study [3] reports that E453P improves stability but reduces expression, and that E453Q yields weaker stabilization with improved expression. This highlights the poorly understood coupling between stability and expression and suggests that some models may systematically downweigh proline substitutions, potentially because prolines are underrepresented in natural sequences, and they appropriately learned to rarely recommend them. The fact that HERMES-*amortized*—but not HERMES-*fixed*—recovers E453P suggests that modeling relaxed neighborhoods helps identify sites that can accommodate Pro's constrained Ramachandran geometry. Finally, we suspect that the utility and stabilizing ability of proline substitutions in antigen design is not fully captured by HERMES when fine-tuned with the cDNA117k or Megascale datasets, as the proteins represented in those datasets are much smaller, and lack the interplay of conformational switching between prefusion and post-fusion.

**DENV-E.** We observe mixed performance on DENV-E stabilizing mutations. S29K is recovered only by ProteinMPNN and ThermoMPNN. Although S29K, T33V, and A35M ("PM4" mutations as a group) are spatially proximal and may act synergistically, they were originally identified by Rosetta site-saturation mutagenesis, suggesting they should be accessible to single point-mutation scoring schemes [4]. In the 1OAN structure [5], introducing Lys at position 29 appears to create substantial steric clashes across rotamers (Fig. S31A). S29K and A35M are not anticipated by BLOSUM62, whereas T33V is neutral according to BLOSUM62 (score of 0, Table S4) but is strongly preferred by all models tested (Fig. 6and Table S4). Structurally, T33 lines a hydrophobic pocket, making the isosteric Val substitution easier to predict. Notably, the PM4 mutations are not present in the best-available SC12 structure (PDB: 6WY1) [6].

A major challenge in stabilizing DENV-E is strengthening homodimer interactions. A259W and T262R were mutations made at the dimer interface with the intention of enhancing the strength of the homodimer. A259W is highly unanticipated (BLOSUM62 = $-3$) but, together with T262R, can form a favorable cation–$\pi$ interaction while improving packing against the neighboring protomer (Fig. 7A.4). Notably, the HERMES model fine-tuned on Megascale favors A259W despite its apparent synergy with T262R; this could reflect sensitivity to the interprotomer packing geometry, although a chance effect cannot be excluded. We also expect the generally weak recovery of T262R across models to improve when Trp is present at position 259, consistent with the coupled nature of this interface motif.

Like E453P in hMPV-F, the proline substitution T280P is recovered only by HERMES-*amortized*, further highlighting the model's ability to reason about proline accommodation. F279W fills an underpacked region. However, comparison of the stabilized 6WY1 structure with the native 1OAN crystal structure suggests that backbone movement (residues 269–281) is required to optimally fit Trp, potentially aided by T280P (Fig. S31B). ThermoMPNN correctly reasons regarding the Trp introduction, and HERMES-*fixed* + Megascale ranks Trp second. Other stability-fine-tuned HERMES models prefer Phe, Ile, or Leu, likely adhering more strictly to the steric limitations of the 1OAN backbone.

We observe strong reasoning across almost all HERMES models, ProteinMPNN, and Rosetta in identifying G106D, which introduces stabilizing polar and electrostatic contacts (Fig. S31C). Curiously, ThermoMPNN struggles with this specific mutation.

**SARS-CoV-2 Spike.** The spike protein of SARS-CoV-2 was initially stabilized through the introduction of two proline mutations ("S-2P"), and subsequent work identified four additional prolines that further improve stability and expression of the full-length spike [7]. Here we evaluate recovery of these four additional proline sites (excluding the original S-2P mutations). Consistent with its strong performance on proline substitutions, HERMES-*amortized* ranks Pro as the top choice at all four positions. ProteinMPNN also performs well, recovering Pro at 3/4 sites.

**Summary.** Our analysis across multiple antigens reveals several consistent themes regarding model performance and decision-making logic.

First, we observe distinct behaviors regarding hydrophobic packing. The HERMES models fine-tuned on stability datasets consistently demonstrate enhanced reasoning for hydrophobic core packing. These models frequently suggest larger hydrophobic residues (e.g., Ile or Phe) to fill underpacked cavities where models without explicit stability training might prefer the native residue or smaller conservative substitutions. However, this sensitivity can sometimes lead to "confusion" among similar hydrophobic amino acids (e.g., Val vs. Ile vs. Leu), where the models correctly identify the chemical property needed but may rank several hydrophobic options similarly.

Second, the interplay between structural rigidity and model flexibility is evident in the prediction of proline mutations. HERMES-*amortized* consistently outperforms other models in identifying stabilizing proline substitutions. This suggests that the model's training, which incorporates relaxed neighborhoods, allows it to better identify backbone locations capable of accommodating the steric constraints of proline, whereas other methods more often penalize these stabilizing mutations due to perceived steric incompatibilities of the provided protein structure.

Third, we note differences in the mutational landscape profiles predicted by different architectures (Fig. S21. ProteinMPNN tends to produce narrow substitution profiles, often assigning very low probabilities to non-native residues unless the signal is overwhelmingly strong. In contrast, HERMES models–particularly HERMES-*amortized* and those fine-tuned on Megascale stability data–exhibit broader mutational profiles. This broader landscape may be more advantageous for design applications, as it provides a richer set of plausible candidate hypotheses for experimental validation and may guide a practitioner's intuition regarding the nature or predicted effects of various substitutions in a more granular manner.

Finally, while ProteinMPNN and Rosetta remain powerful tools for sequence recovery, the HERMES-*amortized* and stability-fine-tuned models demonstrate a unique capacity to prioritize mutations that improve local packing density even when such mutations appear sterically challenging in the provided structure. This highlights the utility of using an ensemble of models to capture different modes of stabilization, from electrostatic optimization to hydrophobic core repacking and backbone rigidification.

### References


[1] McLellan, J. S. *et al.* Structure-Based Design of a Fusion Glycoprotein Vaccine for Respiratory Syncytial Virus. *Science* **342**, 592–598 (2013). URL https://www.science.org/doi/10.1126/science.1243283.

[2] Gonzalez, K. J. *et al.* A general computational design strategy for stabilizing viral class I fusion proteins. *Nature Communications* **15**, 1335 (2024). URL https://www.nature.com/articles/s41467-024-45480-z.

[3] Bakkers, M. J. G. *et al.* Efficacious human metapneumovirus vaccine based on AI-guided engineering of a closed prefusion trimer. *Nature Communications* **15**, 6270 (2024). URL https://www.nature.com/articles/s41467-024-50659-5.

[4] Phan, T. T. N. *et al.* A conserved set of mutations for stabilizing soluble envelope protein dimers from dengue and Zika viruses to advance the development of subunit vaccines. *Journal of Biological Chemistry* **298**, 102079 (2022). URL https://www.sciencedirect.com/science/article/pii/S0021925822005191.

[5] Modis, Y., Ogata, S., Clements, D. & Harrison, S. C. A ligand-binding pocket in the dengue virus envelope glycoprotein. *Proceedings of the National Academy of Sciences* **100**, 6986–6991 (2003). URL https://www.pnas.org/doi/full/10.1073/pnas.0832193100.

[6] Kudlacek, S. T. *et al.* Designed, highly expressing, thermostable dengue virus 2 envelope protein dimers elicit quaternary epitope antibodies. *Science Advances* **7**, eabg4084 (2021). URL https://www.science.org/doi/10.1126/sciadv.abg4084.

[7] Hsieh, C.-L. *et al.* Structure-based design of prefusion-stabilized SARS-CoV-2 spikes. *Science* **369**, 1501–1505 (2020). URL https://www.science.org/doi/10.1126/science.abd0826.

[8] Mei, H., Liao, Z. H., Zhou, Y. & Li, S. Z. A new set of amino acid descriptors and its application in peptide QSARs. *Peptide Science* **80**, 775–786 (2005). URL https://onlinelibrary.wiley.com/doi/abs/10.1002/bip.20296. _eprint: https://onlinelibrary.wiley.com/doi/pdf/10.1002/bip.20296.

[9] Lee, Y.-Z. *et al.* Rational design of uncleaved prefusion-closed trimer vaccines for human respiratory syncytial virus and metapneumovirus. *Nature Communications* **15**, 9939 (2024). URL https://www.nature.com/articles/s41467-024-54287-x.

[10] Milder, F. J. *et al.* Universal stabilization of the influenza hemagglutinin by structure-based redesign of the pH switch regions. *Proceedings of the National Academy of Sciences* **119**, e2115379119 (2022). URL https://www.pnas.org/doi/full/10.1073/pnas.2115379119.

[11] Byrne, P. O. & McLellan, J. S. Principles and practical applications of structure-based vaccine design. *Current Opinion in Immunology* **77**, 102209 (2022). URL https://www.sciencedirect.com/science/article/pii/S0952791522000565.

[12] Blaabjerg, L. M. *et al.* Rapid protein stability prediction using deep learning representations. *eLife* **12**, e82593 (2023). URL https://doi.org/10.7554/eLife.82593.

[13] Diaz, D. J. *et al.* Stability Oracle: a structure-based graph-transformer framework for identifying stabilizing mutations. *Nature Communications* **15**, 6170 (2024). URL https://www.nature.com/articles/s41467-024-49780-2.

[14] Tsuboyama, K. *et al.* Mega-scale experimental analysis of protein folding stability in biology and design. *Nature* **620**, 434–444 (2023). URL https://www.nature.com/articles/s41586-023-06328-6.

[15] Levy, E. D. A Simple Definition of Structural Regions in Proteins and Its Use in Analyzing Interface Evolution. *Journal of Molecular Biology* **403**, 660–670 (2010). URL https://www.sciencedirect.com/science/article/pii/S0022283610010168.

| **Model** | **Accuracy** Pyrosetta pre-processing | **Accuracy** Biopython pre-processing |
|---|---|---|
| HERMES-*fixed* 0.00 | 0.73 | 0.75 |
| HERMES-*fixed* 0.50 | 0.64 | 0.65 |
| HERMES-*amortized* 0.00 | 0.55 | 0.47 |
| HERMES-*amortized* 0.50 | 0.50 | 0.44 |
| HERMES-*fixed* 0.00 + Ros | 0.41 | 0.40 |
| HERMES-*fixed* 0.50 + Ros | 0.38 | 0.37 |
| HERMES-*fixed* 0.00 + cDNA117k | 0.47 | 0.45 |
| HERMES-*fixed* 0.50 + cDNA117k | 0.39 | 0.38 |
| HERMES-*amortized* 0.00 + cDNA117k | 0.37 | - |
| HERMES-*amortized* 0.50 + cDNA117k | 0.34 | - |
| HERMES-*fixed* 0.00 + cDNA117k train ESMFold | 0.46 | 0.49 |
| HERMES-*fixed* 0.50 + cDNA117k train ESMFold | 0.40 | 0.40 |
| HERMES-*fixed* Untr. 0.00 + cDNA117k | 0.09 | - |
| HERMES-*fixed* Untr. 0.50 + cDNA117k | 0.08 | - |

**Table S1 Accuracy of HERMES models on wildtype amino acid classification on all sites across 40 CASP12 test proteins.** Accuracy is defined as proportion of sites for which the wild-type amino acid is predicted with the highest probability among all 20 canonical amino acids. Model names indicate the architecture, the coordinate-noise amplitude used, and when applicable, the fine-tuning dataset (listed after "+"); *Untr.* is short for *Untrained*, indicating models that had no pre-training and were instead only trained on stability effects. Accuracy is reported for the two pre-processing schemes (with PyRosetta and Biopython) used in HERMES.

| **Model** | **hh:mm:ss** |
|---|---|
| HERMES-*fixed* 0.50 | 00:11:23 |
| HERMES-*amortized* 0.50 | 00:11:23 |
| HERMES-*relaxed* 0.50 | 12:13:20 |

**Table S2 Inference speed of HERMES models on the T2837 dataset.** Runtimes are reported in hours (hh), minutes (mm), and seconds (ss) for inference on the T2837 dataset, which comprises 2837 mutation effects across 129 proteins. For HERMES-*fixed* and HERMES-*amortized*, the script 'mutation_effect_prediction_with_hermes.py' was used; for HERMES-*relaxed*, the script 'mutation_effect_prediction_with_hermes_with_relaxation.py' was used. Both scripts, along with the dataset 'csv' file, are available in our GitHub repository. All models were executed using a single CPU and a single A40 GPU.

| Category | Description |
|---|---|
| Hydrophobic Property | 1. Retention coefficient in TFA |
| | 2. Free energy of solution in water |
| | 3. Solvation free energy |
| | 4. Melting point |
| | 5. Number of hydrogen-bond donors |
| | 6. Number of full nonbonding orbitals |
| | 7. Partition energy |
| | 8. Hydration number |
| | 9. Retention coefficient in high performance liquid chromatography (HPLC), pH 7.4 |
| | 10. Retention coefficient in HPLC, pH 2.1 |
| | 11. Partition coefficient in thin-layer chromatography |
| | 12. Retention coefficient at pH 2 |
| | 13. $R_f$ for 1-N-(4-nitrobenzofurazono)-amino acids in ethyl acetate/pyridine/water |
| | 14. $\Delta G$ of transfer from organic solvent to water |
| | 15. Hydration potential or free energy of transfer from vapor phase to water |
| | 16. $R_f$, salt chromatography |
| | 17. $\log D$, partition coefficient at pH 7.1 for acetamide derivatives of amino acids in octanol/water |
| | 18. $\Delta G = RT \log f$, $f$ = fraction buried/accessible amino acids in 22 proteins |
| Steric Property | 19. Average volume of buried residue |
| | 20. Residue accessible surface area in tripeptide |
| | 21. Graph shape index |
| | 22. Normalized van der Waals volume |
| | 23. STERMIMOL length of the side chain |
| | 24. STERMIMOL minimum width of the side chain |
| | 25. STERMIMOL maximum width of the side chain |
| | 26. Average accessible surface area |
| | 27. Distance between $C_\alpha$ and centroid of side chain |
| | 28. Side-chain angle $\theta$ |
| | 29. Side-chain torsion angle $\phi$ |
| | 30. Radius of gyration of side chain |
| | 31. Van der Waals parameter $R_0$ |
| | 32. Van der Waals parameter $\varepsilon$ |
| | 33. Refractivity |
| | 34. Value of $\theta$ (i) |
| | 35. Substituent van der Waals volume |
| Electronic Property | 36. $\alpha$CH chemical shifts |
| | 37. $\alpha$NH chemical shifts |
| | 38. A parameter of charge transfer capability |
| | 39. A parameter of charge transfer donor capability |
| | 40. Nuclear magnetic resonance (NMR) chemical shift of $\alpha$ carbon |
| | 41. Localized electrical effect |
| | 42. Positive charge |
| | 43. Negative charge |
| | 44. Polarity |
| | 45. Net charge |
| | 46. Amphipathicity index |
| | 47. Isoelectric point |
| | 48. Electron-ion interaction potential values |
| | 49. $pK_{NH_2}$ ($NH_2$ on $C_\alpha$) |
| | 50. $pK_{COOH}$ (COOH on $C_\alpha$) |

**Table S3 Table of amino acid properties used for comparison with substitution matrices.** Values associated with each amino acid are listed in ref. [8].

| antigen PDB id | stabilized name | mutation | BLOSUM62 score ; rank | Rosetta | Protein MPNN 0.30 | Thermo MPNN + Megascale | HERMES *-fixed* 0.50 | HERMES *-FT_relaxed* 0.50 | HERMES *-fixed* 0.50 + cDNA117k | HERMES *-fixed* 0.50 + Megascale |
|---|---|---|---|---|---|---|---|---|---|---|
| | | | | $r_{wt} \to r_{mt}$ | $r_{wt} \to r_{mt}$ | $r_{wt} \to r_{mt}$ | $r_{wt} \to r_{mt}$ | $r_{wt} \to r_{mt}$ | $r_{wt} \to r_{mt}$ | $r_{wt} \to r_{mt}$ |
| RSV-F 4JHW | Cav1 [1] | S190F | -2 ; 16 | 19 → 11 | 4 → 6 | 14 → 3 | 4 → 8 | 11 → 4 | 12 → 3 | 13 → 3 |
| | | V207L | 1 ; 3 | 9 → 4 | 2 → 7 | 6 → 5 | 2 → 3 | 3 → 2 | 3 → 2 | 3 → 2 |
| | Uncl. [9] | S215P | -1 ; 13 | 13 → 1 | 1 → 16 | 10 → 20 | 10 → 15 | 6 → 3 | 16 → 19 | 4 → 20 |
| | TriC [1] | D486H | -1 ; 7 | 9 → 2 | 1 → 9 | 1 → 8 | 5 → 4 | 10 → 4 | 20 → 9 | 18 → 9 |
| | | E487Q | 2 ; 3 | 2 → 9 | 1 → 13 | 1 → 12 | 2 → 3 | 4 → 5 | 11 → 12 | 14 → 12 |
| | | F488W | 1 ; 3 | 15 → 17 | 1 → 14 | 3 → 1 | 2 → 6 | 1 → 13 | 1 → 3 | 1 → 2 |
| | | D489H | -1 ; 7 | 5 → 2 | 3 → 14 | 5 → 12 | 11 → 8 | 18 → 15 | 20 → 11 | 17 → 12 |
| HA 7VDF | Universal-HA [10] | H355W | -2 ; 12 | 5 → 1 | 5 → 1 | 4 → 2 | 4 → 3 | 7 → 2 | 5 → 3 | 8 → 3 |
| | | K380I | -3 ; 19 | 17 → 1 | 9 → 1 | 11 → 1 | 8 → 4 | 14 → 3 | 13 → 2 | 17 → 2 |
| | | E432I | -3 ; 19 | 14 → 9 | 12 → 2 | 13 → 2 | 9 → 4 | 15 → 5 | 12 → 3 | 14 → 3 |
| hMPV-F 5WB0 | M104 [2] | L130D | -4 ; 19 | 15 → 3 | 8 → 9 | 4 → 10 | 5 → 9 | 12 → 2 | 8 → 2 | 10 → 4 |
| | | A159L | -1 ; 10 | 6 → 1 | 1 → 12 | 3 → 5 | 1 → 7 | 2 → 4 | 7 → 3 | 5 → 3 |
| | | V203I | 3 ; 2 | 4 → 3 | 1 → 2 | 1 → 2 | 1 → 2 | 2 → 1 | 3 → 1 | 2 → 1 |
| | | V430Q | -2 ; 10 | 12 → 3 | 7 → 4 | 12 → 6 | 8 → 1 | 10 → 5 | 5 → 12 | 10 → 8 |
| | | V449D | -3 ; 16 | 13 → 8 | 13 → 3 | 16 → 2 | 11 → 2 | 15 → 2 | 18 → 2 | 19 → 6 |
| | MPV-2cREKR [3] | V112R | -3 ; 19 | 6 → 13 | 8 → 1 | 2 → 1 | 2 → 9 | 8 → 3 | 4 → 11 | 5 → 10 |
| | | D209E | 2 ; 2 | 8 → 1 | 11 → 1 | 12 → 4 | 3 → 1 | 14 → 3 | 15 → 6 | 16 → 7 |
| | | V231I | 3 ; 2 | 3 → 1 | 3 → 1 | 3 → 1 | 2 → 1 | 2 → 1 | 3 → 1 | 3 → 1 |
| | | E453P | -1 ; 10 | 10 → 2 | 1 → 20 | 1 → 2 | 8 → 19 | 13 → 1 | 17 → 16 | 19 → 6 |
| | Uncl. [9] | E80D | 2 ; 2 | 18 → 13 | 3 → 1 | 13 → 18 | 1 → 12 | 7 → 17 | 15 → 19 | 14 → 18 |
| | | V155P | -2 ; 13 | 10 → 20 | 4 → 20 | 1 → 20 | 1 → 19 | 1 → 20 | 2 → 20 | 4 → 20 |
| DENV-E 1OAN | SC12 [4] | S29K | 0 ; 6 | 4 → 3 | 3 → 1 | 7 → 2 | 1 → 12 | 2 → 5 | 6 → 12 | 9 → 14 |
| | | T33V | 0 ; 3 | 9 → 4 | 3 → 1 | 6 → 1 | 3 → 1 | 5 → 1 | 10 → 1 | 10 → 1 |
| | | A35M | -1 ; 13 | 11 → 4 | 6 → 1 | 10 → 8 | 1 → 12 | 1 → 11 | 1 → 7 | 6 → 7 |
| | | G106D | -1 ; 5 | 18 → 1 | 11 → 1 | 6 → 11 | 8 → 1 | 6 → 2 | 20 → 1 | 18 → 1 |
| | | A259W | -3 ; 20 | 7 → 1 | 1 → 3 | 4 → 11 | 1 → 20 | 1 → 13 | 1 → 2 | 2 → 1 |
| | | T262R | -1 ; 11 | 15 → 3 | 1 → 15 | 18 → 7 | 3 → 14 | 10 → 3 | 16 → 3 | 17 → 4 |
| | | F279W | 1 ; 3 | 3 → 1 | 4 → 11 | 4 → 1 | 1 → 6 | 3 → 7 | 1 → 4 | 1 → 2 |
| | | T280P | -1 ; 13 | 12 → 2 | 7 → 10 | 9 → 14 | 4 → 19 | 6 → 2 | 2 → 20 | 8 → 19 |
| SARS-Cov-2 6VSB | hexapro [7] | F817P | -4 ; 20 | 3 → 1 | 17 → 1 | 1 → 12 | 2 → 17 | 5 → 1 | 3 → 19 | 2 → 19 |
| | | A892P | -1 ; 14 | 16 → 3 | 5 → 1 | 2 → 4 | 2 → 1 | 4 → 1 | 3 → 1 | 6 → 1 |
| | | A899P | -1 ; 14 | 8 → 19 | 7 → 9 | 7 → 18 | 2 → 6 | 4 → 1 | 8 → 15 | 5 → 16 |
| | | A942P | -1 ; 14 | 5 → 12 | 4 → 1 | 2 → 8 | 1 → 5 | 2 → 1 | 2 → 1 | 2 → 1 |
| | proportion of correctly and strongly suggested mutations | | | **20/33** | **15/33** | **11/33** | 8/33 | **19/33** | **15/33** | **13/33** |
| | proportion of correctly and at least moderately suggested mutations | | | **23/33** | **16/33** | 14/33 | 11/33 | **23/33** | **16/33** | **17/33** |
| | proportion of correctly and at least weakly suggested mutations | | | **27/33** | 16/33 | 16/33 | 12/33 | **24/33** | 19/33 | 22/33 |

**Table S4 Predicting antigen-stabilizing mutations with HERMES: extended results.** Recall for different models (columns) is evaluated on 33 previously reported antigen-stabilizing mutations (rows) spanning five viral antigens. For each antigen, we list the PDB structure used for scoring and the publication(s) that originally reported the mutation. Mutations are specified as wild-type→mutant substitutions at the annotated site. Seven models are compared (columns). We additionally report the BLOSUM62 substitution score for each mutation and the mutant's rank among the 20 possible amino acid substitutions for the wild-type residue (per BLOSUM62). For each model and mutation, predicted ranks of the wild-type and mutant amino acids are shown as $r_{wt} \to r_{mt}$. Dagger symbols (†) indicate mutations originally proposed as stabilizing by Rosetta-based pipelines in the source reference. When a model ranks the mutant better than the wild type ($r_{mt} < r_{wt}$), the cell is shaded by the prediction strength based on the value of $r_{mt}$: dark green, strongly suggested ($r_{mt} \leq 3$); light green, moderately suggested ($4 \leq r_{mt} \leq 6$); light yellow, weakly suggested ($r_{mt} \geq 6$). Column summaries report counts of strongly, at least moderately, and at least weakly suggested mutations (out of 33); **bold** indicates significance (p-value $< 0.05$) for the number of recalled mutations relative to a random null model (see Fig. S24 for p-values and Methods for details.) All structures were scored in their native multimeric states, generating symmetric partners when needed. ThermoMPNN's native mode predicts mutation effects only for monomers, ignoring multimeric assemblies even when present in the input structure. "Uncl." stands for uncleaved prefusion-closed state.

| PDB id | # of monomer sites | HERMES GPU [seconds] all sites | HERMES CPU [seconds] all sites | Rosetta [CPU-hours] one site | Rosetta [CPU-hours] all sites |
|---|---|---|---|---|---|
| pre-f. 4JHW | 449 | 57 | 112 | 150 | 67,350 |
| pre-f. 7VDF | 485 | 64 | 118 | 164 | 79,540 |
| pre-f. 5WB0 | 442 | 43 | 154 | 151 | 66,742 |
| pre-f. 1OAN | 394 | 32 | 151 | 59 | 23,246 |
| pre-f. 6VSB | 968 | 69 | 265 | 156 | 151,008 |

**Table S5 Execution times for saturation mutagenesis predictions on the viral antigens considered in this study.** Execution times (in seconds) of HERMES apply to HERMES-*fixed* and HERMES-*amortized* models, regardless of whether zero-shot or fine-tuned. Times were computed when running the script `run_hermes_on_pdbfiles.py` providing as input the pdbfile as well as a single monomeric chain. A single CPU core with 64GB of memory was used, and a NVIDIA A40 GPU when applicable. For Rosetta, we computed times (in CPU-hours) for a single CPU core with 4 GBs of memory, and averaging 10 relaxation instances, which we consider the minimum number of instances for robust results. Times for all sites in the structure were extrapolated by multiplying the calculated average time for a single mutation by the number of monomeric sites.

| antigen PDB id | stabilized name | mutation | mutation type | is synergistic | notes |
|---|---|---|---|---|---|
| RSV-F pre-f. 4JHW | Cav1 [1] | S190F | cavity-filling | False | |
| | | V207L | cavity-filling | False | |
| | Uncl. [9] | S215P | proline | False | |
| | TriC [1] | D486H | electrostatic | True | |
| | | E487Q | electrostatic | True | |
| | | F488W | cavity-filling | True | |
| | | D489H | electrostatic | True | |
| HA pre-f. 7VDF | Universal-HA [10] | H355W | cavity-filling | False | |
| | | K380I | cavity-filling | False | |
| | | E432I | cavity-filling | False | |
| hMPV-F pre-f. 5WB0 | M104 [2] | L130D | electrostatic | False | |
| | | A159L | cavity-filling | False | |
| | | V203I | cavity-filling | False | |
| | | V430Q | electrostatic | False | |
| | | V449D | electrostatic | False | |
| | MPV-2cREKR [3] | V112R | electrostatic | False | |
| | | D209E | | False | same charge, slightly different size: unclear |
| | | V231I | cavity-filling | False | |
| | | E453P | proline | False | |
| | Uncl. [9] | E80D | | False | same charge, slightly different size: unclear |
| | | V155P | proline | False | |
| DENV-E pre-f. 1OAN | SC12 [4] | S29K | electrostatic | False | |
| | | T33V | cavity-filling | False | |
| | | A35M | cavity-filling | False | |
| | | G106D | electrostatic | False | |
| | | A259W | cavity-filling | True | |
| | | T262R | electrostatic | True | |
| | | F279W | cavity-filling | False | |
| | | T280P | proline | False | |
| SARS-Cov-2 pre-f. 6VSB | hexapro [7] | F817P | proline | False | |
| | | A892P | proline | False | |
| | | A899P | proline | False | |
| | | A942P | proline | False | |

**Table S6 Characteristics of antigen-stabilizing mutations.** Hand-curated mutation types are listed for antigen-stabilizing mutations reported in Fig. 6and Table S4. Cavity-filling mutations are defined as substitutions to hydrophobic residues that are larger than the wild-type when the wild-type is also hydrophobic. Electrostatic mutations are substitutions that change the residue's net charge. Proline mutations correspond to substitutions to proline. Synergistic mutations were identified through structural reasoning based on the spatial arrangement of mutations within the corresponding structure; see ref. [11] for a breakdown of mutation types considered in the structure-based vaccine design literature. "Uncl." stands for "Uncleaved Prefusion-Closed".

| | | CASP12 | | ProteinMPNN | |
|---|---|---|---|---|---|
| antigen | query length | hit length | % similarity | hit length | % similarity |
| RSV-F | 574 | 481 | 76.0 | 414 | 65.0 |
| HA | 550 | 317 | 45.8 | 515 | 78.5 |
| hMPV-F | 530 | 481 | 33.0 | 414 | 27.9 |
| DENV-E | 394 | 394 | 96.4 | 402 | 98.0 |
| SARS-Cov-2 | 1288 | 46 | 3.3 | 1274 | 98.9 |

**Table S7 Closest sequence match for each antigen in the CASP12 and ProteinMPNN pre-training sets.**

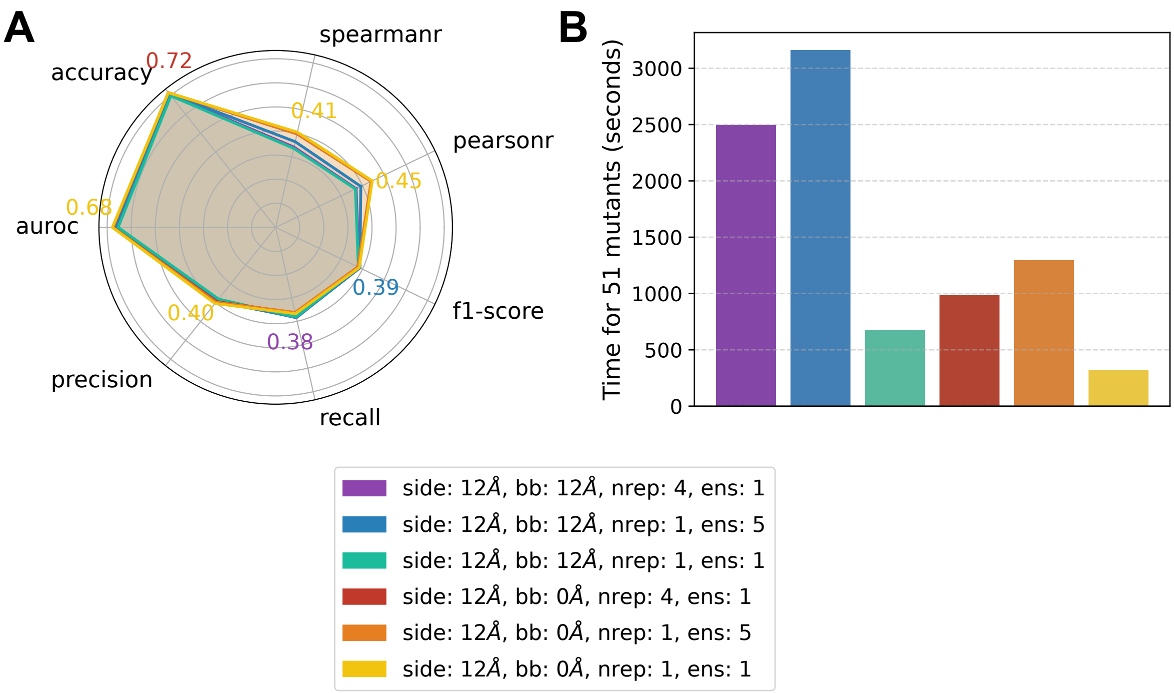


**Figure S1 Ablation of PyRosetta fastrelax parameters for HERMES-*relaxed* 0.50 on the cDNA117k dataset.** **(A)** Classification accuracy metrics, analogous to those shown in Fig. 2, are reported for HERMES-*relaxed* models evaluated on relaxed mutant structures using different PyRosetta fastrelax parameters (indicated by color). HERMES-*relaxed* scores a mutation as the log-probability difference between the mutant and wild-type amino acids. The wild-type log-probability is evaluated on the wild-type structure, while the mutant log-probability is evaluated on the wild-type structure after introducing the mutation and performing local relaxation. We use the PyRosetta fastrelax protocol and vary the following parameters, noting that the procedure is stochastic: (1) **side**, the distance cutoff for side-chain relaxation; (2) **bb**, the distance cutoff for backbone relaxation; (3) **nreps**, the number of protocol repetitions, with the lowest-energy conformation retained; (4) **ens**, the ensemble size, where predictions are averaged over relaxations obtained with different random seeds. **(B)** Inference speed for predicting mutational effects on 51 mutants across the same PyRosetta fastrelax parameters as in (A) (colors). A single NVIDIA A40 GPU and a single CPU with 64G of memory were used for all parameters.

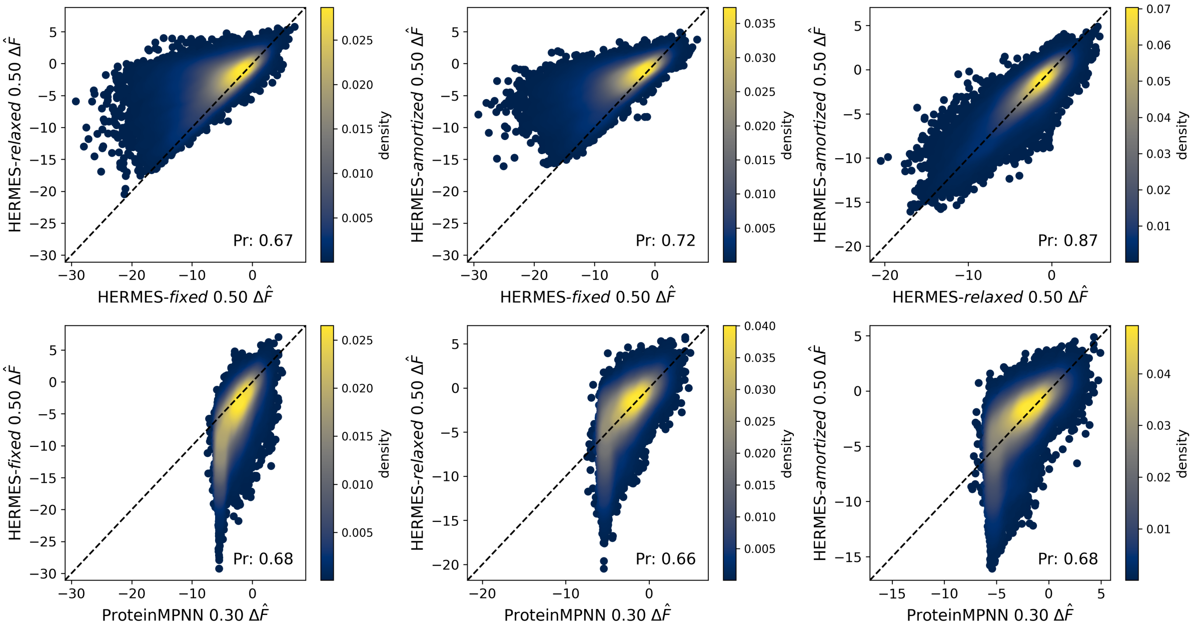


**Figure S2 Comparison of zero-shot model predictions on the Megascale test set.** For each substitution in the Megascale test set, the predicted change in amino acid propensity upon mutation ($\delta \log p$) is compared between two models in each panel. Color indicates local point density (blue denotes low density and yellow denotes high density). The reported "Pr" in each panel corresponds to the Pearson correlation coefficient between the predictions of the model pair. Model names indicate the architecture and the coordinate-noise amplitude used. $\Delta\hat{F}$ indicates the model's prediction, following Equations 1 and 2.

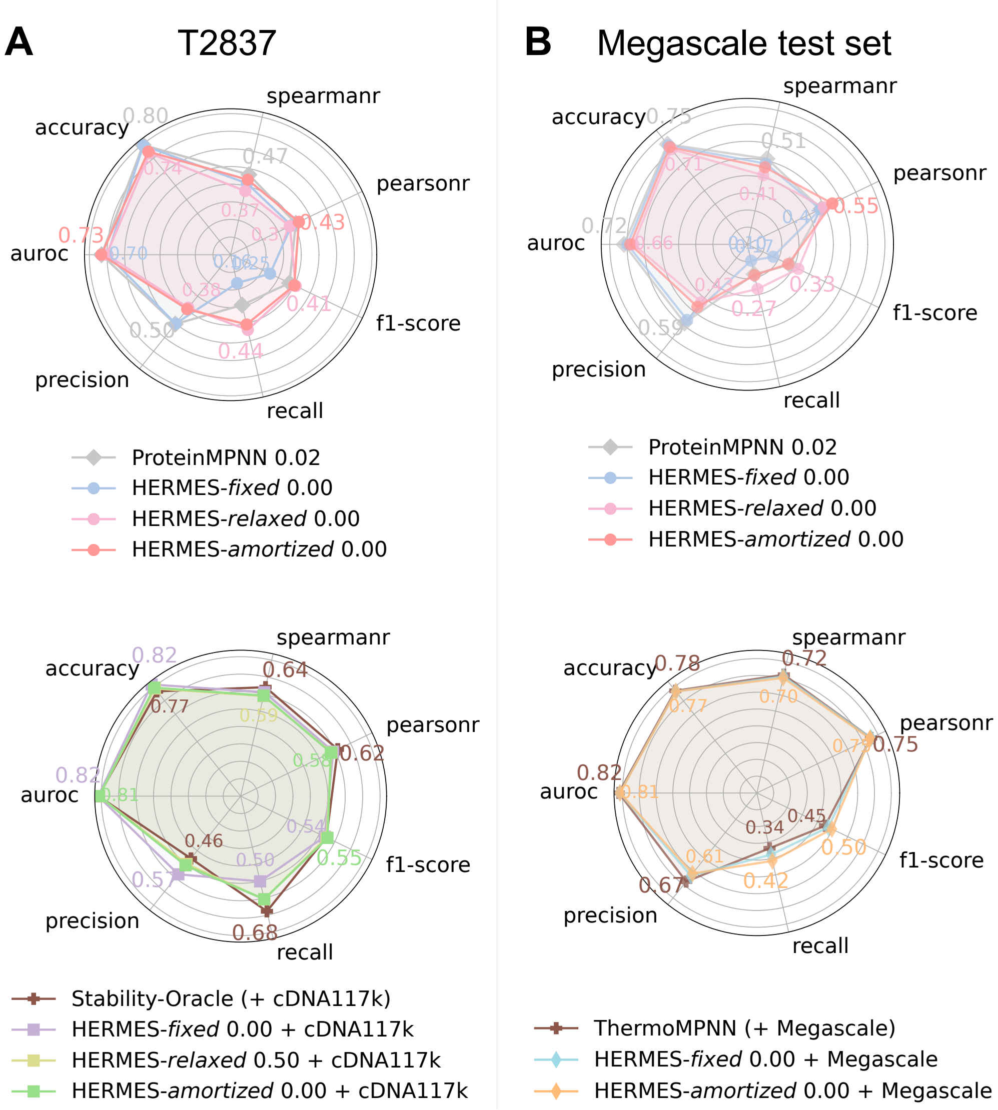


**Figure S3 Predicting mutational effects on thermodynamic folding stability.** Stabilizing-versus-destabilizing classification metrics are computed using $\Delta\Delta G < 0$ (experimental) and $\Delta \log p > 0$ (predicted) as cutoffs for stabilizing mutations. **(A)** Evaluation on the T2837 results: zero-shot models (top) and models fine-tuned on cDNA117k (bottom). **(B)** Evaluation on Megascale test set results: zero-shot models (top) and models fine-tuned on the Megascale training set (bottom). Model names indicate the architecture, the coordinate-noise amplitude used, and when applicable, the fine-tuning dataset (listed after "+"); *Untr.* is short for *Untrained*, indicating models that had no pre-training and were instead only trained on stability effects. Only models trained without coordinate noise are shown; the noise amplitude is indicated within each model name.

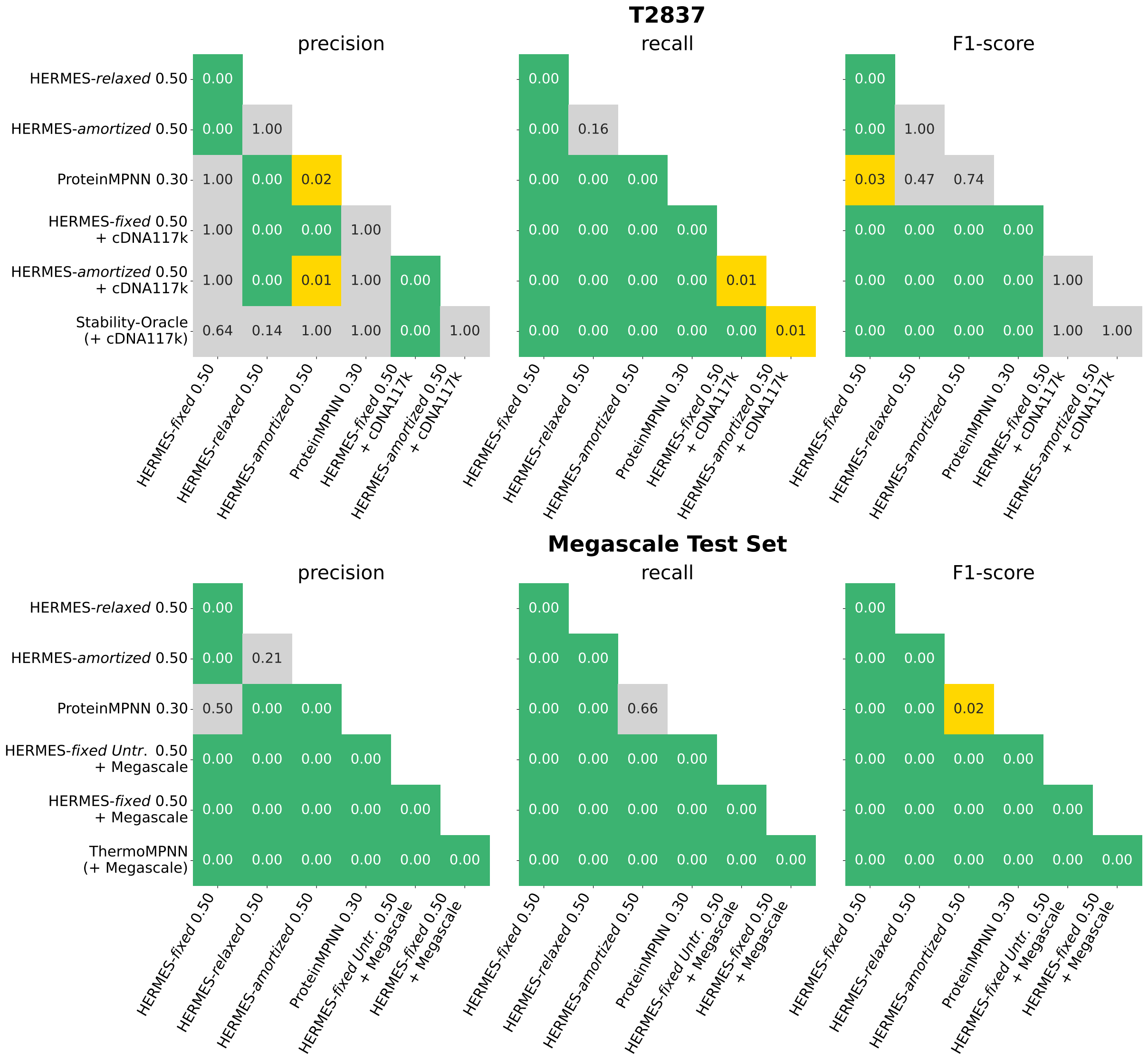


**Figure S4 Statistical significance of model performance differences in identifying stabilizing mutations on the full test set.** Shown are two-tailed p-values for differences in model performance when identifying stabilizing mutations on the full test set. P-values correspond to the performance comparisons shown in Fig. 2. Green indicates strong statistical significance ($p < 0.01$), while yellow indicates weaker significance ($0.01 \leq p < 0.05$). P-values were computed using a permutation test and corrected for multiple comparisons using the Holm-Bonferroni procedure within each performance metric (see Methods for details).

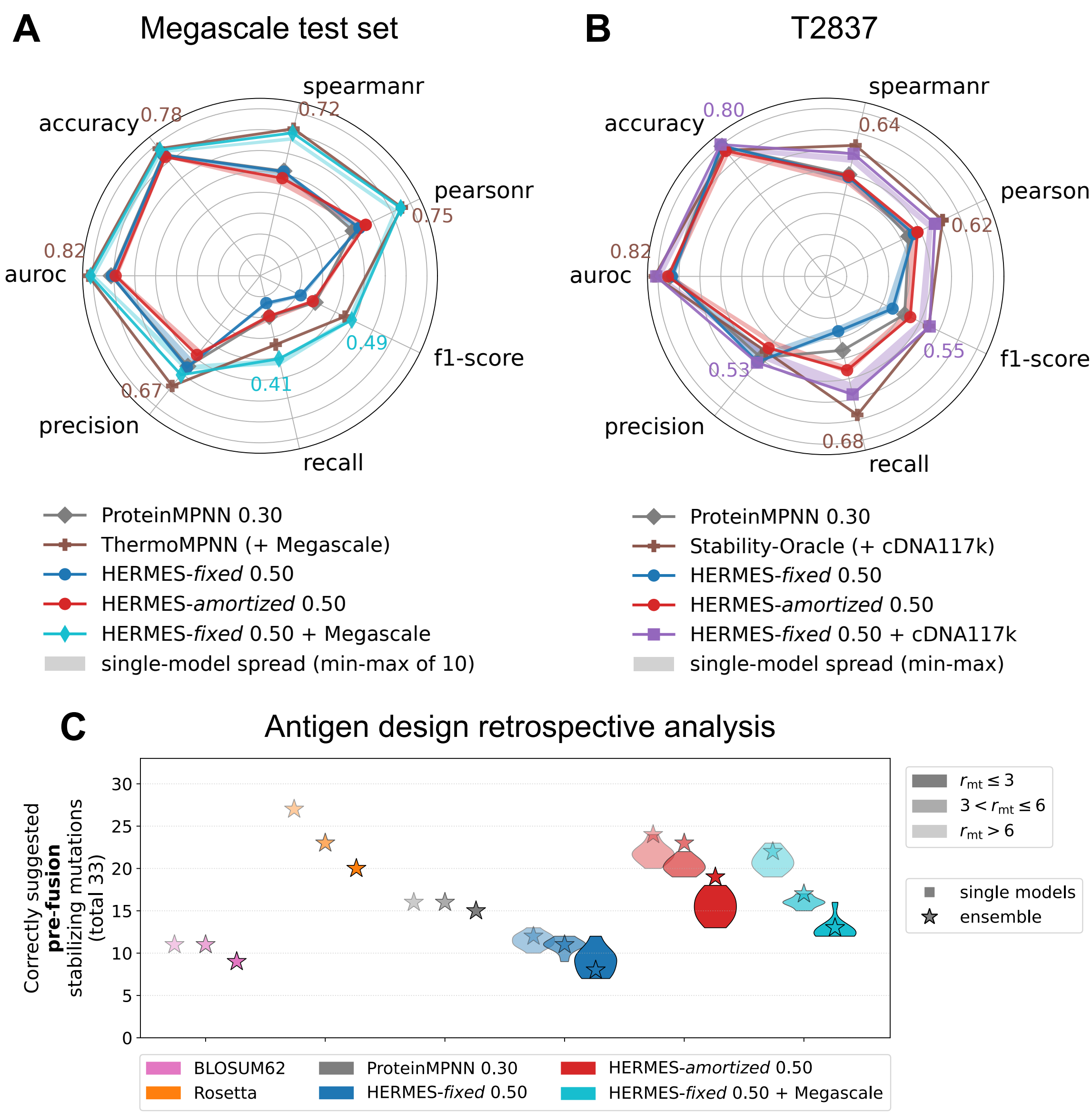


**Figure S5 Performance spread across the 10 individually trained HERMES models in each ensemble. (A)** Performance spread across different performance metrics on the Megascale test set. Solid lines indicate ensemble estimates for HERMES models and single-model estimates for other models (which are *not* ensembled). For HERMES models, shaded regions denote the minimum and maximum metric values across the 10 individual models in each ensemble. **(B)** Performance spread on T2837, reported as in **(A)**. **(C)** Performance spread on the antigen retrospective analysis, where the metric is effectively the recall of individual pre-fusion-stabilizing mutations; see main text and Methods for details.

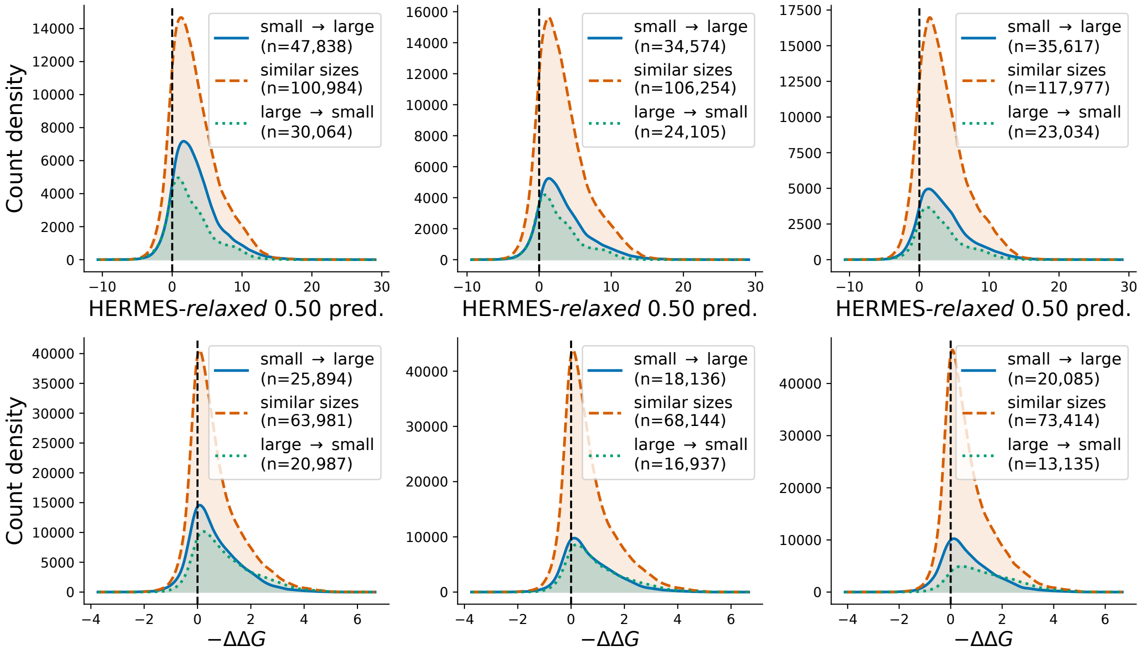


**Figure S6 Distributions of predicted and experimentally measured mutational effects, stratified by amino acid size.** The top row shows the distributions of HERMES-*relaxed* predictions over the distillation set used to fine-tune HERMES into HERMES-*amortized*. The bottom row shows the distributions of experimentally measured $-\Delta\Delta G$ values from the Megascale training set. Each column corresponds to a different splitting of amino acids in size categories, as defined in Fig. 3.

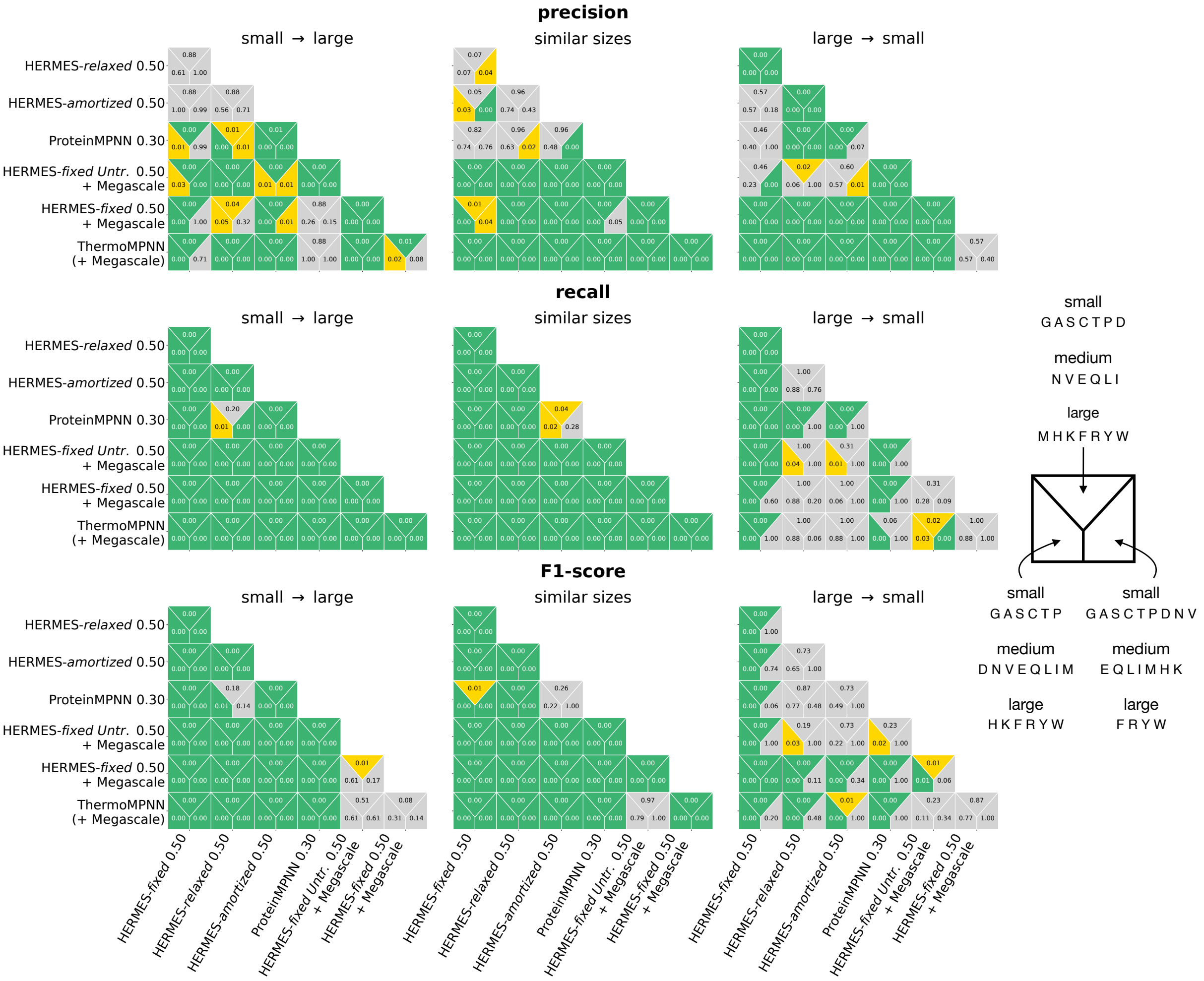


**Figure S7 Statistical significance of model performance differences on stabilizing mutation identification, stratified by amino acid size.** Shown are two-tailed p-values for differences in model performance on identifying stabilizing mutations, across Megascale test subsets defined by the size classes of the wild-type and mutant amino acids. Each model–model comparison (square cell) shows three p-values, corresponding to three distinct groupings of amino acids into size classes (shown on the *right*; same groupings as in Fig. 3for bootstrapping). "Similar sizes" (middle column) refers to the subset of mutations in which the wild-type and mutant amino acids belong to the same size class (grouping together small→small, medium→medium, and large→large). Green indicates strong significance ($p < 0.01$), and yellow moderate significance ($0.01 \leq p < 0.05$). P-values were computed via a permutation test and Holm–Bonferroni-corrected for multiple comparisons within each performance metric (see Methods for details).

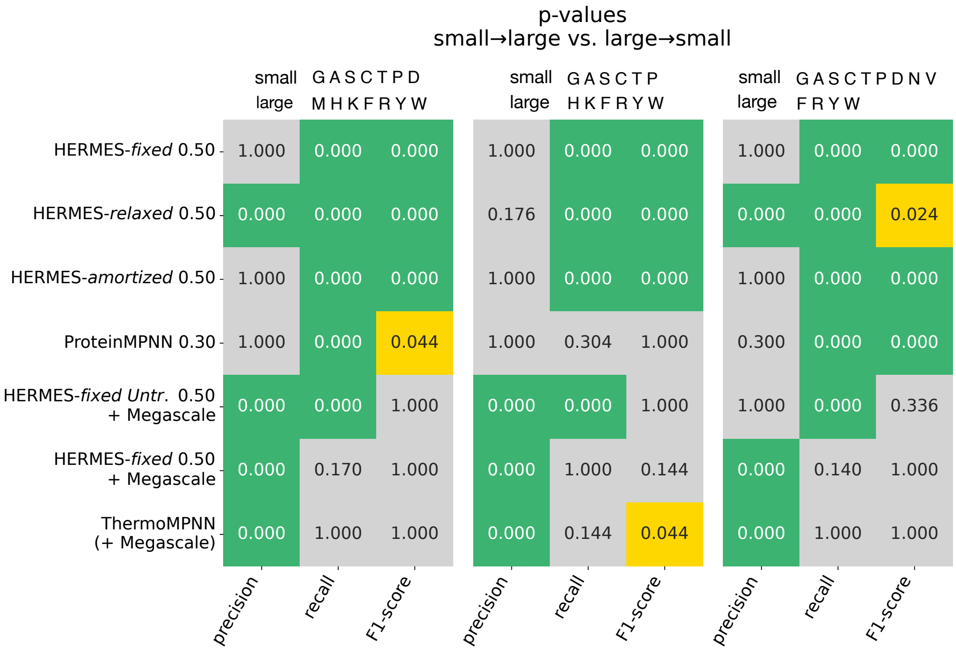


**Figure S8 Statistical significance of within-model performance differences in identifying stabilizing mutations across mutational size classes.** Shown are two-tailed p-values for within-model differences in performance on identifying stabilizing mutations from the small→large versus large→small subsets of the Megascale test set. Each panel corresponds to a distinct grouping of amino acids into size classes, as indicated above the panel (same groupings as in Fig. 3for bootstrapping). Green indicates strong significance ($p < 0.01$), and yellow weaker significance ($0.01 \leq p < 0.05$). P-values were computed via a bootstrap test and Holm–Bonferroni-corrected for multiple comparisons within each performance metric (see Methods for details).

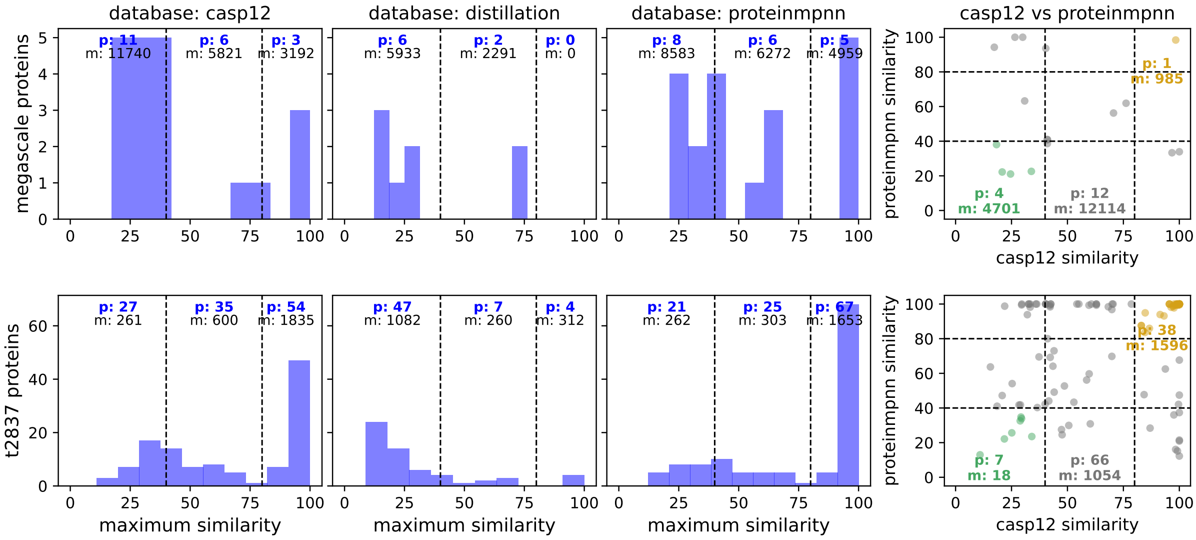


**Figure S9 Sequence similarity (homology) between proteins in the test and pre-training or distillation sets.** We consider the T2837 and Megascale test sets of stability effects (rows). For each, we show the distribution of maximum sequence similarity to the HERMES pre-training set (CASP12), the ProteinMPNN pre-training set, and the HERMES-*amortized* distillation set (columns). Maximum sequence similarity is defined, for each test protein, as its similarity to the closest protein in a given pre-training or distillation set. "Distillation" denotes the set of proteins used to distill HERMES-*relaxed* predictions into HERMES-*fixed* to create HERMES-*amortized*. Each plot is divided at the 40% and 80% similarity cutoffs, with the number of proteins ("p") and number of mutations ("m") shown for each partitioned section. The scatter plots show each test protein's similarity to the ProteinMPNN pre-training set versus that to the HERMES pre-training set (CASP12), with colors indicating regions of low or high similarity for both models. See Methods for details on the similarity computation.

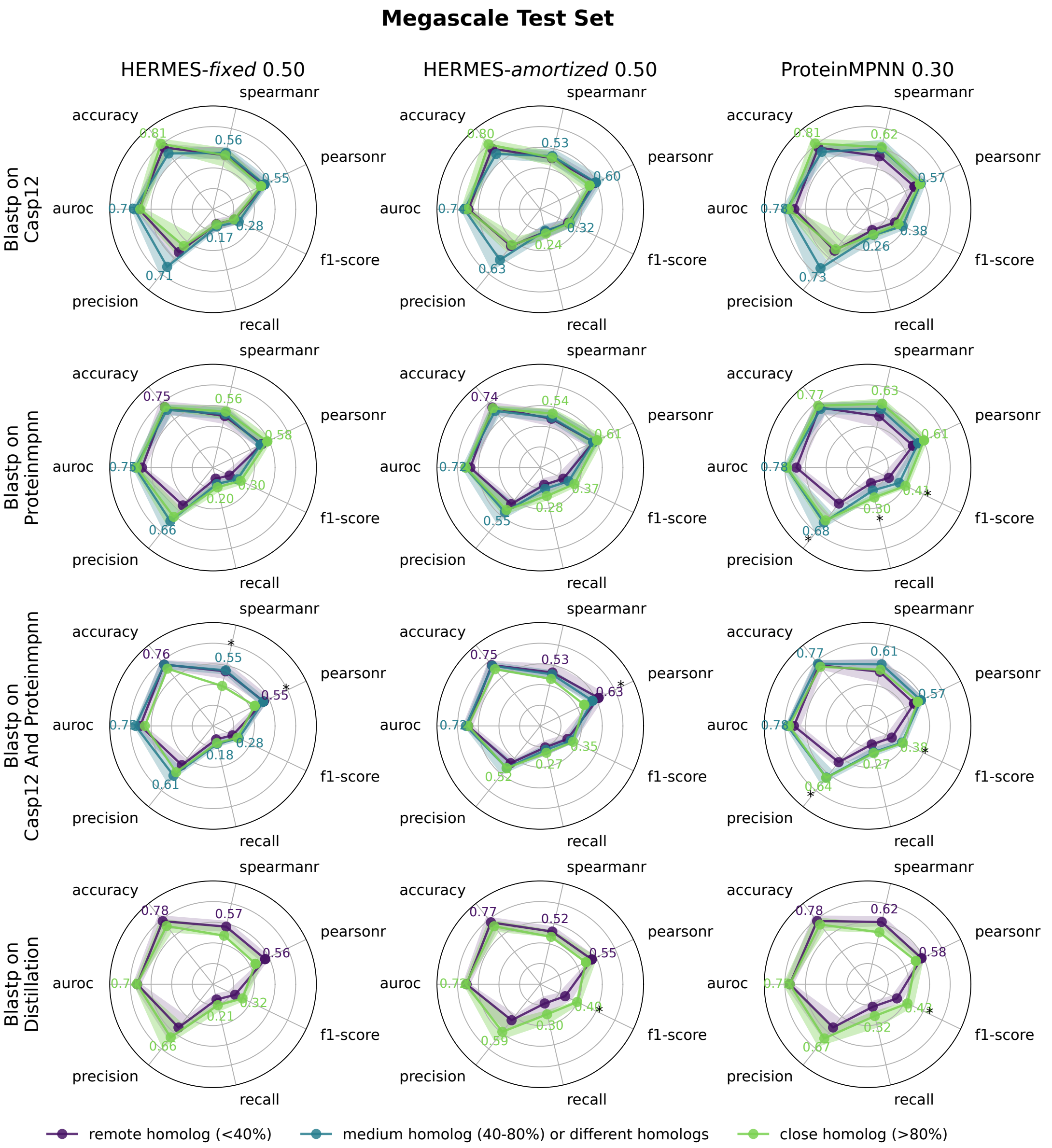


**Figure S10 Predicting mutational effects on folding stability in the Megascale test set, stratified by training-set homology.** Performance on the Megascale test set, split by each test protein's maximum sequence similarity to four reference sets (rows): the HERMES pre-training set, the ProteinMPNN pre-training set, both the HERMES and Protein-MPNN pre-training sets, and the HERMES-*amortized* distillation set. Colors indicate different similarity subsets: remote homologs (< 40% similarity), medium or different homologs (40%–80% similarity or different similarity values in the third row), and close homologs (> 80% similarity). Confidence intervals (shaded areas) are the 5–95 percentiles of bootstrap samples. Stars denote significance of the metric difference between the "remote homolog" and "close homolog" sets. See Methods for details.

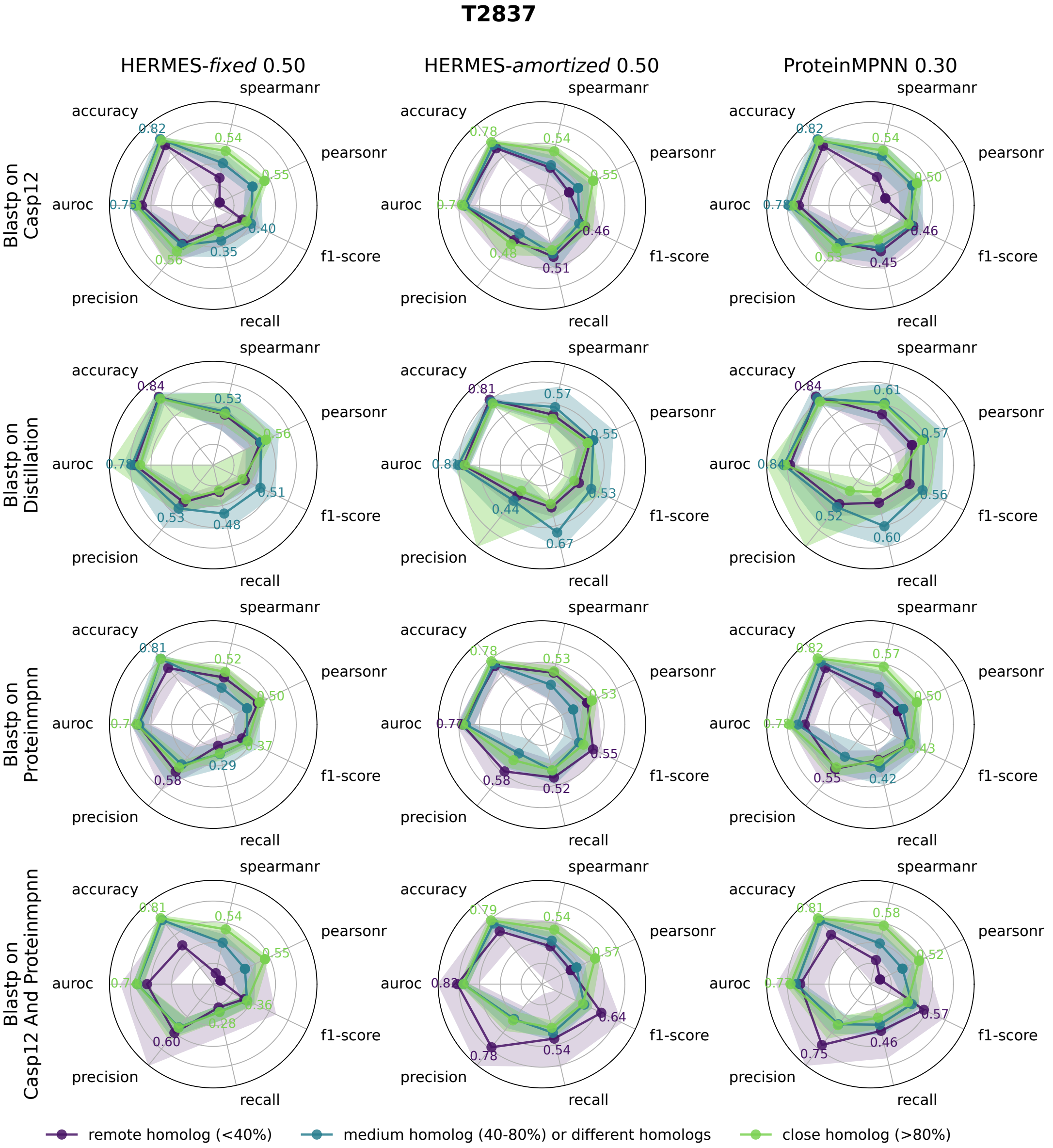


**Figure S11 Predicting mutational effects on folding stability in the T2837 test set, stratified by training-set homology.** Similar to Fig. S10 but for the T2837 test set.

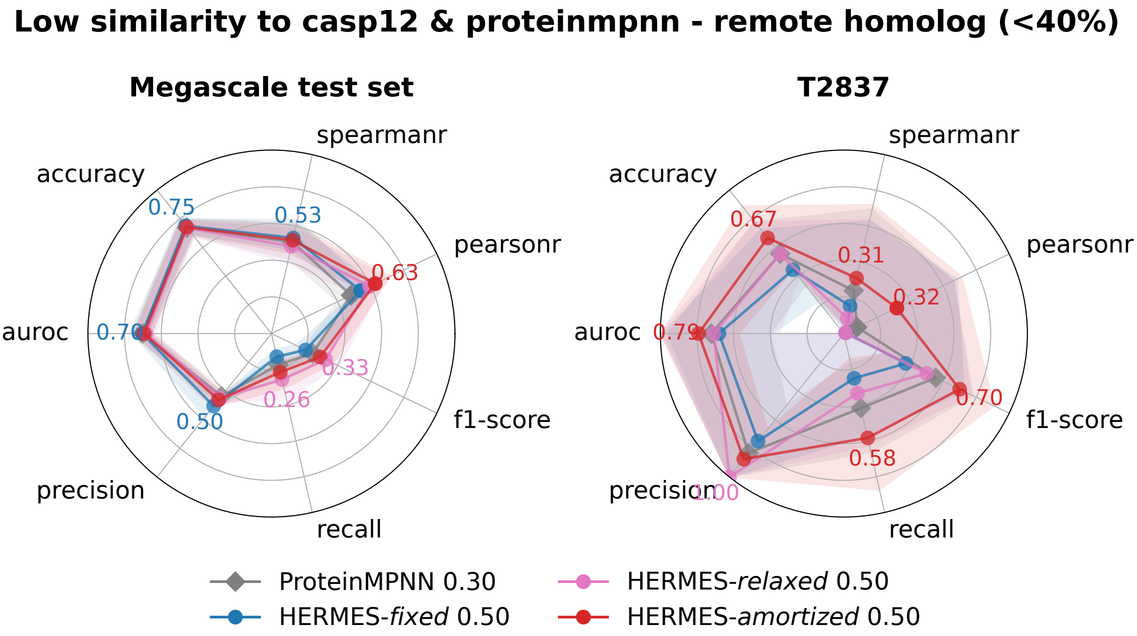


**Figure S12 Predicting stability effects of mutations in proteins with low similarity to pre-training sets.** Accuracy metrics for predicting mutational effects on thermodynamic folding stability, shown for test proteins with low sequence similarity to the pre-training sets of both HERMES models (CASP12) and ProteinMPNN; the test set are remote homologs with $< 40\%$ maximum sequence similarity to both pre-training sets. Confidence intervals (shaded areas) are the 5–95 percentiles of bootstrap samples. Performance metrics do not differ significantly between models after Holm–Bonferroni correction of $p$-values. See Methods for details.

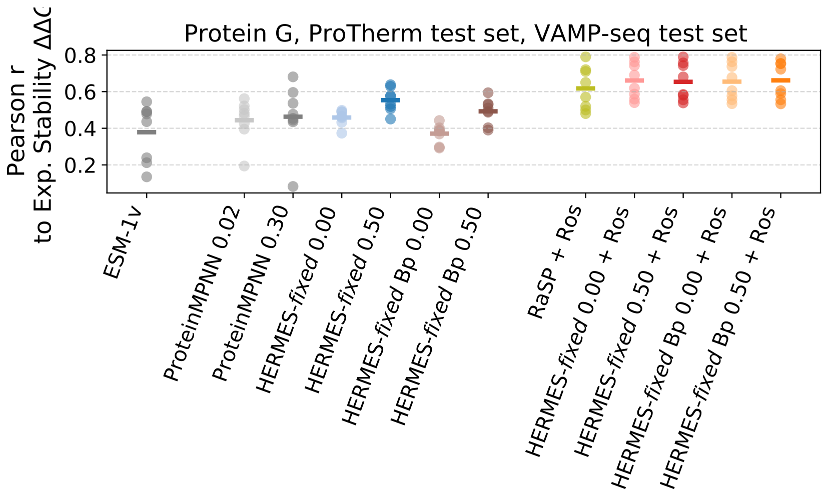


**Figure S13 Pearson correlation between model predictions and experimental stability effects on the RaSP test set (8 proteins) [12].** Each dot represents one protein, and the horizontal bar indicates the mean correlation across proteins. Model labels specify the architecture, the coordinate-noise amplitude, and, when applicable, the fine-tuning dataset (denoted after "+"). "Bp" indicates the use of our open-source Biopython-based protein Pre-processing. See Methods for details on the RaSP dataset.

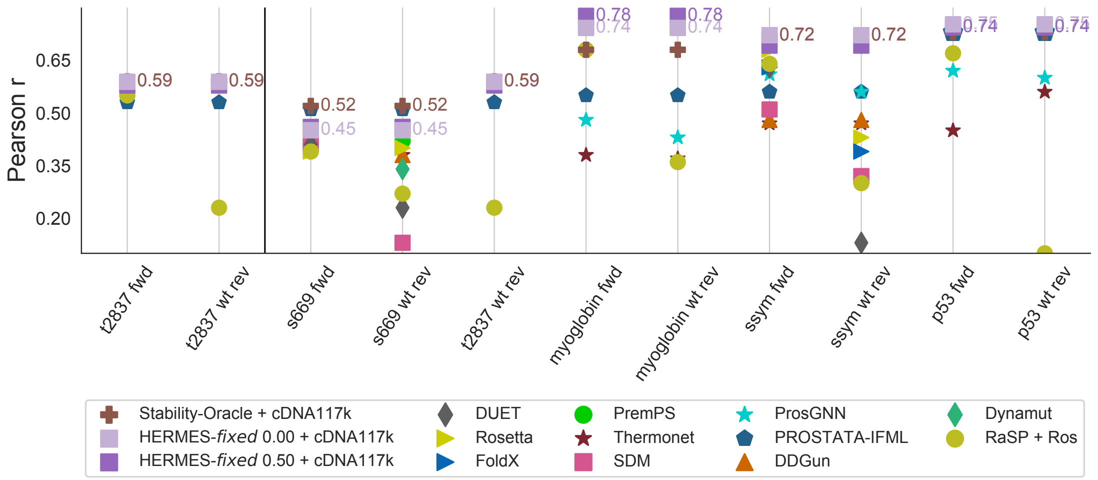


**Figure S14 Pearson correlation between model predictions and experimental stability effects on the T2837 dataset and its subsets.** Pearson correlation values for all models other than HERMES are taken from [13]. This figure closely replicates a figure from ref. [13], with the key difference that predictions for "reverse" mutations are computed here by conditioning on wild-type structures (denoted as "wt rev"). This distinction is made to avoid confusion with "reverse" mutation predictions computed on mutant structures in the Ssym dataset (Fig. 5). For each dataset (x-axis), we denote in text the performance of the HERMES models as well as that of the best-performing model overall.

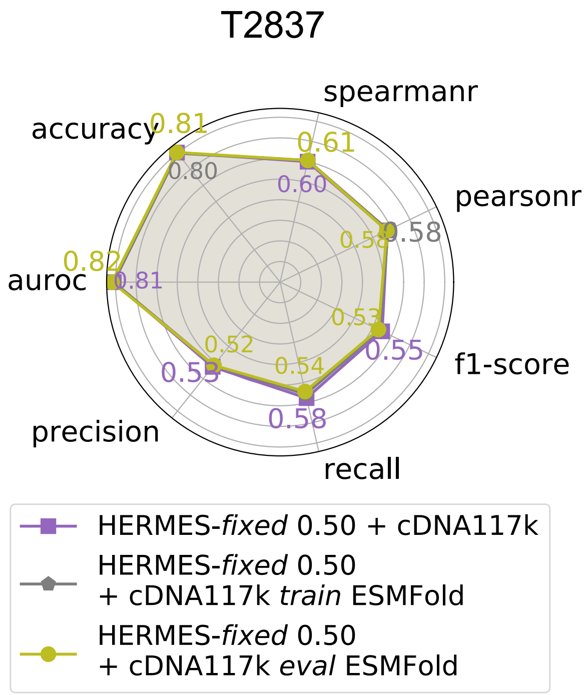


**Figure S15 Predicting mutational effects on thermodynamic stability using ESMfold predicted structures for fine-tuning or testing.** Stabilizing-versus-destabilizing classification metrics are computed using $\Delta\Delta G < 0$ (experimental) and $\Delta \log p > 0$ (predicted) as cutoffs for stabilizing mutations. We report results on the T2837 dataset, after fine-tuning models on cDNA117k. We consider models fine-tuned and evaluated on crystal structures (purple), models fine-tuned on ESMfold predicted structures and evaluated on crystal structures (grey, "*train* ESMfold" in the model name), and models fine-tuned on crystal structures and evaluated on ESMfold predicted structures (olive, "*eval* ESMfold" in the model name).

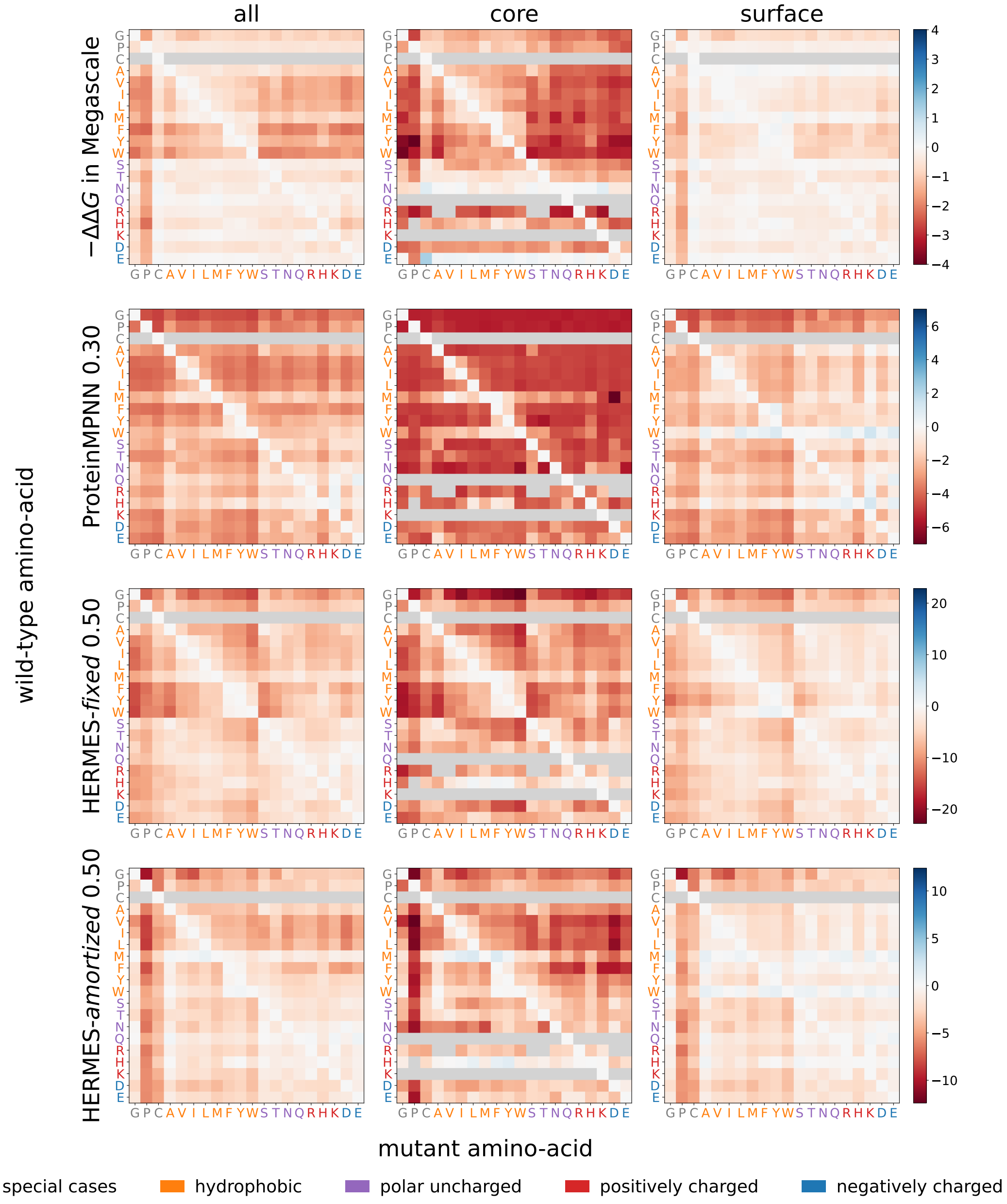


**Figure S16 Amino acid substitution bias matrices for zero-shot models, stratified by protein core and surface residues.** Shown are matrices of amino acid substitution preferences $M^{model}$ (from wild-type (vertical) to mutant (horizontal) amino acids) predicted by different zero-shot models (rows 2–4), computed over sites in the Megascale test set. Columns correspond to all residues (left), core residues (solvent-accessible surface area SASA $< 1$ Å$^2$; center), and surface residues (SASA $> 3$ Å$^2$; right). The first row shows the corresponding matrices of experimental $-\Delta\Delta G$ values, averaged over the same wild-type-to-mutant pairs in the Megascale test set. Amino acids symbols are colored according to their physico-chemical properties.

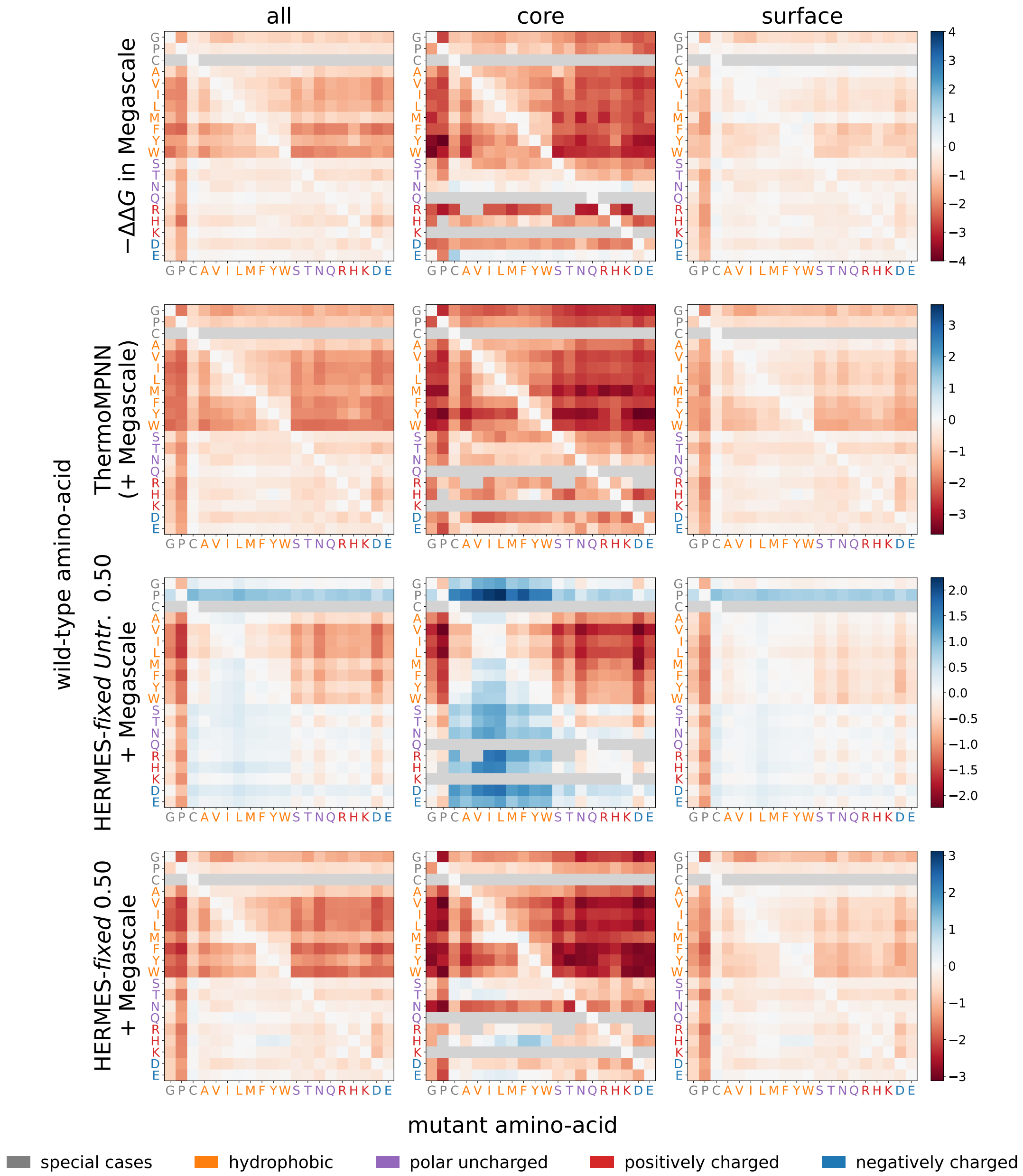


**Figure S17 Amino acid substitution bias matrices for fine-tuned models, stratified by protein core and surface residues.** Similar to Fig. S16 but for models fine-tuned on the Megascale training set.

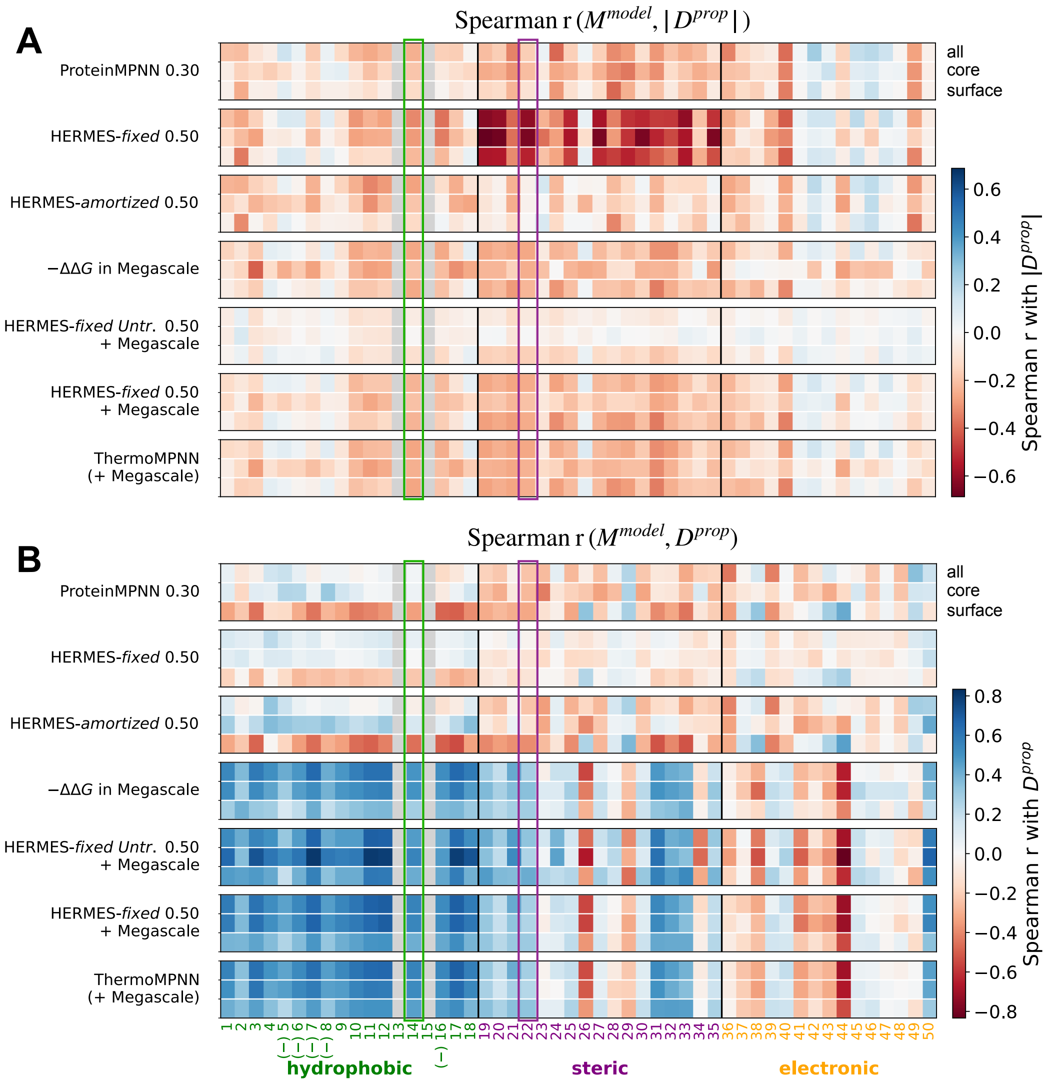


**Figure S18 Correlations between models' substitution bias and changes in amino acid properties.** Spearman $r$ are computed between different models' substitution bias matrices $M^{model}$ (Figs. S16, S17) and the matrices of amino acid property change upon substitution: (A) the absolute change $|D^{prop}|$, and (B) the signed change $D^{prop}$. The 50 properties, and their corresponding property-change matrices, are constructed from Table S3, whose values are taken from ref. [8]. The two properties shown in the main text (Fig. 4) are highlighted (14: transfer $\Delta G$ from an organic solvent to water; 22: van der Waals volume). Properties 5, 6, 7, 8, and 16 are negated so that, within the "hydrophobic" category, a positive change consistently corresponds to increasing hydrophobicity. Properties 13 and 15 are left empty because missing property values introduce NaNs in $|D^{prop}|$ and $D^{prop}$.

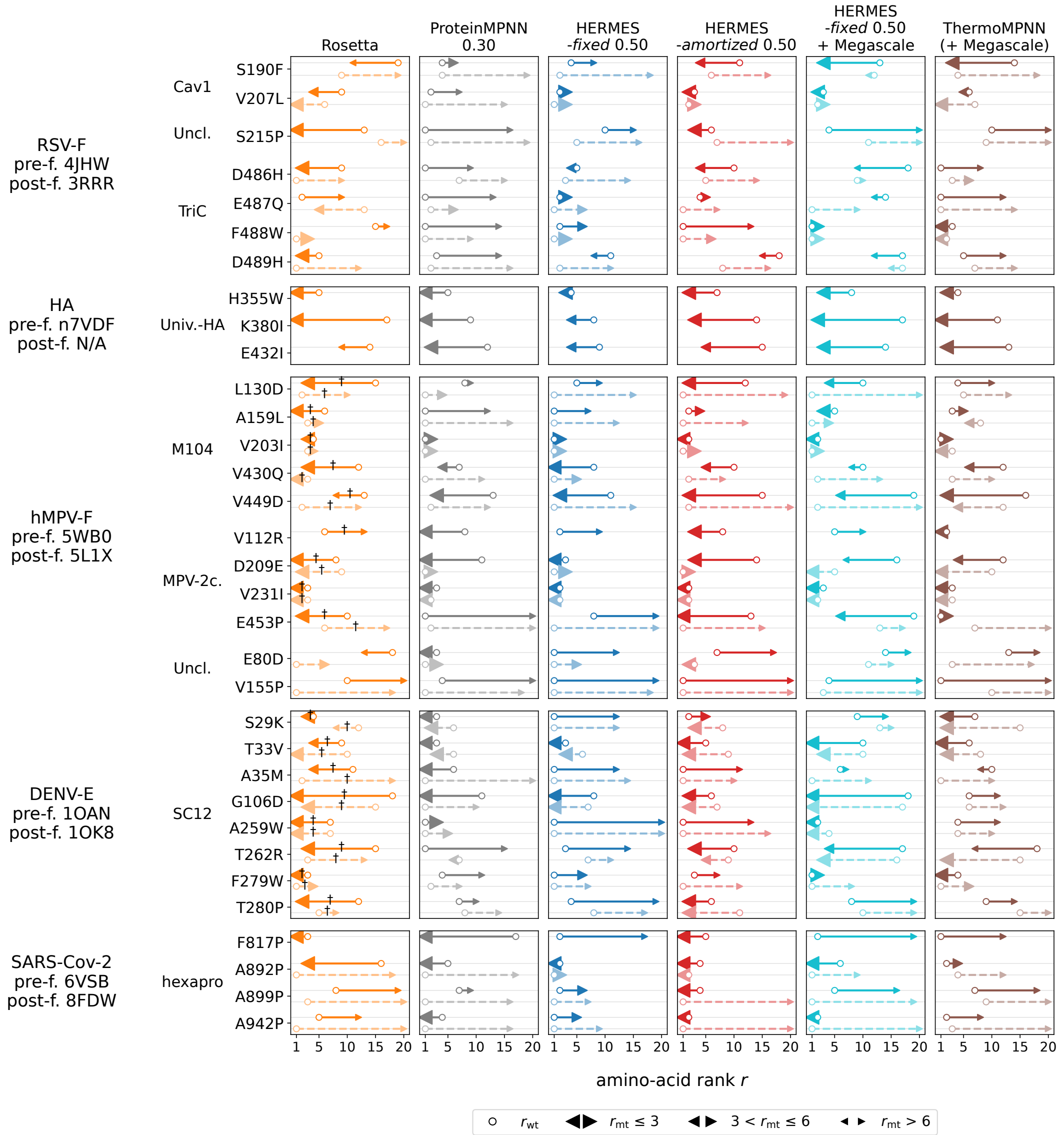


**Figure S19 Recovering experimentally verified antigen-stabilizing mutations.** For 33 previously reported antigen-stabilizing mutations, we report the predictions of 6 models on the relative preference of amino acids in the sites where the mutations are observed. These preferences are reported in the form of amino acid ranks of the wild-type ($r_{\mathrm{wt}}$) and mutant ($r_{\mathrm{mt}}$) amino acids. Arrows depict the change in predicted rank from the wild type (open circle) to the stabilizing mutant (arrow tip); arrow size indicates whether the mutant ranks in the top 3 (large), ranks 4-6 (medium), or ranks $> 6$ (small). Solid-colored arrows indicate predictions made on the pre-fusion structures, whereas light-colored arrows indicate predictions on the post-fusion structures; PDB ids of the structures used for each antigen are shown on the left. The † symbols mark mutations originally proposed as stabilizing by Rosetta. "Univ.-HA" stands for "Universal-HA"; "MPV-2c." stands for MPV-2cREKR; "Uncl." stands for "Uncleaved Prefusion-Closed".

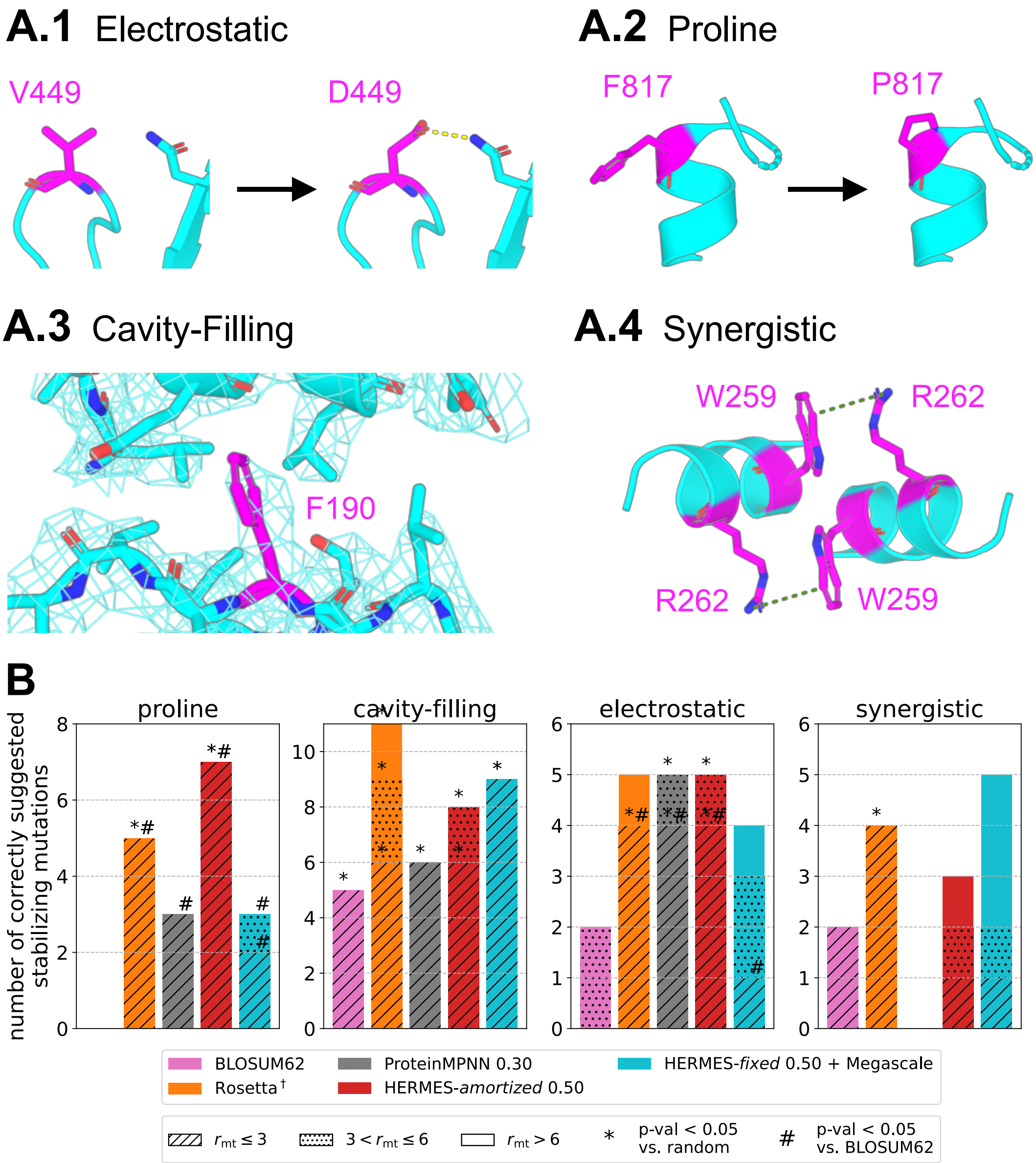


**Figure S20 Model ability to recover different mutation patterns that stabilize antigens. (A)** Representative examples of mutation types commonly seen in antigen stabilization [11], and analyzed in this study. The mutated residue(s) are shown in magenta. **(A.1)** *Electrostatic mutation* in hMPV-F: wild-type structure (PDB ID 5WB0, left), in-silico mutant generated with PyMOL's mutagenesis wizard; right. The introduced Aspartic acid forms an electrostatic interaction with a nearby Asparagine (dashed yellow line). **(A.2)** *Proline mutation* at the N-terminal of a $\alpha$-helix cap in SARS-CoV-2 spike: wild-type structure (PDB ID 6VSB; left), in-silico mutant generated with PyMOL's mutagenesis wizard; right. **(A.3)** *Cavity-filling mutation* in RSV-F: mutant structure (PDB ID 4MMS). A bulky hydrophobic substitution packs a previously underfilled region: the $2Fo$-$Fc$ electron density map is shown as a thin mesh. **(A.4)** *Synergistic dimer-stabilizing mutations* in DENV-E: A259W and T262R in the dimer mutant structure (PDB ID 6WY1). introduced in both chains, create a stabilizing cation-$\pi$ interaction (dashed green lines). **(B)** Number of stabilizing mutations recovered by each model, stratified by mutation class, reported as in Fig. 6A.

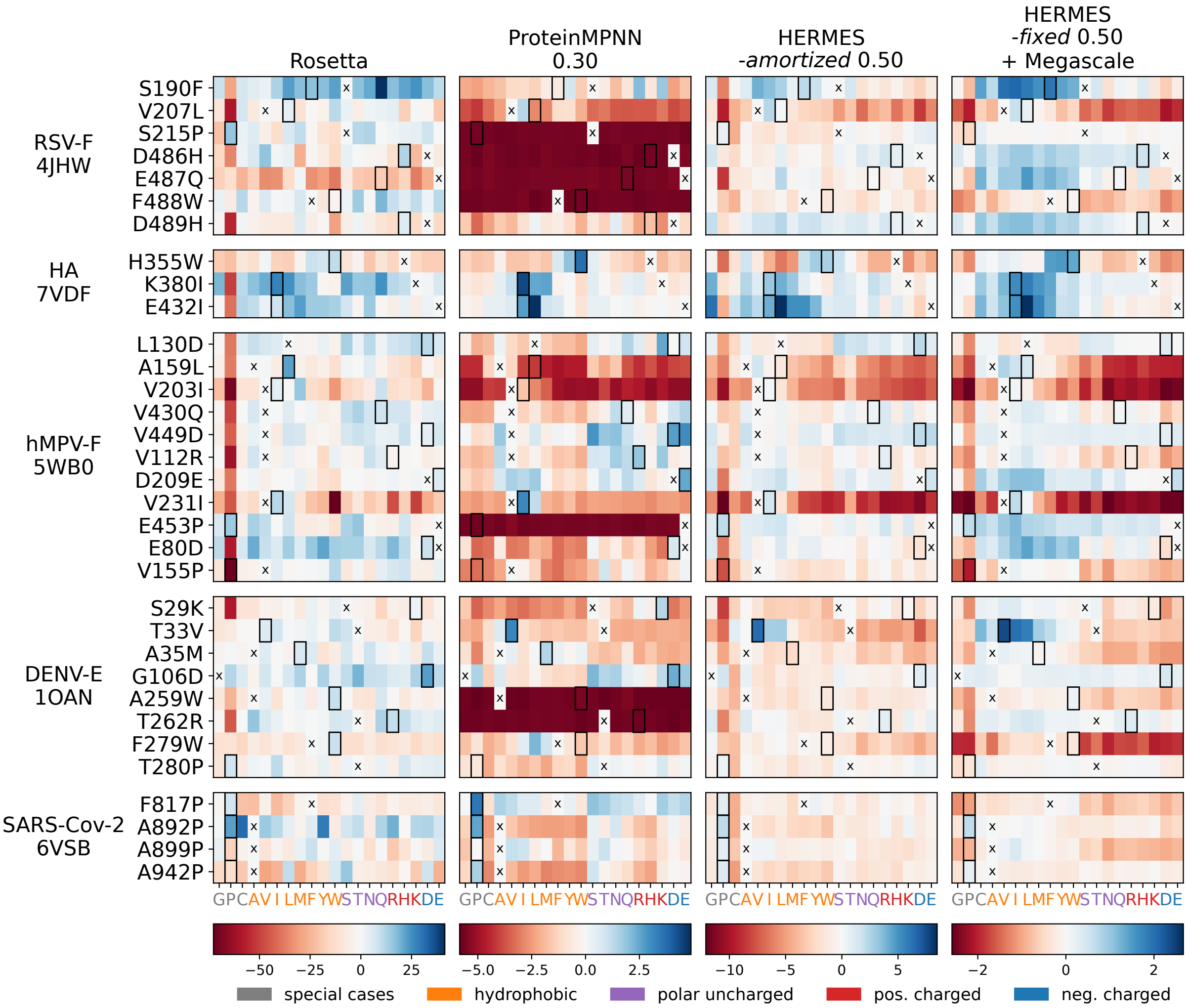


**Figure S21 Model predictions for amino acid preferences at sites with known antigen-stabilizing mutations.** Predictions from different models (columns) are shown for sites with known antigen-stabilizing mutations specified in Fig. 6and Table S4. For each antigen (rows), predictions are computed using the protein structure corresponding to the PDB ID indicated on the left. Predictions are reported as changes in Rosetta Energy Units (REU) for Rosetta, and $\Delta \log p$ for ProteinMPNN and HERMES models. Wild-type amino acids are marked with centered crosses, while stabilizing mutant amino acids are indicated by dark borders. Amino acids are grouped by broad biochemical class (see legend at the bottom) and, within each class, ordered by increasing size (number of atoms).

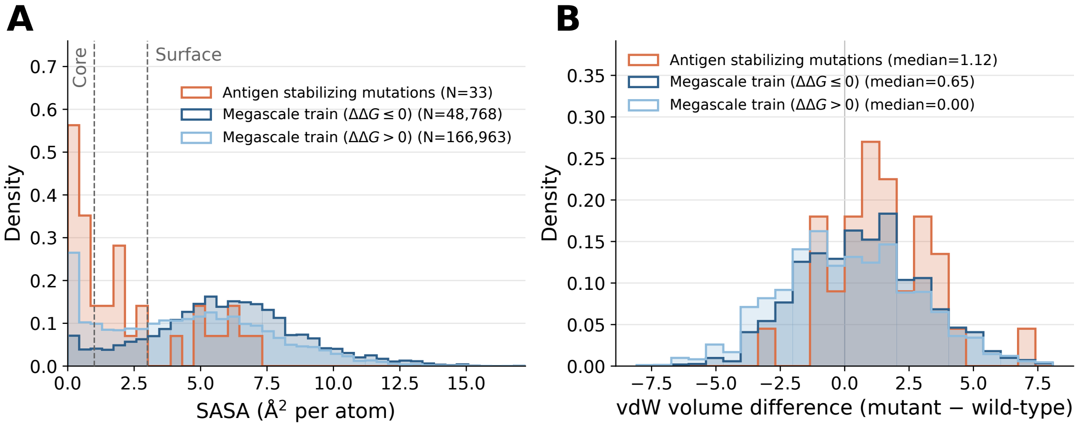


**Figure S22 Physico-chemical properties of antigen-stabilizing mutations and of stabilizing and destabilizing substitutions in the Megascale training set.** Distributions of **(A)** the mean per-atom solvent-accessible surface area (SASA) of the wild-type residue and **(B)** the change in van der Waals volume upon substitution, shown for three classes of substitutions: (i) the 33 verified antigen-stabilizing mutations from Fig. 6, (ii) the stabilizing ($\Delta\Delta G < 0$) mutations, and (iii) the destabilizing ($\Delta\Delta G > 0$) mutations in the Megascale training set. cDNA117k set it is derived from the same data as the Megascale training set [14], so we expect their statistics to be similar. The antigen-stabilizing mutations considered in this study are relatively more enriched in the core (smaller SASA) than the stabilizing mutations used to fine-tune the models. No notable difference in the change of amino acid size (van der Waals volume) is seen between the antigen-stabilizing mutations and the stabilizing mutations in the fine-tuning set.

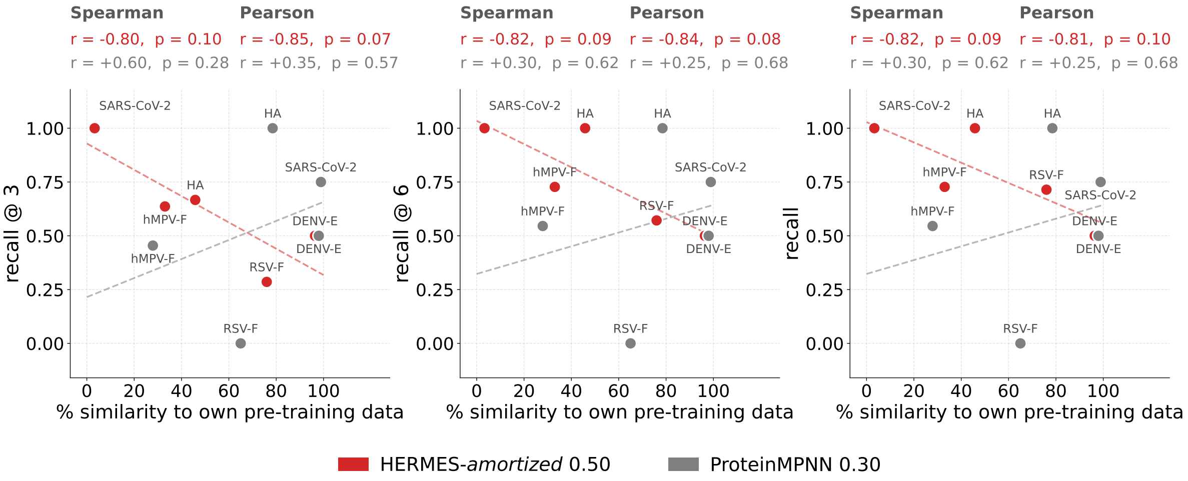


**Figure S23 Impact of sequence similarity to the pre-training set on model performance for identifying antigen-stabilizing mutations.** Each model's recall of antigen-stabilizing mutations is plotted against each antigen's maximum sequence similarity to the pre-training set; for HERMES-*amortized*, the pre-training set is CASP12 (see Table S7 for the raw similarity values). Recall@3and Recall@6 (left and center panels) denote the fraction of stabilizing mutations predicted with $r_{\rm mt} < r_{\rm wt}$ and $r_{\rm mt} \leq 3$ or $r_{\rm mt} \leq 6$, respectively, while Recall (right panel) denotes the fraction predicted with $r_{\rm mt} < r_{\rm wt}$ alone. Dotted lines denote the line of best fit (ordinary least-squares estimate). P-values are the raw $t$-approximation values (the default in `scipy.stats`).

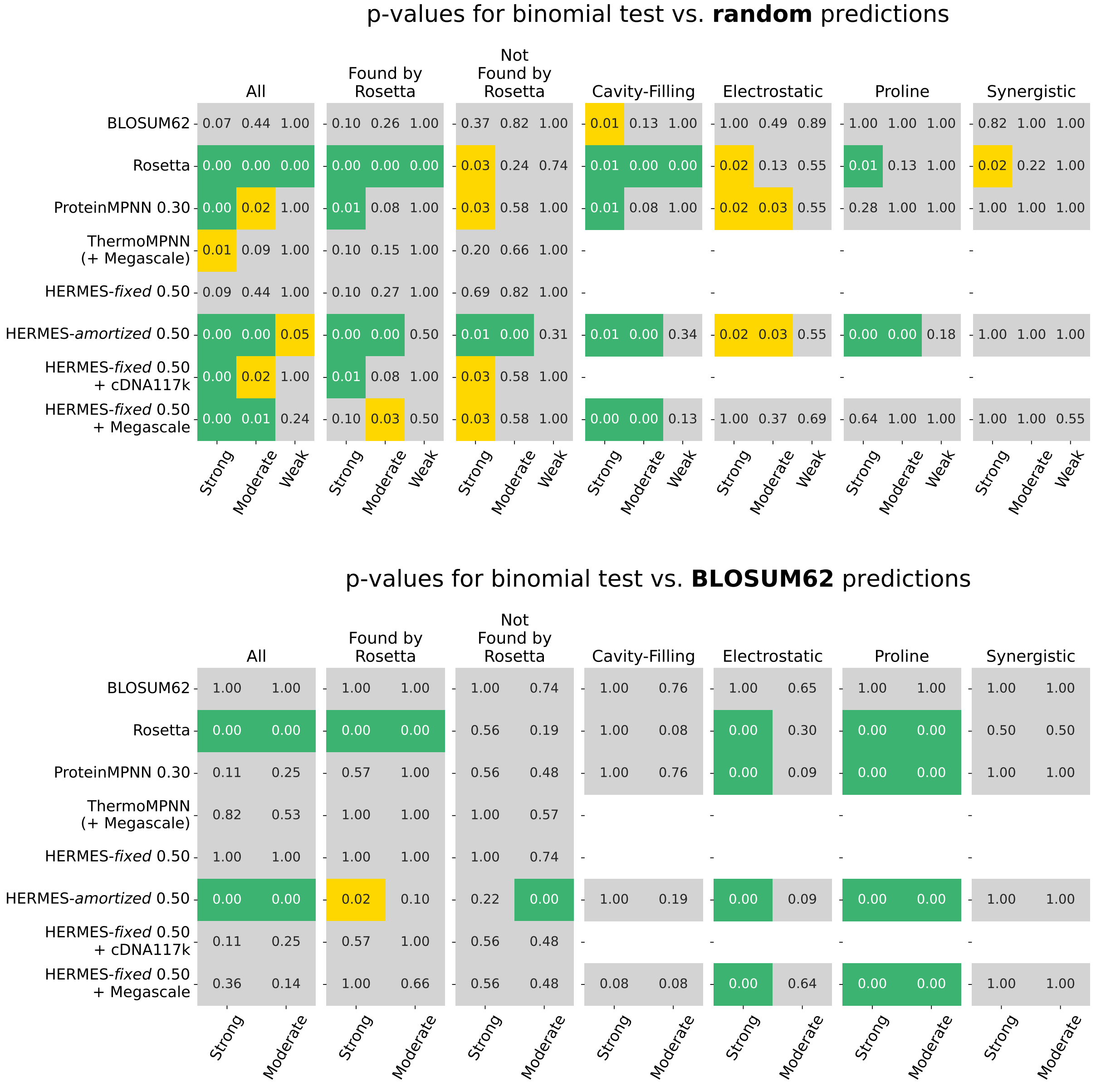


**Figure S24 Statistical significance for the number of retrieved antigen-stabilizing mutations.** Shown are p-values from binomial tests comparing the number of antigen-stabilizing mutations retrieved by each model (rows) against random expectation (top) and the BLOSUM62 predictions (bottom). These significance tests correspond to the results reported in Figs. 6A, S20 and Table S4; see Methods for details of p-value computation.

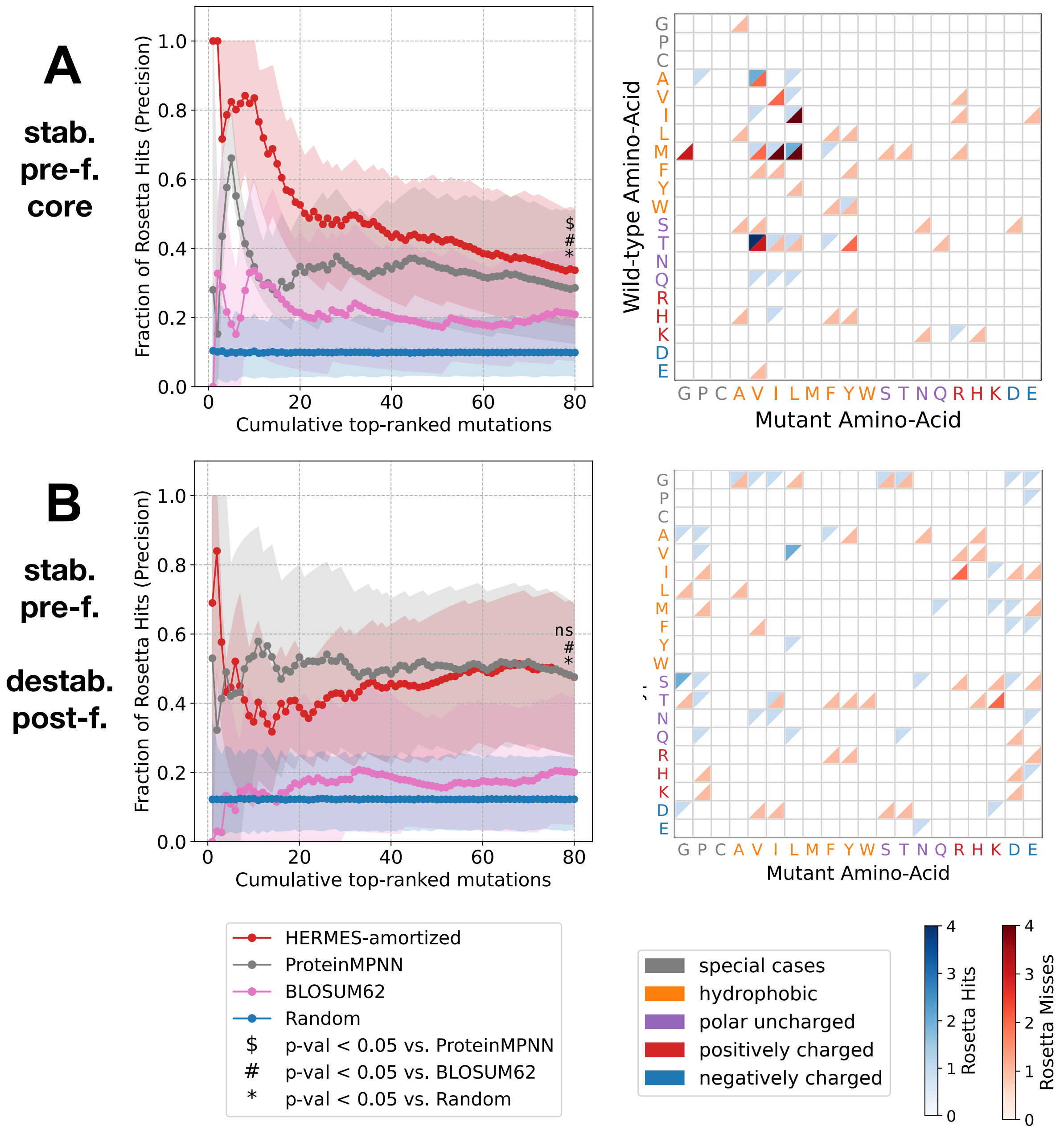


**Figure S25 Computational workflow with HERMES for designing a mutational library for antigen stabilization in DENV-E.** Similar to Fig. 7 A, B but for DENV-E.

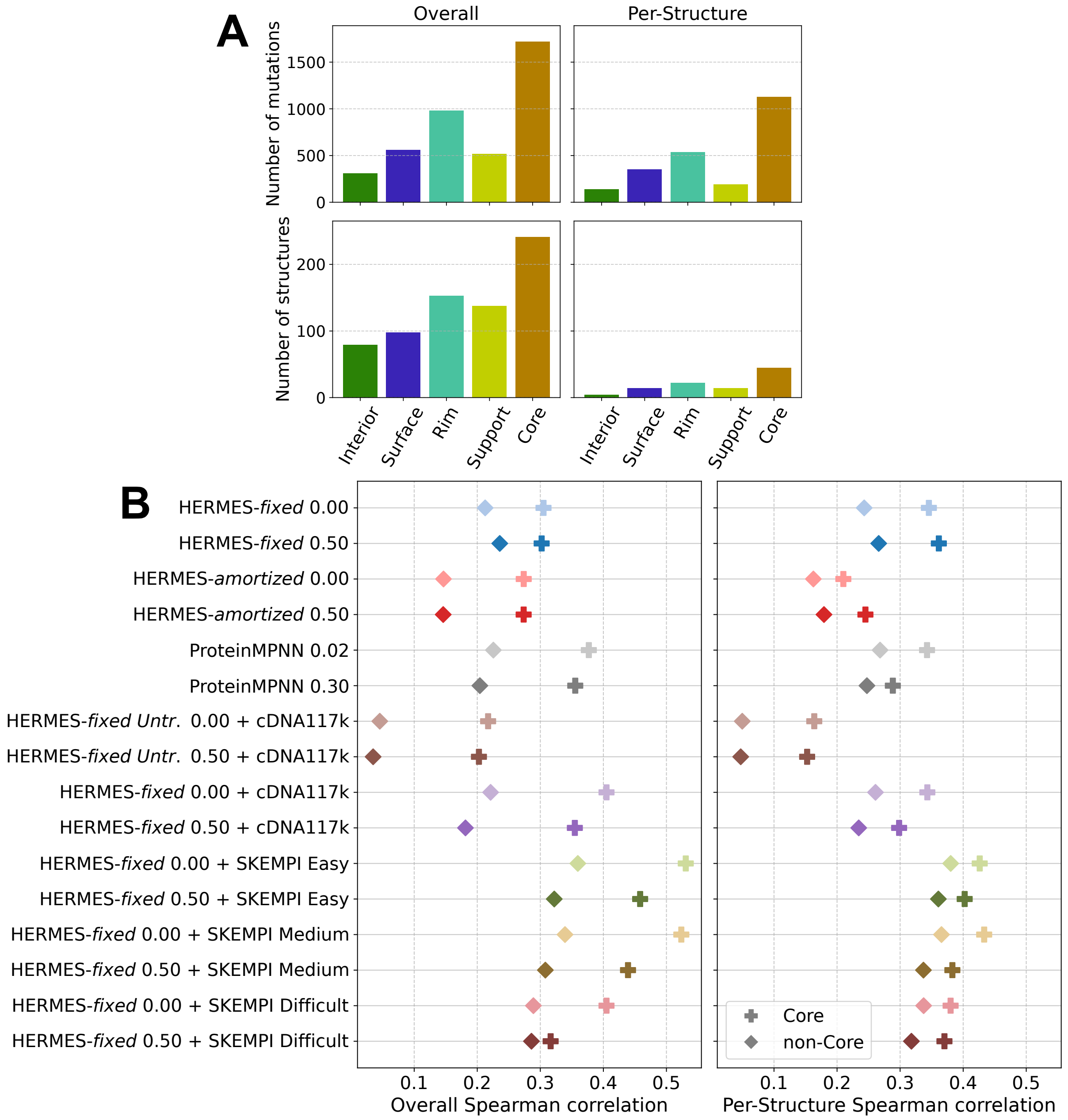


**Figure S26 Model performance for predicting the effect of mutations on binding affinity, stratified by structural context. (A)** Number of mutations and of protein structures from which the mutations are derived in the SKEMPI v2.0 dataset, stratified by structural context as defined in ref. [15]. **(B)** Spearman correlation between model predictions and experimentally measured binding affinity, stratified by mutations at the interface core versus mutations in all other contexts (symbols).

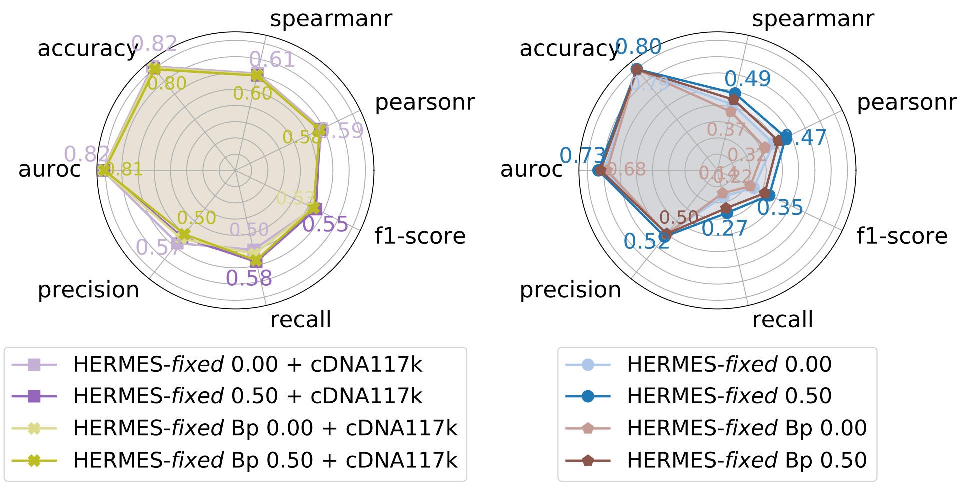


**Figure S27 Comparison of PyRosetta and Biopython Pre-processing pipelines for predicting mutation stability effects on T2837.** Classification accuracy metrics, analogous to those shown in Fig. 2, are reported for fine-tuned models (left) and zero-shot models (right). In each case, models trained using PyRosetta-based Pre-processing are compared with those using Biopython-based Pre-processing (denoted by "Bp" in the model name). Model labels specify the architecture, the coordinate-noise amplitude, and, when applicable, the fine-tuning dataset (listed after "+"). Consistent with results on the RaSP dataset (Fig. S13), Biopython-Pre-processed models show slightly reduced performance relative to PyRosetta-Pre-processed models; however, this difference becomes statistically insignificant after fine-tuning.

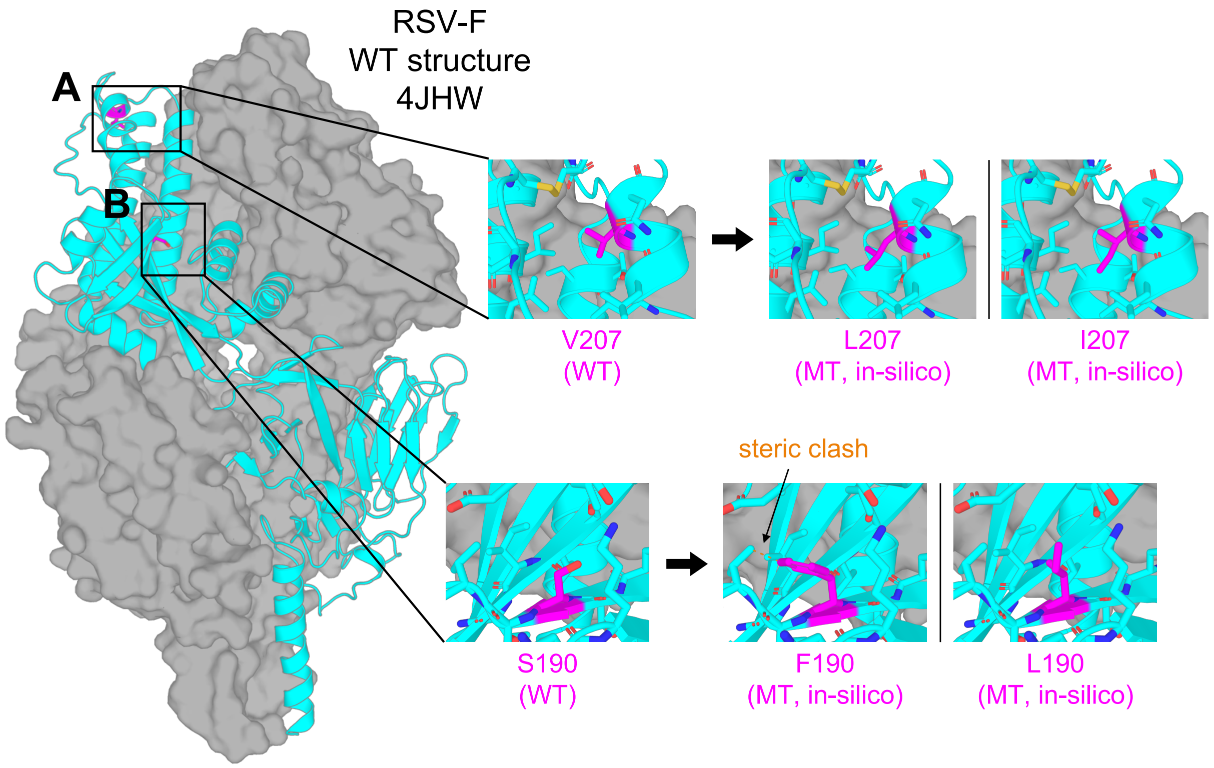


**Figure S28 Structure of the wild-type pre-fusion RSV-F antigen highlighting observed and candidate antigen-stabilizing mutations.** The wild-type structure is shown on the left (PDB ID: 4JHW). A single protomer of the trimer is displayed as a cyan cartoon, with the remaining protomers shown as a grey surface. WT denotes wild type and MT denotes mutant. Mutations labeled as "in silico" were introduced using PyMOL's Mutagenesis Wizard starting from the wild-type structure. Steric clashes (orange dashed lines) were identified using PyMOL's "find clashes" command, and polar contacts (yellow dashed lines) were identified using the corresponding PyMOL command. **(A)** The L207 mutant has been experimentally shown to stabilize the pre-fusion conformation [1] and appears to enhance intraprotomer packing. We speculate that the I207 mutant, which HERMES-*amortized* predicts to have a comparable ranking to L207 in Fig. S21, would pack similarly and may therefore represent an additional stabilizing mutation worth screening. **(B)** Mutant F190 is observed to be stabilizing [1], though it appears to slightly over-pack the region. We speculate that L190, which HERMES-*amortized* predicts to have a comparable ranking to F190 in Fig. S21, would provide a similarly stabilizing effect without over-packing the region.

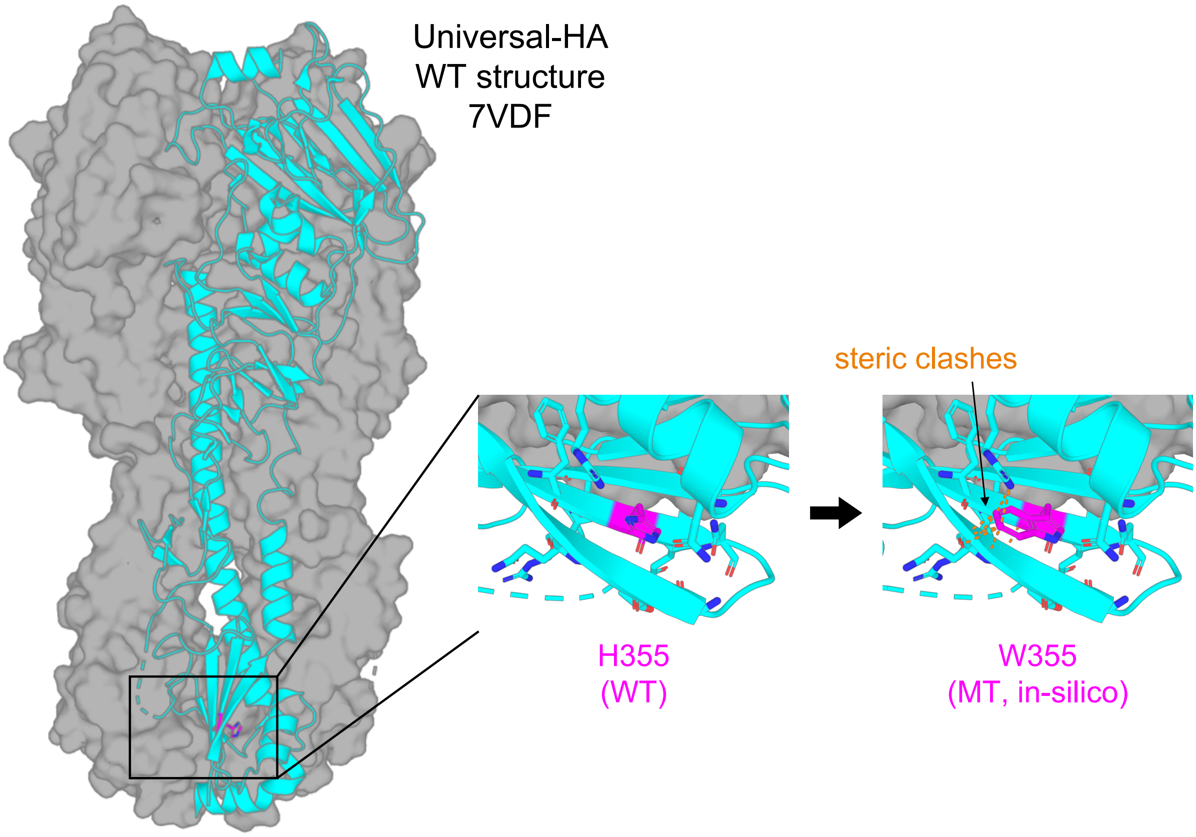


**Figure S29 Structure of the wild-type pre-fusion Universal-HA antigen, with highlighted observed and candidate antigen-stabilizing mutations.** The wild-type structure is shown on the left (PDB ID 7VDF). A single protomer of the trimer is shown as cyan cartoon; the other copies are shown as grey surface. WT denotes wild type and MT denotes mutant. Mutations labeled as "in silico" were introduced using PyMOL's Mutagenesis Wizard starting from the wild-type structure. Steric clashes (orange dashed lines) were identified using PyMOL's "find clashes" command, and polar contacts (yellow dashed lines) were identified using the corresponding PyMOL command. We highlight the H355W mutation which, despite stabilizing the pre-fusion conformation, exhibits substantial steric clashes in the wild-type structural context.

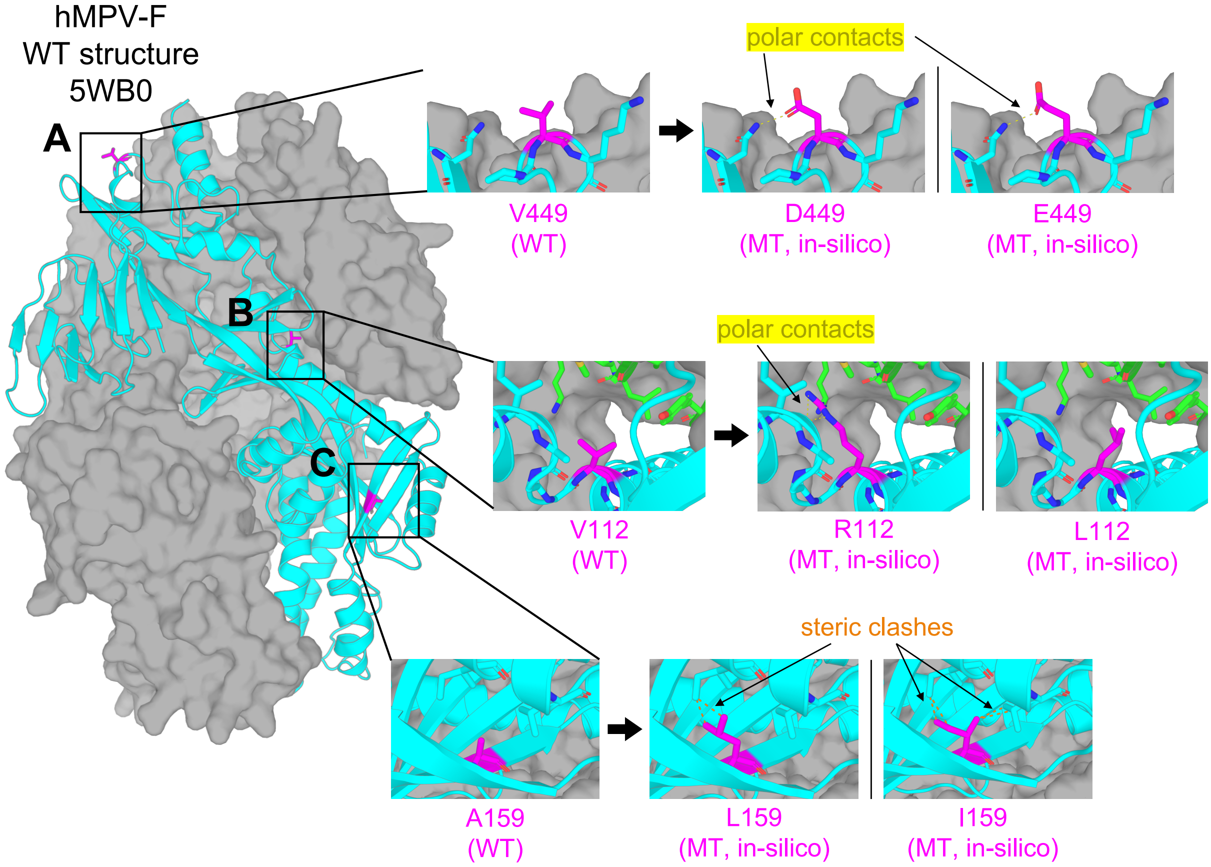


**Figure S30 Structure of the wild-type pre-fusion hMPV-F antigen, with highlighted observed and candidate antigen-stabilizing mutations.** The wild-type structure is shown on the left (PDB ID 5WB0). A single protomer of the trimer is shown as cyan cartoon; the other copies are shown as grey surface. WT denotes wild type and MT denotes mutant. Mutations labeled as "in silico" were introduced using PyMOL's Mutagenesis Wizard starting from the wild-type structure. Steric clashes (orange dashed lines) were identified using PyMOL's "find clashes" command, and polar contacts (yellow dashed lines) were identified using the corresponding PyMOL command. **(A)** The proposed V449D mutation is highlighted, which likely stabilizes the complex by removing a surface-exposed hydrophobic residue and introducing an intraprotomer polar contact. We speculate that substitution with glutamic acid, which HERMES-*amortized* predicts to have a comparable ranking to D449 in Fig. S21, would produce a similar stabilizing effect. **(B)** Introduction of an Arginine in place of a Valine at position 112 likely introduces an intraprotomer polar contact, as well as packing against the adjacent protomer (shown in green); we believe L112, which HERMES-*amortized* predicts to have a comparable ranking to R112 in Fig. S21, would also pack against the adjacent protomer better than Valine. **(C)** A159L stabilizes the complex likely due to a cavity-filling effect, albeit necessitating nearby I137 to adopt a different rotamer; we believe I159, which HERMES-*amortized* predicts to have a comparable ranking to L159 in Fig. S21, would also fulfill a similar role on the condition of L141 also adopting a different rotamer.

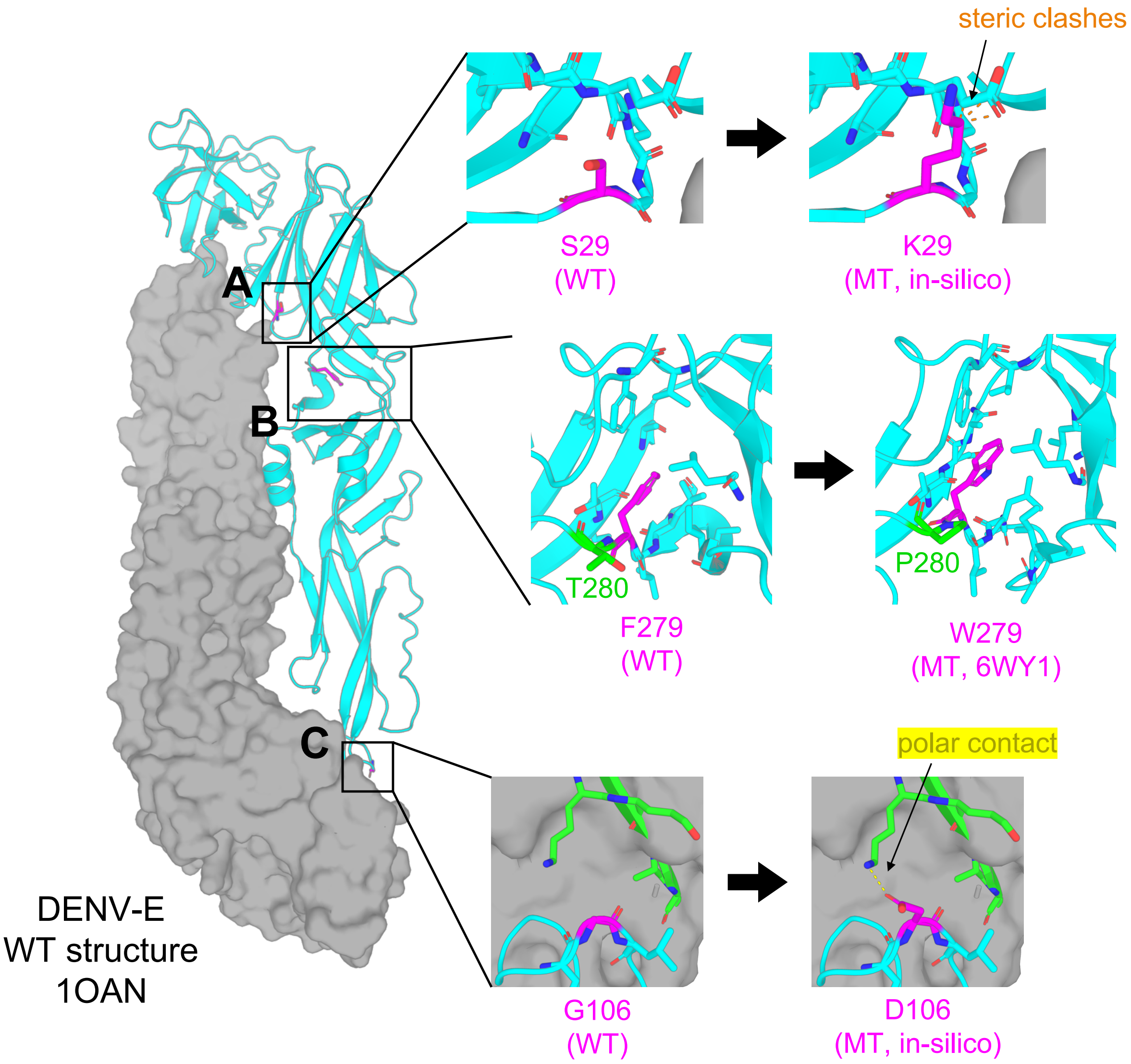


**Figure S31 Structure of wild-type pre-fusion DENV-E antigen, with highlighted observed and candidate mutations.** The wild-type structure is shown on the left (PDB ID 1OAN). A single protomer of the trimer is shown as cyan cartoon; the other copies are shown as grey surface. WT denotes wild type and MT denotes mutant. Mutations labeled as "in silico" were introduced using PyMOL's Mutagenesis Wizard starting from the wild-type structure. Steric clashes (orange dashed lines) were identified using PyMOL's "find clashes" command, and polar contacts (yellow dashed lines) were identified using the corresponding PyMOL command. **(A)** The S29K mutation is experimentally observed to be stabilizing [4], despite appearing to introduce substantial steric clashes when inspected in the wild-type structure, suggesting that stabilization requires a shift in backbone conformation. **(B)** The F279W mutation is experimentally observed to be stabilizing [4], likely by filling an under-packed cavity. A substantial backbone rearrangement is also observed in the mutant structure (residues 269-281), potentially facilitated by the T280P mutation. **(C)** The G106D mutation is experimentally observed to be stabilizing [4], likely through the introduction of a polar contact with the adjacent protomer (shown in green) and/or interactions with surrounding water molecules.
